# Impact of nitrogen and phosphorus addition on resident soil and root mycobiomes in beech forests

**DOI:** 10.1101/2020.12.29.424645

**Authors:** S. Clausing, L.E. Likulunga, D. Janz, H.Y. Feng, D. Schneider, R. Daniel, J. Krüger, F. Lang, A. Polle

## Abstract

In forest soils, the pools of N and P available for microbes and plants are strongly dependent on soil properties. Here, we conducted a P and N fertilization experiment to disentangle the effects of nutrient availability on soil-residing, root-associated and ectomycorrhizal fungi in beech (*Fagus sylvativa*) forests differing in P availability. We tested the hypothesis that in P-poor forests, P fertilization leads to enhanced fungal diversity in soil and roots, resulting in enhanced P nutrition of beech and that N fertilization aggravates P shortage, shifting the fungal communities towards nitrophilic species. In response to fertilizer treatments (1x 50 kg ha^−1^ P, 5x 30 kg ha^−1^ N within 2 years), the labile P fractions increased in soil and roots, regardless of plant-available P in soil. Root total P decreased in response to N fertilization and root total P increased at the low P site in response to P addition. The relative abundances of ectomycorrhizal fungi, but not their species richness, increased in response to P or N addition in comparison with that of saprotrophic fungi. While some fungal orders (Trechisporales, Atheliales, Cantharellales) were moderately decreased in response to fertilizer treatments, Boletales increased in response to P and Russulaes to N addition. N or P fertilization resulted in functional trade-off, shifting away from saprotrophic towards symbiotrophic potential. Our results suggest that chronic exposure of forest ecosystems to increased nutrient inputs may overcome the resistance of the resident mycobiome structures resulting in nutritional imbalance and loss of forest ecosystem services.

## 1 Introduction

Nitrogen (N) and phosphorus (P) are essential nutrients that determine growth and productivity in many terrestrial ecosystems (Elser et al. 2007; Vitousek et al. 2010). Natural sources of N and P are the biological N fixation from atmospheric di-nitrogen and rock weathering for P (Augusto et al. 2017; Wardle 2004). Therefore, the P availability mainly depends on soil parent material and soil age (Augusto et al. 2017; Wardle, 2004), while N can also be influenced by anthropogenic inputs, for example, by atmospheric N deposition from agriculture and burning of fossil fuels (Galloway et al. 2008). Since forests are usually not fertilized, tree growth is limited by the least available resource. Consequently, alterations in the input and demand of P and N can change the nutrient regime of ecosystems (Vitousek et al., 2010). Unbalanced N depositions into forest ecosystems have increased the primary production in N-limited forest ecosystems (Schulte-Uebbing and De Vries 2018; Du and De Vries 2018). Increasing N:P ratios have therefore been suggested to indicate that many European forest ecosystems are transitioning from N to P limitation (Peñuelas et al. 2013; Jonard et al. 2015).

In temperate and boreal forest soils, large fractions of P and N are bound by organic matter and, thus, not directly available for uptake by trees (Lambers et al. 2008; van der Heijden et al. 2008). Trees benefit from P and N mineralization by microbial decomposers (Schimel and Schaeffer 2012, Baldrian 2017). Soil fungi are generally more efficient regarding the degradation of complex plant compounds than other soil microbiota (Stursová et al. 2012; López-Mondéjar et al. 2018). Thus, the taxonomic diversity and functional composition of soil fungal microbiomes is of high relevance for forest P and N nutrition. Both saprotrophic and ectomycorrhizal fungi (EMF) contribute to N and P mobilization by secretion of organic acids, and the production of hydrolytic and oxidative exoenzymes (Courty et al. 2010, Pritsch and Garbaye 2011, Bödeker et al. 2014, Op De Beeck et al. 2018). In deciduous temperate forest soils, the fraction of EMF is similar to or even higher than that of saprotrophs confirming that EMF have eminent functions for nutrient recycling in these ecosystems (Awad et al. 2019). However, the impact of changes in N and P availability on these functional groups and P nutrition of trees is still unknown.

Among the multiple abiotic and biotic habitat filters such as climate, geographic location, soil type, vegetation, etc. that drive soil fungal community structures (Wubet et al. 2012, Suz et al. 2014, Tedersoo et al. 2014, Bahr et al. 2015, Goldmann et al. 2016, Kolaříková et al. 2017, Bahnmann et al. 2018, van der Linde et al. 2018), N is an important factor (Cox et al. 2010, de Witte et al. 2017, Schröter et al. 2018, van der Linde et al. 2018, Almeida et al. 2019, Nguyen et al. 2020). For example, Lilleskow et al. (2002) reported a shift in the EMF community towards nitrophilic species and, thus, a loss in diversity along a gradient of increasing N deposition in Alaska (Lilleskov et al. 2002). In boreal spruce forests, N fertilization caused a significant turnover of soil fungal communities, decreased fungal biomass, and increased the N:P ratio of the needles (Almeida et al. 2019). Other studies reported only weak or no effect of N treatments on the fungal community structures (Nicolás et al. 2017, Purahong et al. 2018, Maaroufi et al. 2019) and relationships with P mobilization were not detected (Forsmark et al. 2020). Only few studies investigated fungal communities after P fertilization in forests. Almeida et al. (2019) reported significant community turnover and loss in fungal biomass after P fertilization in a spruce forest. However, along a natural P gradient in temperate beech forests, EMF diversity increased with increasing P availability (Zavišić et al. 2016). After addition of superphosphate to P-limited young beech trees, the EMF community structures were altered but microbial biomass was unaffected (Zavišić et al. 2018). These disparate observations indicate that the responses of EMF and other fungi to P inputs depend on the P supply by the soil and probably also the interaction of P and N supply (de Witte et al. 2017). Furthermore, different avalilabilities of carbon for fungal communities colonizing the root surface and those living in soil (Clausing et al. 2020) may influence the responses of fungi to N and P availabilities. However, experiments that enhance our understanding of these ecological processes are lacking.

The aim of our study was to investigate how shifts in N and P availabilities affect fungal community structures and functional composition in belowground habitats with different reliance on root resources: functional ectomycorrhizas with root tips (EMF), root-associated fungal assemblages (RAF) and soil-associated fungi (SAF). Furthermore, we studied the impact of P and N soil treatments on root P and N contents. To address these goals, we established plots fertilized with either P, N or P+N along a geosequence in beech (*Fagus sylvatica* L.) forests, where P nutrition shifts from P acquiring (high P) to P recycling (low P) conditions (Lang et al. 2016, Lang et al. 2017). We used these plots to determine soil and root P and N contents and characterized the mycobiomes of EMF, RAF and SAF. Since tree nutrition depends mainly on P from the organic layer under low P availability, while P extracted by EMF from deeper layers (mineral soil) also contributes significantly to plant P supply in P-rich soils (Clausing and Polle 2020), we investigated the vertical stratification of the fungal assemblages. Specifically, we tested the following hypotheses: (i) N fertilization aggravates root P shortage in P-poor soil and has no effects on root P contents and fungal community structures in P-rich soils. (ii) P fertilization increases fungal richness and diversity in P-poor soils because fungal communities adapted to P shortage are composed of specialized fungi, while P-rich soils foster diverse fungal communities. (iii) N fertilization reduces fungal diversity with stronger effects on the EMF than on the saprotrophic fungal taxa because plant roots become less reliant on ectomycorrhizal nutrient supply than under low nutrient availability.

## 2 Material and Methods

### 2.1 Site characteristics and study plots

The N and P fertilization experiments were carried out in three beech (*F. sylvatica* L.) forests, differing in parent material and thus, soil P stocks. The high-P site (HP) Bad Brückenau is located in the biosphere reservation ‘Bayerische Rhön’ on basalt, the medium-P site (MP) Mitterfels is situated in the Bavarian Forest on paragneiss and the low-P site (LP) Luess is located in the North German Plain on sandy till. P stocks in the A horizon (1 m soil depth) at the HP, MP and LP site are approximately 9.0, 6.8 and 1.3 t ha^−1^, respectivlely (Supplement Table S1, Lang et al., 2017). The pH of the soils ranged from 3.5 to 3.8 (Supplement Table S1). Information on the climate (1981 to 2010) and weather during sampling was obtained from www.wetterzentrale.de (Supplement Table S2).

For this study, twelve plots with an area of 400 m^2^ each were installed in each forest in summer 2016 under 120- to 140-year-old beech trees (Supplement Table S1). One control (Con) and three different fertilizer treatments (N, P, P+N), each replicated three times per forest (= a total of 36 plots), were treated as follows: phosphorus (P) was applied once in late summer 2016 as KH_2_PO_4_ at the dosage of 50 kg P ha^−1^ to the P and N+P plots. Nitrogen (N) was applied as NH_4_NO_3_ five times (late summer 2016, spring 2017, summer 2017, fall 2017, spring 2018) corresponding to a dosage of 30 kg N ha^−1^ per treatment on the N and N+P plots. To account for the K input in the P treatments, KCl was applied once in fall 2016 to the Con and N plots. The minerals were dissolved in tap water and applied with garden sprayers.

### 2.2 Harvest and processing of soil cores

Soil was sampled in the third year after the start of the treatments in spring (LP: 16.04.2018, HP: 23.04.2018, MP: 02.05.2018) and fall (LP: 17.09.2018, HP: 25.09.2018, MP: 01.10.2018). The weather conditions in the month of sampling and before as well as the long-term climate (1981 to 2010) are shown in the supplementary materials (Supplement Table S2). The sampling took place before the application of N, meaning that approximately 6 months passed after N addition and before soil sampling. In each plot, twelve randomly distributed soil cores (depth 0.21 m, diameter 55 mm) were extracted after removal of surface litter. Each soil core was separated in organic (OL) and mineral topsoil (ML). The respective layers were pooled yielding one OL and one ML sample per plot. Each sample was fractionated into bulk soil, fine roots (< 2 mm), coarse roots (> 2 mm) and residual materials (fruits, twigs, and stones) in the field. Each sub-sample was directly divided into three aliquots: a fresh sample that was kept cool at 4 °C until use, a sample that was immediately frozen in liquid nitrogen (and stored at −80 °C in the laboratory) and a sample that was dried (40 °C, 14 days). Bulk soil was sieved (mesh width: 4 mm) and the root samples were carefully washed before the aliquots were taken. All fractions were weighed in the laboratory. During the harvest in fall 2018, an additional soil core was collected in each plot. The sampling position was located adjacent to that of the soil cores for chemical analysis. The extra soil core was used to collect the roots for mycorrhizal morphotyping.

### 2.3 Determination of soil and root chemistry

#### Inorganic N

To determine the concentrations of ammonium (NH_4_^+^) and nitrate (NO_3_^−^) in soil, 20 g of fresh sieved bulk soil was extracted at the field site in 40 ml CaCl_2_ solution for 60 min under shaking, subsequently filtered with phosphate free filter paper (MN 280 ¼, Macherey-Nagel, Düren, Germany) and kept cool. In the laboratory, the extracts were dried twice by cryodessication for 72 h (BETA I, Christ, Osterode am Harz, Germany) and dissolved in 1.5 ml ultra-pure water. The concentrated extracts were used to determine nitrate (# 109713, Merck, Darmstadt, Germany) and ammonium concentrations with kits (# 100683, Merck) according to the manufacturer’s instructions. The extinction of the assays was measured in an UV-Vis spectrophotometer (Shimadzu 1601, Hannover, Germany) at 690 nm for NH_4_^+^, and 340 nm for NO_3_^−^.

#### Soil pH

To determine the pH values, 10 g of oven dried milled soil was suspended in 25 ml deionized water and shaken for 1 hour at 200 rpm. After sedimentation of the particles, the pH was measured by a pH meter (WTW, Weilheim, Germany). After addition of 0.01 M CaCl_2_ (1:5 soil-to-solution ratio) and 16 h equilibration, the samples were measured again (ISO10390, 2005).

#### Element content

Dry soil and root samples were milled (Retsch MN 400, Haan, Germany) to a fine powder. For the determination of total P (P_tot_), 50 mg of the powder was weighed and extracted in 25 ml of 65% HNO_3_ at 160 °C for 12 h according to Heinrichs et al. (1986). For the determination of the labile P (P_lab_) fraction about 100 mg of soil or root powder was extracted in 150 ml of Bray-1 solution (0.03 N NH_4_F, 0.025 N HCl) for 60 min on a shaker at 180 rpm (Bray and Kurtz 1945). The extracts were filtered using phosphate free filter paper (MN 280 ¼, Macherey-Nagel) and used for elemental analysis by inductively coupled plasma–optical emission spectroscopy (ICP–OES) (iCAP 7000 Series ICP-OES, Thermo Fisher Scientific, Dreieich, Germany). P was measured at the wavelength of 185.942 nm (axial) and calibrated with a series of concentrations by element standards (P: 0.1 mg l^−1^ to 20 mg l^−1^) (Einzelstandards, Bernd Kraft, Duisburg, Germany). In addition to P, we also determined K, Ca, Mg, Mn, Fe, Al, and S in the P_tot_ extracts.

#### Carbon and nitrogen

Subsamples of 2 to 12 mg of soil or 1.5 mg of root powder were weighed into tin capsules (size of 4 × 6 mm, IVA Analysentechnik, Meerbusch, Germany) using a microbalance (Model MC5, Sartorius, Goettingen, Germany). The range of weights for the soil samples was necessary to avoid overflow of the measuring unit of the mass spectrometer since the C concentrations in the OL and ML and between sand of other soil types varied drastically. The amounts of C and N of the soil and plant samples were measured at the KOSI (Kompetenzzentrum Stabile Isotope, Göttingen. Germany) using an isotope mass spectrometer (Delta Plus, Finnigan MAT, Bremen, Germany) and acetanilide (10.36% N, 71.09% C) as the standard.

### 2.4 Determination of ectomycorrhizal fungal species abundances by morphotyping and Sanger sequencing

Roots from the extra soil core collected in fall were separated according to organic layer and the mineral topsoil and immediately processed after sampling. The beech roots were gently washed in 4 °C precooled tap water, spread in water in a glass dish, and categorized according to their visual appearance under a stereomicroscope (Leica M205 FA, Wetzlar, Germany) as vital ectomycorrhizal, vital non-mycorrhizal or dry (Winkler et al. 2010). All root tips in each soil core were inspected and counted. Two soil cores did not contain any roots. Ectomycorrhizal colonization and root tip mortality were calculated as:

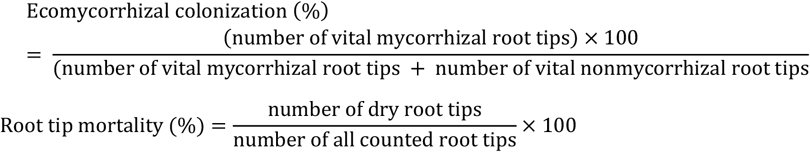

The abundance of different morphotypes was determined under a stereomicroscope (Leica M205 FA, Wetzlar, Germany) using a simplified identification key (after Agerer, 1987-2012) as described by Pena et al. (2010). All root tips in each soil core were categorized and counted. For mycorrhizal species identification, all different morphotypes were collected, which comprised at least three root tips per sample. Samples with no ectomycorrhizal root tips were included as zero values. We distinguished 44 morphotypes and sequenced 19 morphotypes, which were most abundant and covered >90% of the beech root ectomycorrhizal fungal community. We used the protocol of Pena et al. (2017) for DNA extraction, ITS sequencing and species identification. Fungal sequences have been deposited in NCBI GenBank under the accession numbers MT859114 to MT859131 (Supplement Table S3).

### 2.5 DNA extraction and preparation of soil and root samples for Ilumina sequencing

Frozen soil and root samples that had been stored at −80 °C were milled in a ball mill (Retsch GmbH, Haan, Germany) in liquid nitrogen. DNA was isolated from 250 mg soil or from 180 mg roots with the DNeasy^®^ PowerSoil^®^ Pro kit (Qiagen, Hilden, Germany) or innuPREP Plant DNA kit (Analytik Jena AG, Jena, Germany), following the manufacturer’s recommendations. DNA was purified using the DNeasy^®^ PowerClean^®^ Pro Cleanup kit (Qiagen). The amount of isolated DNA was measured in a NanoDrop ND-1000 spectrophotometer (PEQLAB Biotechnologie GmbH, Erlangen, Germany). For each DNA extraction, a PCR was performed in a reaction volume of 50 μl using 0.3 μl of Phusion High-Fidelity DNA Polymerase (2 U μl^−1^, New England Biolabs (NEB), Frankfurt, Germany), 6 μl of 5x Phusion HF buffer (NEB), 0.09 μl of MgCl_2_ (50 mM, NEB), 0.6 μl of dNTP mix (10 mM each, Thermo Fisher Scientific, Osterode am Harz, Germany), 0.6 μl of the forward (ITS3-KYO2) and reverse primer (ITS4) (10 mmol/l, Microsynth, Wolfurt, Austria) and about 250 (roots) to 1050 (soil) ng of template DNA in 5 μl. The primers ITS3-KYO2 (Toju et al. 2012) and ITS4 (White et al. 1990) included the adapters for MiSeq sequencing. The PCR reactions were performed in a Labcycler (SensoQuest, Göttingen, Germany). The cycling parameters were 1 cycle of 98 °C for 30 s; 30 cycles of 98 °C for 10 s, 47 °C for 20 s and 72 °C for 20 s; and a final extension at 72 °C for 5 min. The PCR products were subjected to electrophoresis in 2% agarose gels (Biozym LE Agarose, Biozym Scientific GmbH, Hessisch Oldendorf, Germany) using GelRed (10 000×, VWR, Darmstadt, Germany) to stain the 1 kb DNA ladder (NEB) that was used for the determination of the product size. The PCR products were visualized with an FLA-5100 Fluorescence Laser Scanner (Raytest GmbH, Straubenhardt, Germany) and an Aida Image Analyser v. 4.27 (Raytest GmbH). All PCR reactions were performed in triplicate, pooled, and purified using the MagSi-NGS^PREP^ Plus Kit (Steinbrenner Laborsysteme, Wiesenbach, Germany). Quantification of the purified PCR products was performed with a Quant-iT dsDNA HS assay kit (Life Technologies GmbH, Darmstadt, Germany) in a Qubit fluorometer (Life Technologies GmbH, Darmstadt, Germany) following the manufacturer’s recommendations.

### 2.6 Amplicon sequencing and bioinformatic processing

Amplicon sequencing was conducted at Göttingen Genomics Laboratory on the MiSeq platform using the MiSeq Reagent Kit v3 (Illumina Inc., San Diego, USA). For amplicon sequence variant (ASV) assembly paired-end sequencing data from the Illumina MiSeq were quality-filtered with fastp (version 0.20.0) using default settings with the addition of an increased per base phred score of 20, base pair corrections by overlap (-c), as well as 5’- and 3’-end read trimming with a sliding window of 4, a mean quality of 20 and minimum sequence size of 50 bp (von Hoyningen-Huene et al. 2019). Subsequently, quality-filtered reads were merged using PEAR v.0.9.11 (Zhang et al. 2014) with default parameters. Primer sequences were clipped with cutadapt v.2.5 (Martin 2011). VSEARCH v.2.14.1 (Rognes et al. 2016) was used for size exclusion of reads <140 bp, dereplication, denoising (UNOISE3, default settings) and chimera removal (de novo followed by reference-based chimera removal). ASVs were clustered at 97 % sequence identity [corresponding to operational taxonomic units (OTUs), the usual threshold in most fungal studies] employing VSEARCH (--sortbysize and --cluster_size). Reads were mapped to OTUs and used to create a count table using VSEARCH (--usearch_global, -id 0.97).

OTUs were taxonomically classified using the BLAST algorithm against the UNITE+INSDC 8.2 public database (Kõljalg et al. 2013) with an identity cutoff of 90%. Unclassified and non-blast hit OTUs (<90% identity) were aligned against the GenBank (nt, 2020-01-17) database (Geer et al. 2010) and only OTUs with a fungal classification were kept in the OTU table. The OTU count table was rarefied to the count number of 11,000 (minimum number reads in one sample) using the rrarefy() function of the package vegan v2.5.6 (Oksanen et al. 2019). OTUs were functionally annotated as symbiotroph, pathotroph and saprotroph using the FUNGuild database (Nguyen et al. 2016).

### 2.7 Statistical procedures and calculations

The statistical analyses were performed with R version 3.6.0 (R Core Team 2012). Normal distribution and homogeneity of variances were tested by analyzing the residuals of the models and performing a Shapiro-Wilk test for chemical soil and root parameters and EMF abundance. Data were logarithmically or square root-transformed to meet the criteria of normal distribution and homogeneity of variances, where necessary. Count data of fungal orders and trophic groups were not transformed. Visual inspection of their residuals showed homogenous distribution and therefore, these data were also analyzed by linear mixed effect models using Poisson distribution (O’Hara and Kotze 2010).

To determine the effects of forest type, soil layer, season, habitat, and treatment linear mixed effect models (‘lmer’, R package lme4) were used. The factor plot was used as random effect and the factor season (spring and fall) was defined as repeated measurement in the model. Pairwise comparisons of the sample means were conducted using Tukey’s HSD (package: ‘multcomp’). Means were considered to be significantly different from each other when p ≤ 0.05. Data are shown as means and standard error (±SE) of the three plots per treatment, if not indicated otherwise.

We used the following fungal communities for our analyses: EMF, SAF, and RAF. RAF and SAF were distinguished in the Illumina dataset and further discerned as SYM, SAP and PAT corresponding to the groups of ectomycorrhizal (SYM), saprotrophic (SAP) and pathogenic fungi (PAT). To obtain ectomycorrhizal fungi, the guild of symbiotrophic fungi was manually screened and other symbiotrophic fungi (arbuscular mycorrhizal, orchid mycorrhizal, endophyte, lichenized, combinations of saprotrophic or pathotrophic with EMF) were eliminated and then used as SYM for further analyses.

Non-metric multidimensional scaling (NMDS) was used to explore and visualize the main factors affecting fungal community composition using Bray Curtis as the dissimilarity measure. We tested multicollinearity by Pearson’s pairwise correlation for nutrient elements (P_tot_, P_lab_, N, C, Na, K, Ca, Mg, Mn, Fe, Al, S) and RWC (relative water content) in bulk soil and roots using the ‘hmisc’ package (Harrell Jr 2016). We tested variables that were unrelated with the exception of Ca, P, and N as explanatory variables for the NMDS and showed the significant vectors (p < 0.05). The function ANOSIM (analysis of similarity) (package: ‘vegan’) (Oksanen et al., 2019) was used to analyze the dissimilarities among the fungal communities for the factors: fores type, habitat, layer, season and treatment.

## 3 Results

### 3.1 Influence of P and N addition on soil and root chemistry

P was applied only once in fall 2016 and nitrogen four respective five times. Therefore, our soil samples collected in spring 2018 were exposed to 120 kg N ha^−1^ and those in fall 2018 to 150 kg N ha^−1^, whereas the P dose was 50 kg ha^−1^ for all samples. Most of the parameters analyzed in soil and roots varied with forest site and season, while treatment effects were less abundant and mainly found for P (Table 1).

**Table 1:**
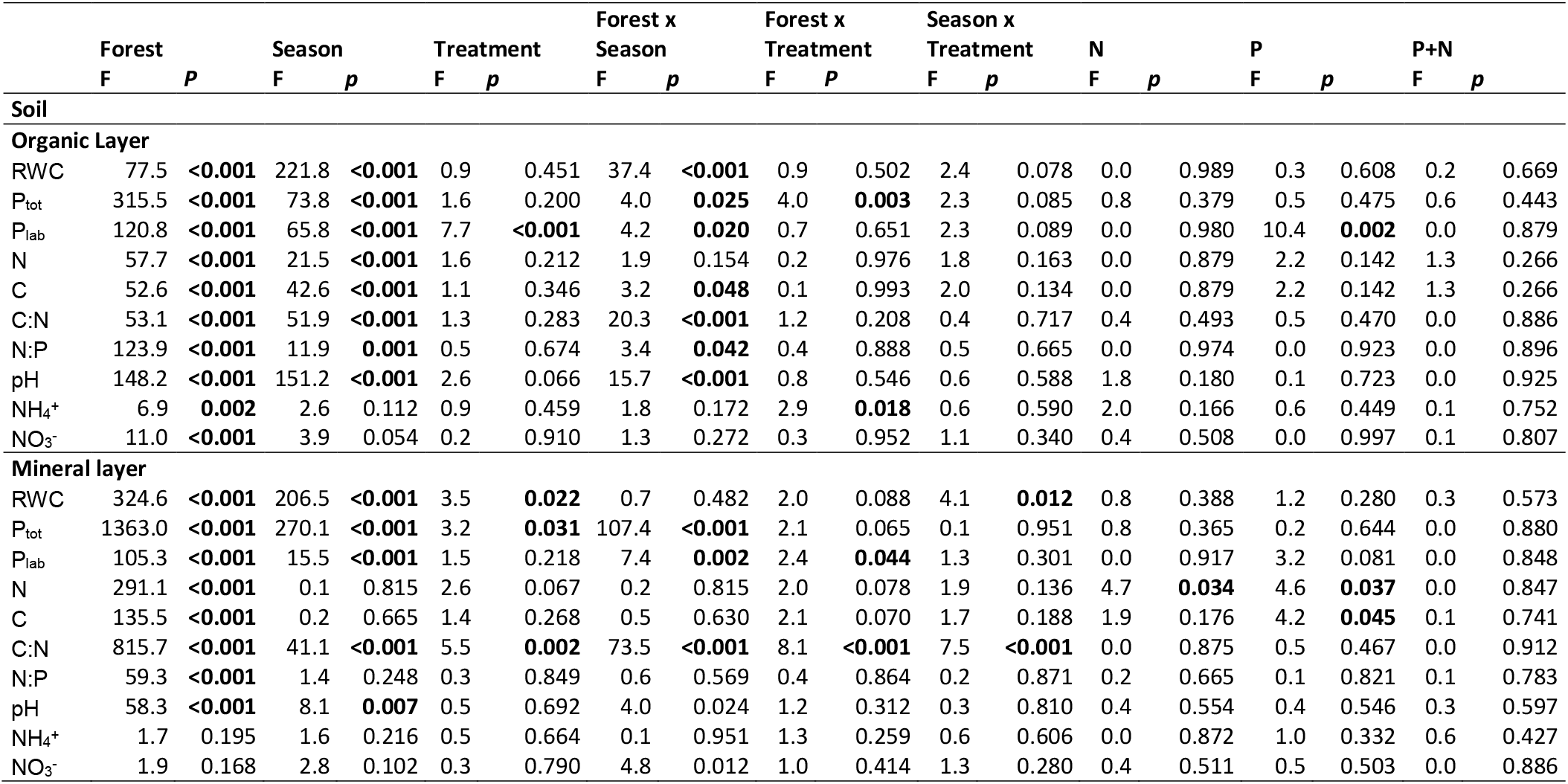
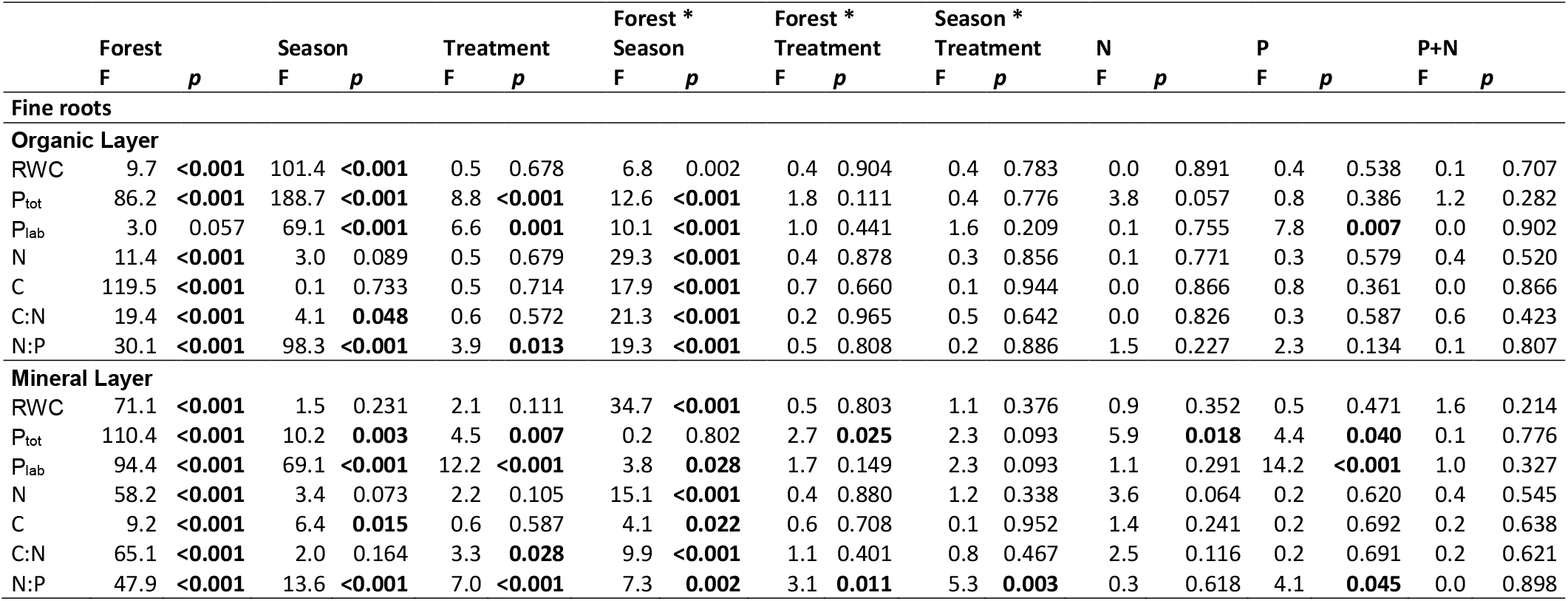
Statistical information on the effects of forest site, season and fertilization treatments on soil and root chemistry in the organic layer and in the mineral topsoil. Soils and roots were collected in a P-rich (HP), P-medium (MP) and P-poor (LP) forest in spring and fall 2018 and separated into organic layer and the mineral topsoil for analyses. Means can be found in the supplementary information (Supplement Table S4). Differences among means per forest type, season, treatment and the two-way interactions (Forest x season, Forest x treatment, season x treatment) were tested by a linear mixed effect model with plot number as random effect. Calculations were performed separatley for the organic layer and mineral topsoil. Tukey HSD was used as posthoc test. Bold letters indicate significant differences at p ≤ 0.05. RCW = relative water content.

P addition resulted in increased soil P_lab_ concentrations across the three study forests but did not affect the soil P_tot_ concentrations (Fig. 1 a-c, Table 1). The effects were also present when the forests were fertilized with P+N (Fig. 1a-c) and were more pronounced in fall than in spring (Table 1, Supplement Table S4). Furthermore, the P fertilization effects were stronger in the organic layer than in the mineral soil (Fig. 1 a-c, Table 1, HP: F = 12.6, *p* = 0.001; LP: F = 111.1, *p* < 0.001), with the exception of the MP forest soils (F = 0.02, *p* = 0.902, Fig. 1 b).

**Figure 1:**
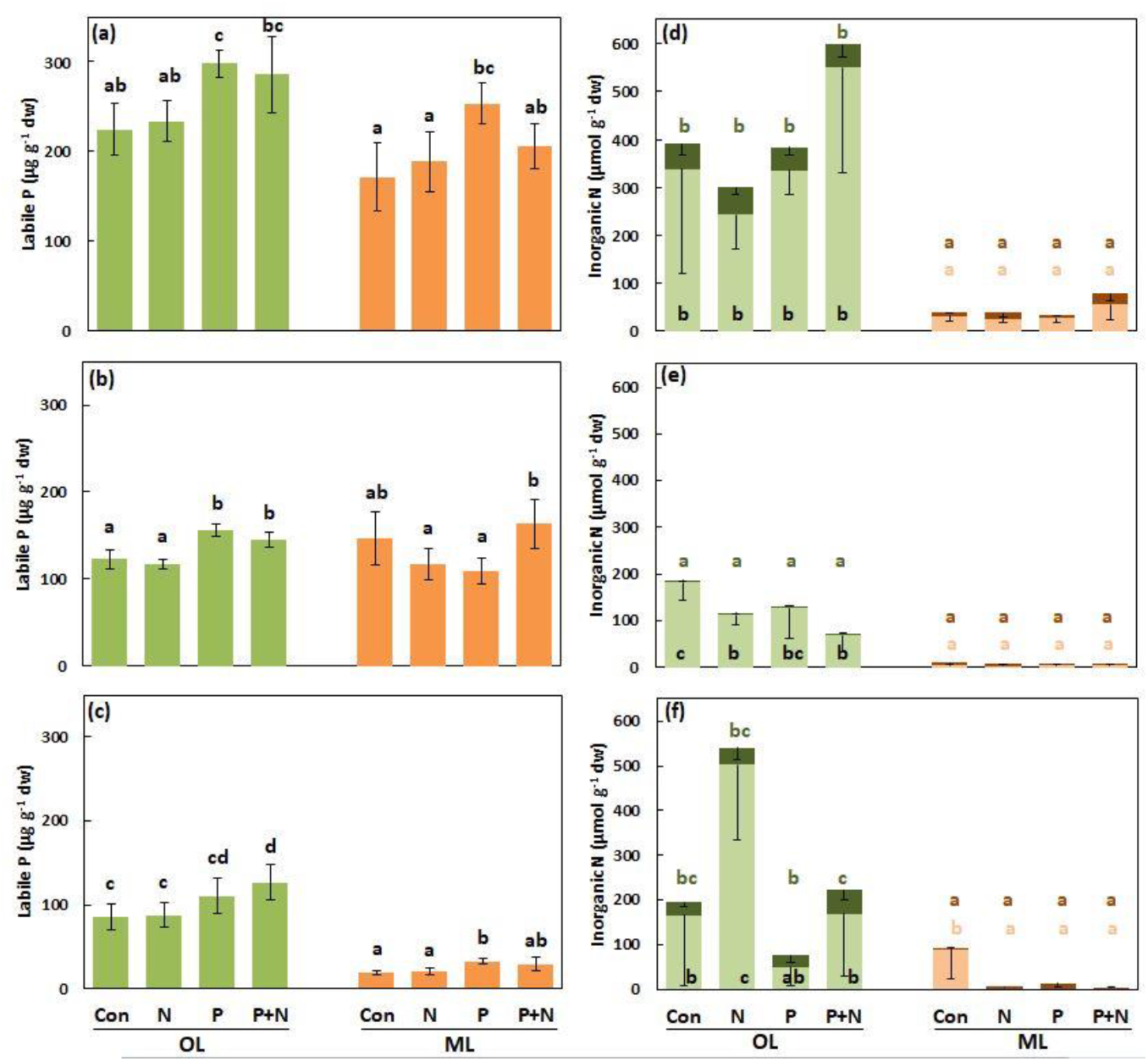
Labile phosphorus (a-c) and inorganic nitrogen (d-f) concentrations the soil of beech forests (*Fagus sylvatica* L.). Soils were collected in a P-rich (a, d), P-medium (b, e) and P-poor (c, f) forest and separated into organic layer (green) and the mineral topsoil (orange) for analyses. Stacked bars show ammonium (bright colors) and nitrate (dark colors). Data for spring and fall were pooled. Data indicate means (n = 6 ± SE). Differences among means were tested by a linear mixed effect model with plot number as random effect and Tukey HSD posthoc test. Different letters indicate significant differences at *p* ≤ 0.05.

The higher availabilities of P_lab_ in P-fertilized forest soils resulted in higher fine root P_tot_ concentrations in the LP forest but not in the HP and MP forests (Fig. 2a-c). In all three forests, the P_lab_ concentrations of the roots were higher after P fertilization (Fig. 2 d-f). Across the three forests fine root P_lab_ was higher in spring (556 ± 29 μg g^−1^ dw) than in fall (320 ± 19 μg g^−1^ dw, F = 25.3, *p* <0.001 and higher in the organic layer (526 ± 31 μg g^−1^ dw) than in the mineral layer (351 ± 20 μg g^−1^ dw, F = 22.0, *p* <0.001) (Supplement Table S4).

**Figure 2:**
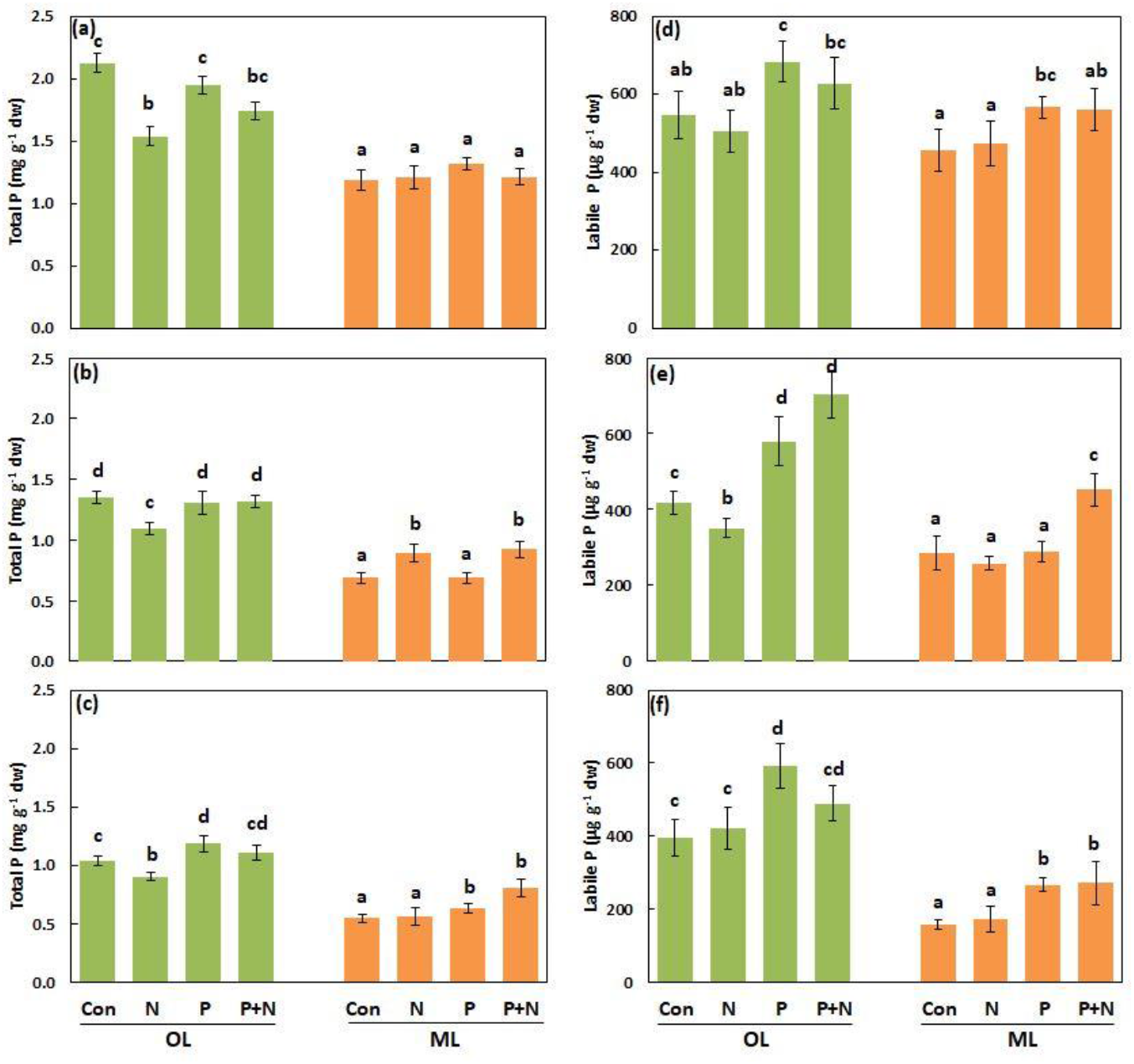
Total phosphorus (a-c) and labile phosphorus (d-f) in roots of beech forests (*Fagus sylvatica* L.). Fine roots were collected in a P-rich (a, d), P-medium (b, e) and P-poor (c, f) forest and separated into organic layer (green) and the mineral topsoil (orange) for analyses. Season data were pooled. Data indicate means (n = 6 ± SE). Differences among means were tested by a linear mixed effect model with plot number as random effect and Tukey HSD posthoc test. Different letters indicate significant differences at *p* ≤ 0.05.

N fertilization caused small alterations in the C/N ratio towards slightly lower values at MP and higher values at LP (Table 1, Supplement Table S4). The N addition did not result in increases in the soluble, inorganic N concentrations (NO_3_^−^, NH_4_^+^) in the mineral soil (Fig. 1 d-f). In the organic layer, the N-fertilized MP forest contained lower NH_4_^+^ concentrations than the controls (Fig. 1 e), while no effect was found in the HP and an increase in the LP forest (Fig. 1 d,f). Significant seasonal effects of the treatments on the NO_3_^−^ or NH_4_^+^ concentrations were not observed (Table 1). Since N turnover is rapid and great variations of NO_3_^−^ and NH_4_^+^ in soil solution are known (Ollivier et al. 2011, Cheng et al. 2019), our analyses of NO_3_^−^ and NH_4_^+^ represent snapshots during our sampling campains.

N fertilization caused significant decreases in P_tot_ concentrations in the roots from the organic layer (Fig. 2a-c) but did not result in changes in root N concentrations (Table 1, Supplement Table S4). Therefore, the treatments also caused increases in the N:P ratios of the roots but not in soil (Table 1, Supplement Table S4).

### 3.2 P addition has a moderate effect on fungal diversity in the SAF at the high-P forest

We analyzed fungal species richness at three scales EMF, RAF, and SAF (Table 2, Supplement Table S5). For the morphotyping-Sanger sequencing approach of EMF, all root tips per soil core were counted, resulting a total of 44 morphotypes. The most abundant morphotypes (n = 19), which colonized a total >90% of the mycorrhizal root tips, were sequenced (Supplement Fig. S1, Supplement Table S3). We detected a mean of 8.0 ± 0.4 EMF species per treatment and forest (Supplement Table S5). EMF richness in the organic layer varied among the forests but not in response to the treatments (Table 2, Supplement Table S5). Diversity and evenness were also unaffected by the treatments (Table 2, Supplement Table S5). Fertilization did not affect the fraction of mycorrhizal root tips (99.9 ± 0.4%) nor the fraction of dead root tips (29.7 ± 1.6%) (Supplement Table S6). However, in the mineral layer at the MP and LP forests root mortality was higher than at the HP forest (Supplement Table S6).

**Table 2:**
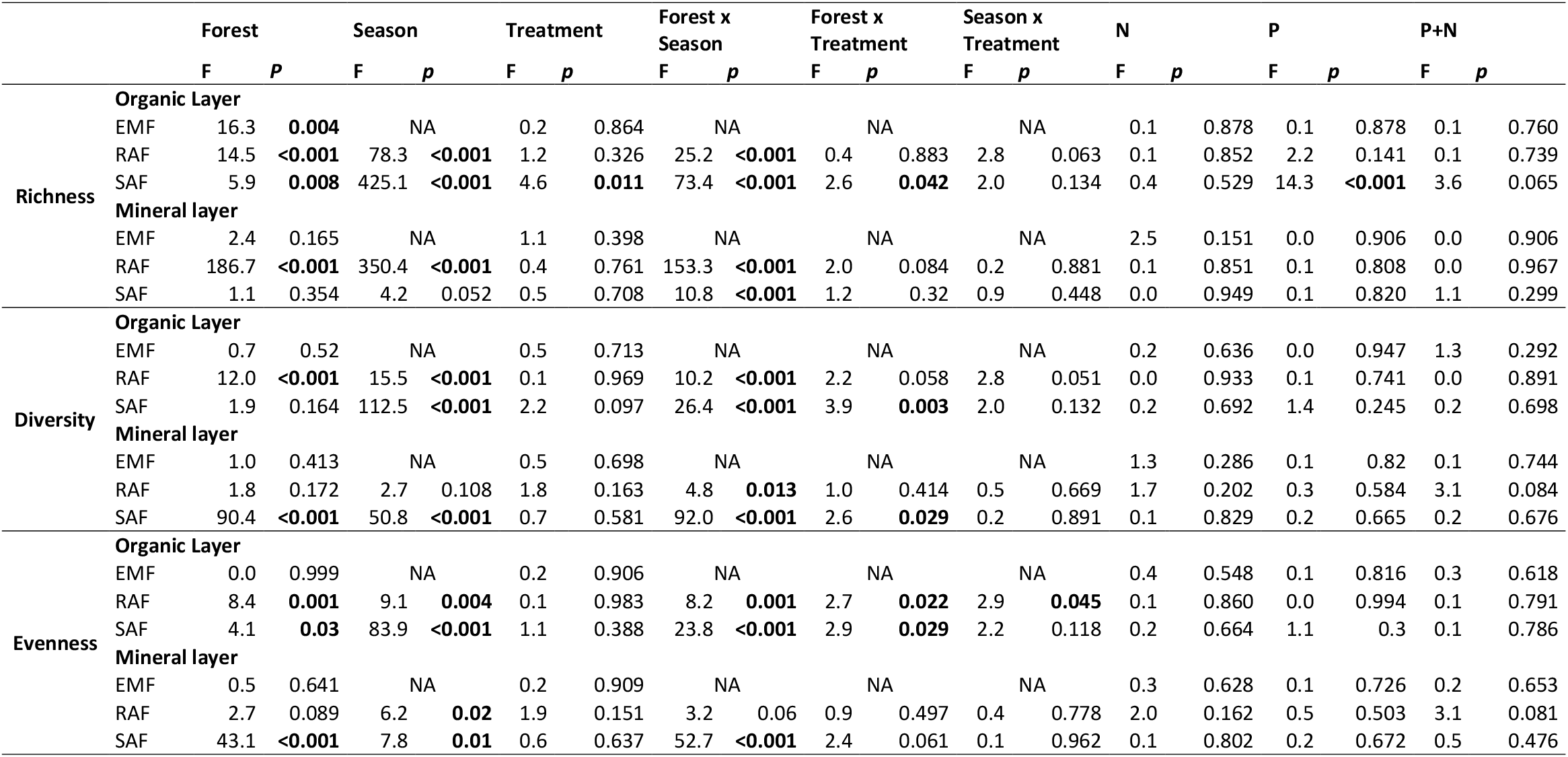
Statistical information on the effects of forest site, season and fertilization treatments on diversity parameters of EMF, RAF and SAF in the organic layer and in the mineral topsoil. Soils and roots were collected in a P-rich, P-medium and P-poor forest in spring and fall 2018 and separated into organic layer and the mineral topsoil for analyses. Means can be found in the supplementary information table (Table S5). Differences among means per forest type, season, treatment and the two-way interactions (Forest x season, Forest x treatment, season x treatment) were tested by a linear mixed effect model with plot number as random effect. Calculations were performed separately for the organic layer and mineral topsoil. Tukey HSD was used as posthoc test. Bold letters indicate significant differences at *p* ≤ 0.05. NA = not available

Using Illumina sequencing (288 samples in total), we obtained 3,169 million fungal sequence reads, which clustered into 4,134 different OTUs. Across all conditions, RAF communities contained approximately three times less OTUs (219 ± 12) than the SAF communities (764 ± 10, F = 16155, *p* < 0.001). Most of the diversity indices of the RAF and SAF (richness, Shannon diversity and evenness) showed significant variation for the comparisons among forests and between seasons (Table 2, Supplement Table S5). Fungal OTU richness of the RAF and the SAF was higher in the organic than in the mineral layer and higher in fall than in spring (Supplement Table S5). However, no clear pattern with increasing P availability in soil was found (Table 2, Supplement Table S5).

We did not observe any effects of P or N additions on the OTU richness or diversity in the RAF communities (Table 2) but the SAF exhibited a treatment effect in the organic layer (Table 2) because in fall the P-fertilized plots at LP contained higher OTU richness than the unfertilized plots (Supplement Table S5). Differential abundance measurements (using for instance DESeq2 according to Love et al., 2014) to identify which fungal OTUs were increased were not successful because the majority of OTUs were represented by low and variable numbers of sequence reads that spread across 92 fungal orders (Supplement Table S7).

### 3.3 Fungal community structures are governed by soil and root chemistry

The dissimilarities of the fungal community structures of EMF, RAF and SAF were visualized by NMDS (Fig. 3). NMDS and ANOSIM showed that the EMF community structures differed significantly among the forests and between the organic and mineral soil layers but not in response to the fertilization treatments (Fig. 3 a). Similarly, the composition of the RAF and SAF communities were unaffected by treatments, whereas forest, soil layer and season caused significant differentiation (Fig. 3 b, c). The fungal community structures also differed significantly between the soil and the root habitat (F = 0.479, p <0.001). The main factors that explained the dissimilarities of the SAF were relative water content, P_tot_, Ca, and N of both soil and roots (Fig. 3, Table 3). The relative water content of the soil was less important for the variation in the dissimilarities of the RAF and EMF structures than for the SAF (Table 3). In contrast to the SAF, soil N was no explanatory variable for the RAF and neither soil N nor root N for the EMF. Nutrients that were influenced by the treatments in soil such as P_lab_ (Fig. 1) were correlated with P_tot_ and excluded due to multicollinearity. The fungal community structures were mainly explained by gross differences due to soil contents of P and Ca (Table 3).

**Figure 3:**
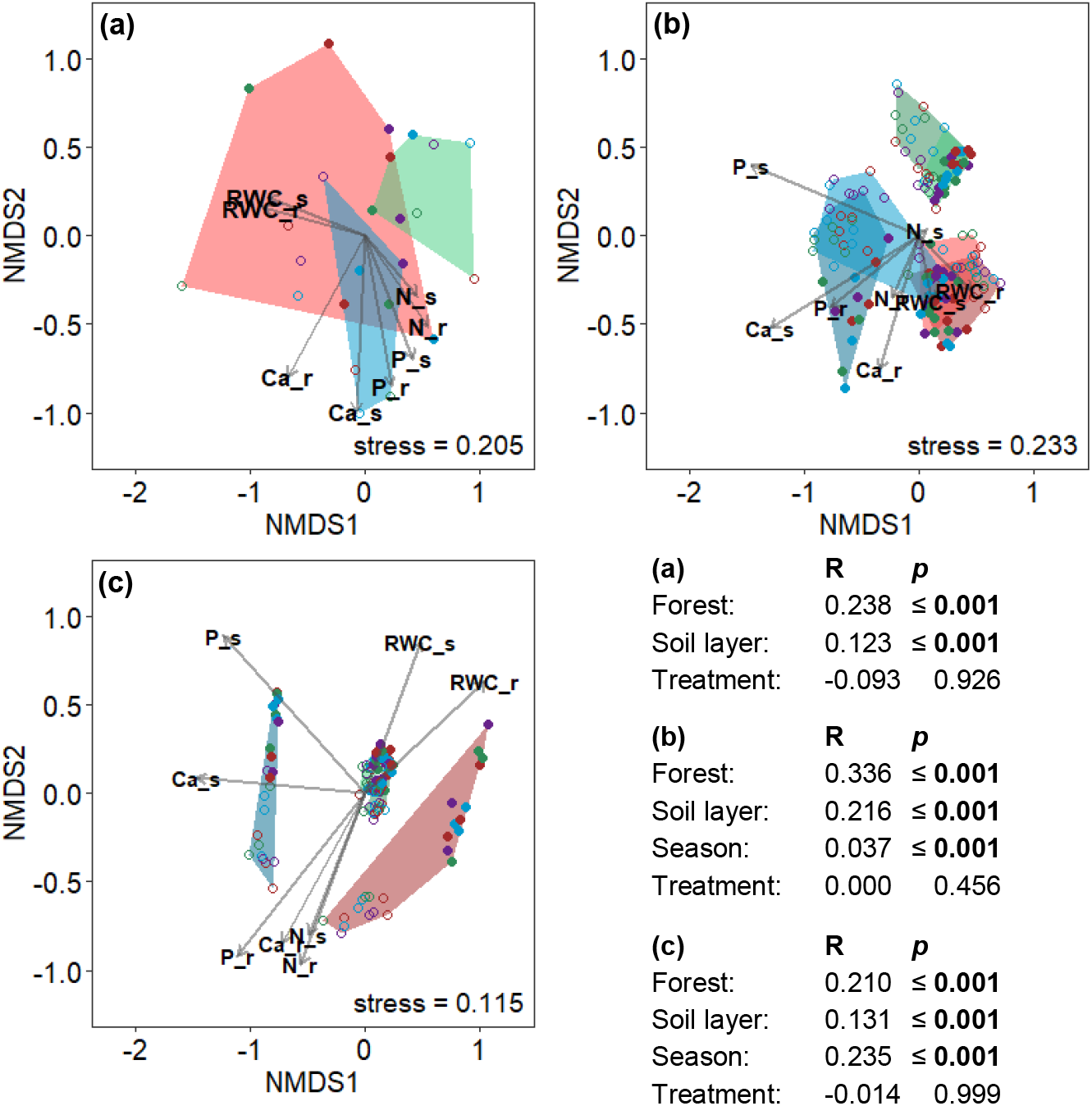
Non-metric multidimensional scaling (NMDS) for ectomycorrhizal (a), root (b) and soil fungi (c) of beech forests (*Fagus sylvatica* L.). Soil and fine root samples were collected in a P-rich (blue), P-medium (green) and P-poor (red) forest in 2018. The figure shows samples from the organic layer (open circles), the mineral topsoil (filled dots) collected in spring (dark colors) and fall (pale colors). Different fertilization treatments are labeled with colored symbols (red = Con = unfertilized, green = N = nitrogen, purple= P = phosphorus, blue = P+N = phosphorus and nitrogen). Differences between the groups were analyzed by ANOSIM for the factors forest type, soil layer, season and treatment. Data indicate means (n = 24). Abbreviations: RWC = relative water content, Ca = calcium, P = total phosphorous, N = nitrogen; lower case letters s and r at the end of the variables indicate soil and fine roots respectively.

**Table 3:** Correlation of environmental variables with ectomycorrhizal (EMF), root (RAF) and soil (SAF) fungi. Abbreviations: RWC = relative water content, P_tot_ = total phosphorus, Ca = calcium, N = nitrogen; lower case letters s and r at the end of the variables indicate soil and fine roots respectively

| Variable | EMF |  | RAF |  | SAF |  |
| --- | --- | --- | --- | --- | --- | --- |
|  | R<sup>2</sup> | *p* | R<sup>2</sup> | *p* | R<sup>2</sup> | *p* |
| **RWC_s** | 0.32 | **0.027** | 0.05 | **0.038** | 0.26 | **0.001** |
| **RWC_r** | 0.35 | **0.014** | 0.09 | **0.001** | 0.40 | **0.001** |
| **P<sub>tot</sub>_s** | 0.28 | **0.028** | 0.71 | **0.001** | 0.62 | **0.001** |
| **P<sub>tot</sub>_r** | 0.33 | **0.022** | 0.23 | **0.001** | 0.55 | **0.001** |
| **Ca_s** | 0.42 | **0.009** | 0.60 | **0.001** | 0.58 | **0.001** |
| **Ca_r** | 0.46 | **0.003** | 0.21 | **0.001** | 0.33 | **0.001** |
| **N_s** | 0.14 | 0.223 | 0.00 | **0.889** | 0.23 | **0.001** |
| **N_r** | 0.24 | 0.061 | 0.05 | **0.026** | 0.33 | **0.001** |

### 3.4 Phylogenetic and functional groups of fungi respond to N and P treatments

Since important fungal traits for nutrient use and turnover are conserved at the classification levels of the order or phylum (Zanne et al. 2020), we grouped the OTUs of the RAF and SAF according to orders (Supplement Table S7, Supplement Fig. S2). Among the total number of 92 orders present in the SAF and RAF, we chose relatively abundant orders (containing at least >1% of the total reads) to test their responses to N or P addition (Supplement Table S7). The most dominant orders, the Agaricales (27% of the reads) and Heliotales (17% of the reads), were unaffected by the treatments (Fig. 4a, Supplement Table S8, Supplement Table S9).

**Figure 4:**
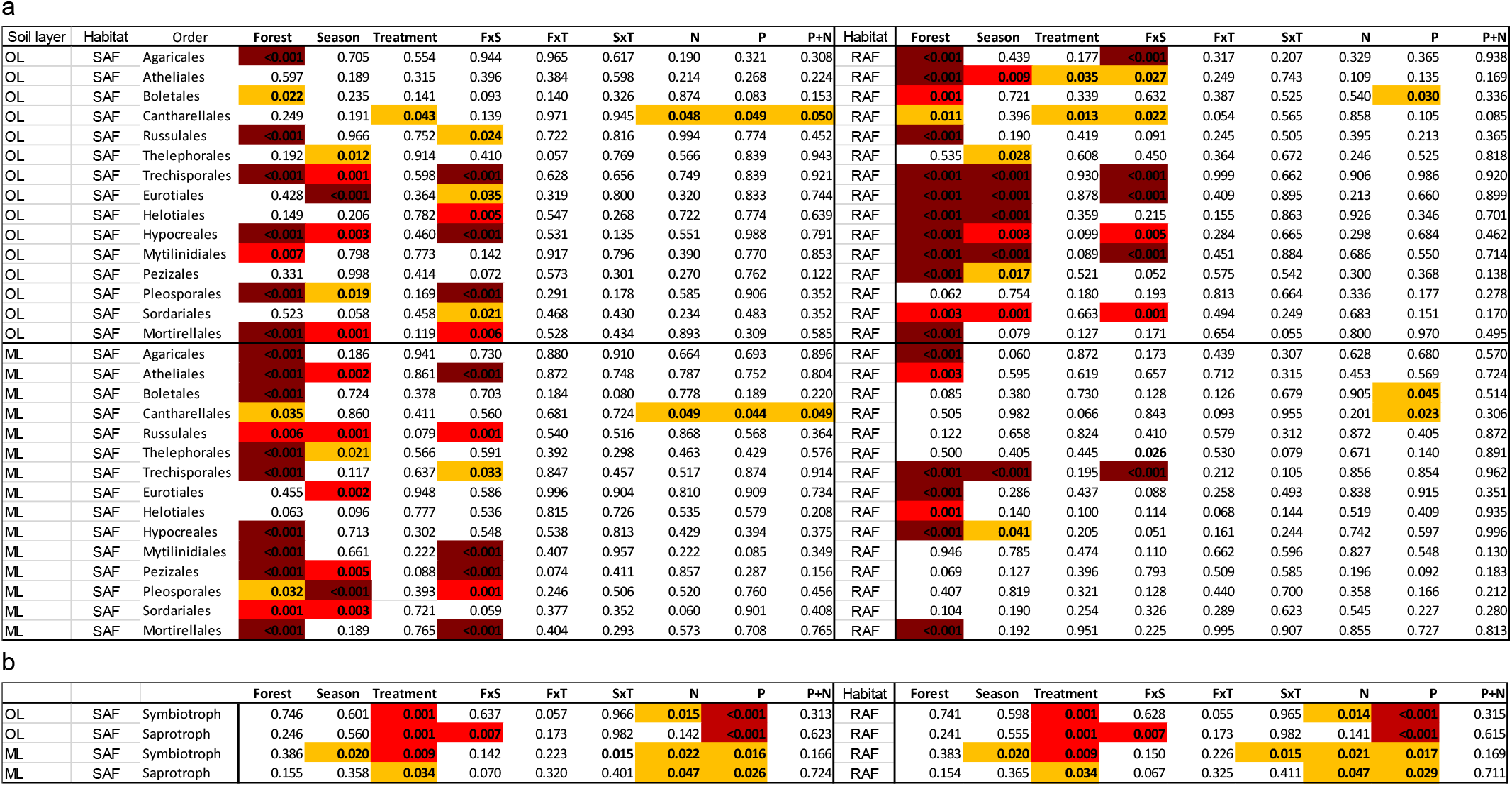
Statistical information on the effects of forest site, season and fertilization treatment on the relative abundance of fungal orders (a) and trophic guilds (b) in the organic layer and mineral topsoil. Soils and roots were collected in a P-rich, P-medium and P-poor forest in spring and fall 2018 and separated into organic layer and the mineral topsoil for analyses. Means can be found in the supplementary information Table S8 and F values in Table S9. Differences among means per forest type, season, treatment and the two-way interactions (FxS: Forest x season, FxT: Forest x treatment, SxT: Season x treatment) were tested by a linear mixed effect model using Poisson distribution with plot number as random effect. Calculations were performed separately for the organic layer and mineral topsoil. Tukey HSD was used as posthoc test. Significant differences are indicated by dark red (p ≤ 0.001), red (0.001 < p ≤ 0.01) and orange (0.01 < p ≤ 0.05).

General treatment effects across all forests, layers and habitats were found in the orders of Atheliales (F = 2.9, *p* = 0.049), Boletales (F = 3.4, *p* = 0.021), Cantharellales (F = 6.4, *p* = 0.001), Russulales (F = 3.8, *p* = 0.013) and Trechisporales (F = 2.8, *p* = 0.043). Across all forests and habitats, the abundance of Atheliales (F = 2.9, *p* = 0.049) decreased in response to P addition and that of the Trechisporales (F = 4.0, *p* = 0.048) in response to N addition. A detailed analysis of treatment effects on these orders in different forests or habitats did not reveal any significant variation (Fig. 4a).

Cantharellales in the SAF were significantly decreased in response to the N, P and P+N treatments (Fig. 5). Russulales were unaffected in the SAF and decreased in the RAF in response to P+N (Fig. 5). Boletales were unaffected in the SAF and increased in response to P in the RAF (Fig. 5).

**Figure 5:**
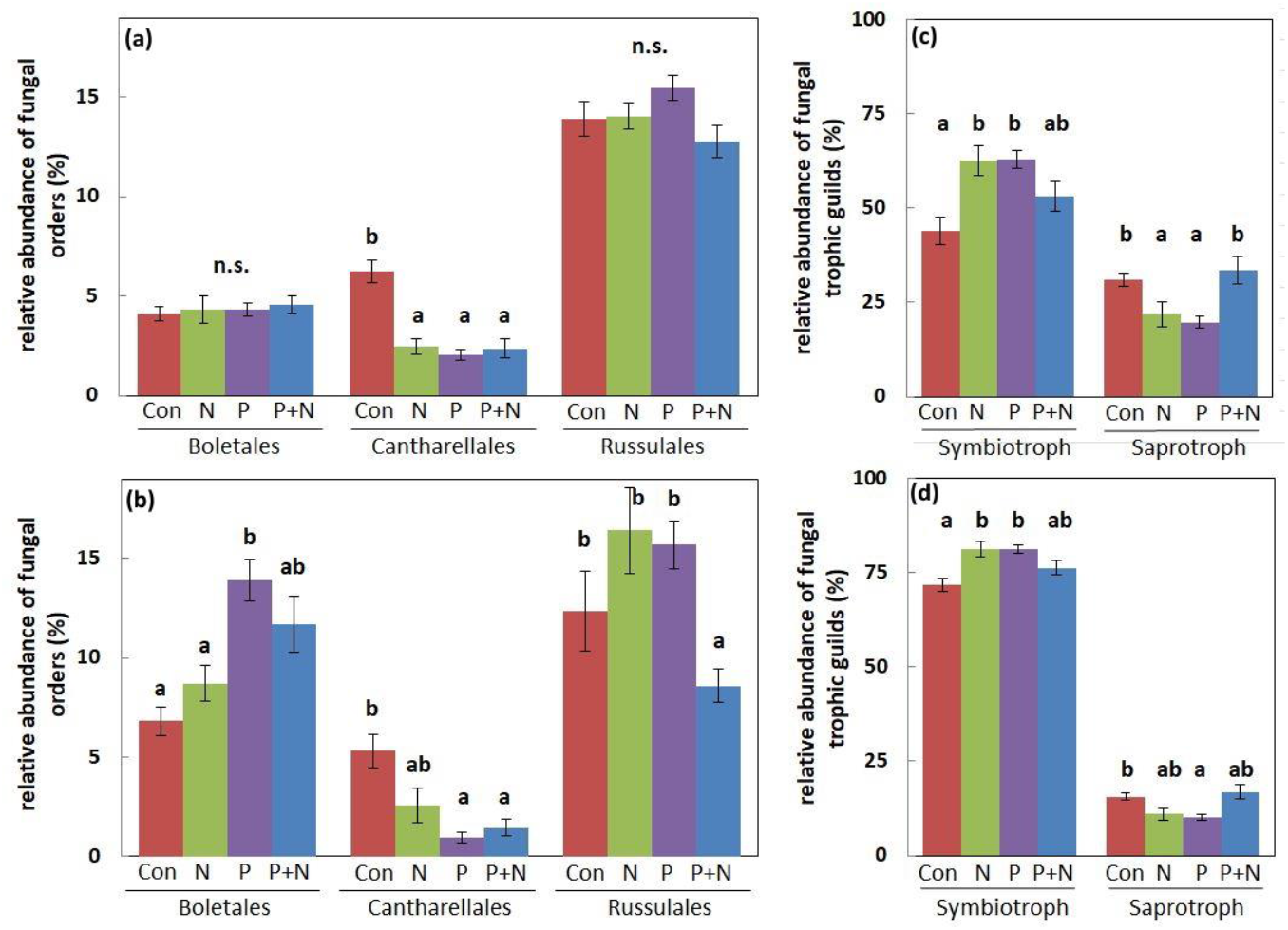
Relative abundance of fungal orders (a-b) and trophic guilds (c-d) in response to fertilization treatments (Con, N, P, P+N)., which were significantly influenced by the treatment. Soil and fine root samples of the organic layer and mineral topsoil were collected in a P-rich, P-medium and P-poor beech forest (*Fagus sylvatica* L.) in spring and fall 2018. Data of the forest types, layers, and season were merged to evaluate effects of the treatments on the soil-residing fungi (A, C) and root-associated fungi (B, D). Data indicate means (n = 36 ± SE). Significant differences between the treatments were calculated by a linear mixed effect model using Poisson distribution and Tukey HSD posthoc test for the treatments with site as random effect and season as repeated measure. Different letters indicated significant differences for each fungal order separately. Controls (Con) = red, N = green P = purple, P+N = blue. N.s. = not significant

The response of the fungi to fertilization treatments varied between soil layers. Cantharellales were negatively affected in both soil layers, while the treatments affected other fungal orders only in the organic layer (Supplement Fig. S3). For instance, the abundance of Boletales and Russulales increased under P fertilization (Supplement Fig. S3).

Since we observed treatment effects in those orders that consisted mainly of EMF, we tested whether symbiotrophs or saprotrophs were influenced by N, P or P+N treatments (Fig. 4b). The relative abundance of symbiotrophic fungi in the SAF and RAF increased, whereas the saprotrophic fungi decreased in these compartments under P and N treatments (Fig. 4b, Fig. 5). We excluded pathotropic fungi from these analyses because their mean relative abundance was below 1%.

Despite the significant differences in soil properties among the forests and soil layers (Table 1, Fig. 1), the relative abundances of symbiotrophic and saprotrophic fungi in soil or on the roots did not show any significant site-related differences (Fig. 4b).

## 4 Discussion

### 4.1 P and N inputs affect P nutrition of beech

In agreement with our first working hypothesis, we found that N addition decreased and that P addition increased the P concentrations in roots under low P soil availabilities. Unexpectedly, we found that N additions also decreased the P concentrations in roots from soils with higher P availabilities, i.e., in the HP and MP forests. These observations suggest that the applied amounts of N (here 60 N kg ha^−1^ a^−1^, other studies: 15-25 kg N ha^−1^a^−1^, De Vries et al. 2014; Wardle et al. 2016; Gonzales and Yanei 2019), which exceed ambient deposition in unpolluted areas (approximately 6 kg N ha^−1^ a^−1^, Schwede et al. 2018), might have caused N:P imbalances. Etzold et al. (2020) reported tipping points at 24–34 kg N ha^−1^ a^−1^ for positive growth responses of forest trees in Europe, with potential negative effects at higher deposition values. In a meta-analysis, Deng et al. (2016) reported decreases in tissue P concentrations upon N fertilization, although the labile P pools in soils were unaffected (Deng et al. 2016). These reports concur with the present results.

A number of previous studies in the LP and HP forests clearly demonstrated that soil microbes and young beech trees in LP soil are limited by low P availabilities (Bergkemper et al. 2016; Lang et al. 2017; Zavišić et al. 2018; Pastore et al. 2020). Experimental studies with young trees in HP and LP soil showed that negative effects of P limitation such as the reduction in photosynthesis were rescued by P fertilization, while the photosynthesis of beech trees in HP soil was unaffected by P addition (Yang et al. 2016; Zavišić et al. 2018). In the present study, the enrichment of NH_4_^+^ in LP but not in HP soil after N addition suggests that N utilization was impaired due to P shortage. The observation that the accumulation of NH_4_^+^ in soil was relieved by P fertilization further supports this suggestion. In line with other studies (Liu et al. 2013; Li et al. 2015; Xia et al. 2020), P addition rescued the negative effects of high N input on root P contents in our experiment.

P addition caused increases in potentially plant-available P in soil in all three forests. This result might have been expected since the annual P uptake of forest trees is much lower (ca. 9 kg ha^−1^ a^−1^, Rosling et al. 2016) than the amount of added P. In our study, the increment in P_lab_ was small compared to the content of P_tot_ in soil and therefore, we did did not observe significant increases in P_tot_. Consequently, the N:P ratios of the soils remained stable after N and P additions. Soil N:P ratios of approximately 15 have been suggested to indicate harmonious nutrition in beech forests (Mellert and Göttlein 2012). Here, we found huge differences among the forests and seasons for these ratios (organic layer: 7 to 31, mineral layer: 3 to 17). However, with regard to tree nutrition, the plant-available fractions of N and P are critical. Here, the N_min_:P_lab_ ratios decreased slightly in response to P or N addition in the HP forest but increased more than 3-times after N addition in the organic layer of the LP forest compared to that in the HP forest (estimated with data from the organic layer, Supplement Table S4). This dynamics was partly also detected in roots, where N fertilization caused decreases in P_tot_, while P fertilization caused increases in P_lab_, supporting metabolic flexibility of beech to cope with differences in nutrient availabilities (Zavisic et al. 2016; Zavisic and Polle 2018; Meller et al. 2019).

### 4.2 Fungal taxonomic structures are driven by long term habitat conditions

A main goal of this study was to disentangle the responses of the soil fungal community structures to addition of N, P and P+N under conditions of P shortage or P sufficiency. In general, the assembly processes of soil fungi are predominately driven by deterministic processes such as abiotic habitat conditions, soil properties, etc., while stochastic effects play a minor role in shaping the community structures (Chase 2007; Štursová et al. 2014; Mykrä et al. 2016; Peay et al. 2016; Glassman et al. 2017). In agreement with the expectation that abiotic habitat filters were important drivers of the fungal community structure, our results show that environmental variables such as humidity, P, Ca and N in soil and roots explained differences among fungal community structures in different forests.

In line with other studies (Goldmann et al. 2016; Zhang et al. 2017; Luo et al. 2019) RAF were less diverse than SAF. Furthermore, the EMF assemblage involved in active symbiosis was by far less diverse than the EMF detected by Illumina or other deep sequencing methods (Danielsen et al. 2012; Pena et al. 2017; Schröter et al. 2019). For example, Pena et al. (2017) found about 10 to 15 EMF species colonizing root tips per plot, while Schröter et al. (2019) detected about 50 EMF species by pyrosequencing in the same plots. Our results for EMF were in a similar range. The high number of OTUs is partly due to a methodological bias (Nilsson et al. 2019; Castaño et al. 2018) resulting in species overestimation. However, the enhanced EMF species richness discovered in the SAF and RAF compared to EMF colonizing root tips also reflects the ability of EMF, which are not engaged in an active symbiosis, to live as saprotrophs in soil or on root surfaces (Iwański and Rudawaska 2007; Phillips et al. 2014; Kohler et al. 2015; Lindahl and Tunlid 2015).

According to ecological theory, stress results in reduction of diversity, filtering out species that can tolerate harsh environments (Chase 2007). This theory has also support for soil fungi including ectomycorrhizal communities (Štursová et al. 2014; Glassman et al. 2017; Schröter et al. 2019). For example, along a biogeographic gradient in temperate forests, the fungal assemblages were generally less diverse in dry and more acidic soil than those in a more humid and nutrient rich soil (Schröter et al. 2019; Goldmann et al. 2016; Pena et al. 2017). Therefore, we antipicated here that P fertilization resulted in stress relief leading to more species-rich, diverse assemblages. While no effect on EMF was found, the influence of P fertilization on fungal richness caused a moderate increment of the saprotrophic fungal richness (+7%) in the organic layer at the LP forest. Analyses of the taxonomic fungal community composition did not reveal any significant effect in response to the fertilization treatments, thus, indicating that the taxonomic dissimilarities and species turnover among the forests overruled small effects at the level of OTUs.

As outlined in the introduction, high N inputs often caused reductions in fungal diversity and shifts in the community towards nitrophilic assemblages (Lilleskov et al. 2002, 2008; Cox et al. 2010; Bahr et al. 2013; Suz et al. 2014; de Witte et al. 2017). Field studies in temperate forests also identified soil N as an important driver of RAF structures (Schröter et al. 2019; Lilleskov et al. 2019; Nguyen et al. 2020). In other studies, N deposition did not influence fungal community structures (Purahong et al. 2018; Lilleskov et al. 2019). Similarly, we did not observe effects of N addition on the fungal assemblages, irrespective of whether the fungi in soil or those in direct contact with the roots were inspected. Upon N fertilization, less carbon is allocated belowground to ectomycorrhizal fungi associated with roots (Högberg et al. 2017, Högberg et al. 2020). Reduced carbon availability to the EMF causes a reduction in P uptake of the roots (Clausing et al. 2020). These physiological feed-back controls might have caused the decreased P concentrations in roots after N addition found in our study without requiring strong reshaping of the fungal assemblage. In conclusion, our hypothesis that P fertilization increases fungal richness in P-poor soil had no support for EMF.

### 4.3 N and P inputs affect phylogenetic and functional structures of fungal assemblages

Phylogenetic structures carry information on ecological assembly processes because the relatedness between members of a community suggests similar ecological requirements or functions (Cavender-Bares et al. 2009; Pausas & Verdú 2010). Treseder and Lennon (2015) analysed fungal traits required for nutrient cycling (e.g. phosphatase, ammonium transporters) in fungal genomes. They found that these traits were more conserved in terms of gene counts in phylogenetically more closely related taxa (up to the subphylum level) than in the more distant ones (Treseder and Lennon 2015). Therefore, we reasoned that adaptation of fungal community structures to enhanced N or P inputs might be traceable after aggregation of OTUs to higher classification levels at the level of orders. Applying this approach, we found that Cantharellales encompassed apparently the most susceptible group of fungi to environmental nutrient inputs since their abundance decreased in response to N or P in soil and roots. Addition of organic or inorganic fertilizers to paddy soils also decreased the abundances of Cantharellaes (Nie et al. 2018) but in a boreal conifer forest no significant effects were found for this fungal order upon the addition of N fertilizers (Allison et al. 2007). However, it should be noted that Cantharellales were not very abundant and therefore, subtle decreases in abundance are difficult to uncover. Nevertheless, we consider the decreases critical because Cantharellales contain valuable edible mushrooms such as chanterelle (*Cantharellus cibarius*). Long-lasting inadvertent nutrient inputs, even with only small effects on the whole system, thus, may lead to a decline in an important cultural forest service (Felipe-Lucia et al. 2018).

Members of the russula lineage formed a dominant group in our study forests (Zavišić et al. 2016; Clausing et al. 2020; this study). All known members of the Russulaes are ectomycorrhizal fungi and are very common in temperate beech forests (Buée et al. 2005; Lang et al. 2011; Pena et al. 2010). Here, Russulales showed a significant decrease in response to the combined P+N treatment but tended to increase in the organic layer, when N or P were applied as single factors. *Russula* sp. lack extensive extramatrical hyphae and, thus, absorb nutrients from their immediate surroundings (Agerer 2001). Litter-derived nitrogen is available to *Russula* sp. with a delay of approximately one year in temperate beech forests, indicating low saprotrophic properties (Pena et al. 2013). Therefore, elevated inorganic nutrient availability may favour this fungal genus. Similarly to our study, Mason et al. (2020) found subtle increases in Russulales after P fertilization (5 years) of an LP forest (Ohio, USA). Nicolás et al. (2017) found no significant effects of N fertilization on *Russula* sp. in a boreal forest. However, root colonization and sporocarp formation of *Russula* sp. increased significantly after strong long-term disturbance by high N input (16 years, 170 kg N ha^−1^ a^−1^) (Avis et al. 2003). Therefore, various *Russula* sp. were classified as nitrophilic species (Avis, 2012). Our results indicate that N availability alone was not decisive. Rather the N:P ratio regulated the abundance of this important fungal order as Russulaes declined significantly when high N addition was accompanied by high P availabilities.

P fertilization exerted the most pronounced effects on Boletales. The members of this order are characterized by long-distance rhizomorphs and thick hyphal mantle (Agerer 2001). The relative abundance of Boletales increased almost twofold upon P addition. This result was surprising because the investment into high-biomass fungi is considered profitable in nutrient-limited ecosystems to access distant resources (Hobbie and Agerer 2010). For example, Almeida et al. (2019) found increases in *Imleria badia* (formerly known as *Boletus badius*) hyphae accessing apatite (a recalcitrant P source) in N fertilized soil but not if the fertilized N soil was amended with easily available P sources. Therefore, they argued that *Imleria* is a P-efficient species that responds to an enhanced P demand of the tree (Almeida et al. 2019). However, our results do not support this suggestion because the roots of N fertilized plots showed decreases in P, which would support an enhanced tree-demand under these conditions, whereas the increases in Boletales occurred only in P or P+N fertilized plots. Boletales accumulate P in the hyphal mantle and store P in polyphosphate granules in the mycelium (Kottke et al. 1998). Therefore, our results suggest that the Boletales enhanced P supply to the host and were responsible for P accumulation found in roots from the P-fertilized plots.

In addition to fertilizer treatments, the fungal orders also showed differences between the soil layers, especially for the saprotrophic fungi. Other studies also reported shifts in fungal composition with increasing soil depth (Toju et al. 2016; Peršoh et al. 2018). For example, Toju et al (2016) found a lower abundance of Russulales in the organic layer than in deeper horizons. However, this observation deviates from our results. We found that orders mainly harboring mycorrhizal fungal species did not vary between the layers (this study; Clausing and Polle 2020), whereas orders known to contain many saprotrophic species showed significant differences between the organic layer and the mineral soil (organic layer: Hypocreales, Pleosporales, Sordariales; mineral topsoil: Trechisporales, Pezizales, Mortierellales) (Supplement Table S8). In general, the saprotrophic taxa were significantly less abundant in the RAF than in the SAF and tended to lower abundances in the mineral topsoil compared to the organic layer. This pattern reflects different nutritional strategies of saprotrophs and mycorrhizal fungi. Saprotrophs prefer environments where bound nutrients can be unlocked from organic compounds, while mycorrhizal fungi mine the mineral layer for inorganic compounds and rely on plant-derived carbohydrates.

An important result of our study was that the relative abundance of saprotrophic fungi was reduced in response to N as well as to P fertilization. This observation suggests that an enhanced availability of mineral nutrients occurs with a trade-off in saprotrophic potential. The shift towards symbiotrophs suggests that addition of inorganic nutrients leads to a competitive advantange for growth of ectomycorrhizal fungi, because they obtain carbohydrates from their host, while free-living saprotrophic fungi need to degrade organic compounds to access carbon. Field experiments showed that enhanced ectomycorrhizal fungal growth also results in enhanced phosphodiesterase activities, and thus higher P_lab_ availability (Müller et al. 2020). Pot experiments under controlled conditions found that ectomycorrhizal diversity fostered P uptake efficiency (Köhler et al. 2018) and in forest soils ectomycorrhizal P uptake efficiency was further related to P_lab_ availability (Clausing and Polle 2020). These results suggest that P_lab_ availability sets off a positive feed-back mechanism for plant nutrition. The increase in symbiotrophic relative to saprotrophic fungal abundances upon N or P addition supports that the ectomycorrhizal fungi are superior to saprotrophs in capturing mineral nutrients in beech forests. In contrast to enhanced P_lab_ availability, enhanced N_min_ availability resulted in a decrease in root P contents. The shift away from saprotrophic towards symbiotrophic activities may have resulted in lower P mineralization, thereby, decreasing P availability and contributing to the reduced P content in roots of the N fertilized plot. These considerations underline the important role of saprotrophic fungi for the mineralization of organic P.

## 5 Conclusion

Our study shows a strong resistance of fungal community structures (based on OTUs) to N and P fertilization. However, shifts in the composition of fungal assemblages in response to N or P were manifested at higher phylogenetic levels, thus, supporting that genetic relatedness can be applied as a proxy for trait-based approaches in complex microbial communities (Amend et al. 2016; Treseder and Lennon 2015; Nguyen et al. 2020).

In contrast to our hypothesis, we found a decline in the root P contents under N fertilization, irrespective of the soil P contents. Furthermore, increases in labile P concentrations in the organic layer and the mineral soil occurred after P fertilization, irrespective of the soil P stocks and were accompanied by unexpected increases in the abundance of ectomycorrhizal fungi but not by the appearance of new species. A moderate increase in fungal richness, but a decline in their abundance was confined to the saprotrophs in the organic layer. Overall, our results underline the importance to distinguish different habitats and to include the major nutrients N and P to better understanding the drivers of fungal communities in relation nutrient cycling. Although we did not find general decreases in fungal richness or abundances in response to high N input, we observed declines in distinct orders such as the Cantharellales or Russulales under P+N treatment. Since Cantharellales are known for valuable edible fungi and Russulales contained a number of endangered species (*R*. *veternosa, R. melzeri* and *R. brunneoviolacea*), it is important to protect beech forest ecosystems from uncontrolled nutrient inputs in inorganic forms to prevent impoverishment of fungal richness. Increased availabilities of mineral P and N shifted the functional composition towards increased abundances of ectomycorrhizal fungi. These shifts may lead to nutrient imbalances, when the mineralization of limited nutrients is impaired.

## Supporting information

all supplements

## 6 Declarations

### 6.2 Authors‘ contributions

SC and AP conceived the study. FL and JK realized the fertilization experiment, maintained the plots and contributed field data. SC conducted field and laboratory measurements. LEL conducted EMF sampling and analyses. HYF determined CN. SC, LEL, DJ, DS, RD, and AP analyzed data. SC compiled the data and wrote the first manuscript draft. AP revised the draft. All authors contributed to, commented and agreed on the final version.

### 6.4 Data availability

The datasets generated for this study can be found in the Dryad digital repository (<u>xxx</u>). The sequences for identified mycorrhizal fungi are available in NCBI GenBank under accession numbers MT859114 to MT859131. The Illumina MiSeq sequences of fungal ITS2 can be found in the Sequence Read Archive (SRA) of NCBI under Bioproject <u>PRJNA680926</u>.

### 6.5 Ethical statement

Not required

## 7 Acknowledgements

We are grateful to Nadine Kubsch and PhD Karolin Müller for help with the harvests. We appreciate the skilled technical assistance by M. Franke-Klein (Forest Botany and Tree Physiology, Göttingen), G. Lehmannn, and T. Klein (Laboratory for Radio-Isotopes, LARI, Göttingen).

