## Supplementary material for "Impact of nitrogen and phosphorus addition on resident soil and root mycobiomes in beech forests": all supplements

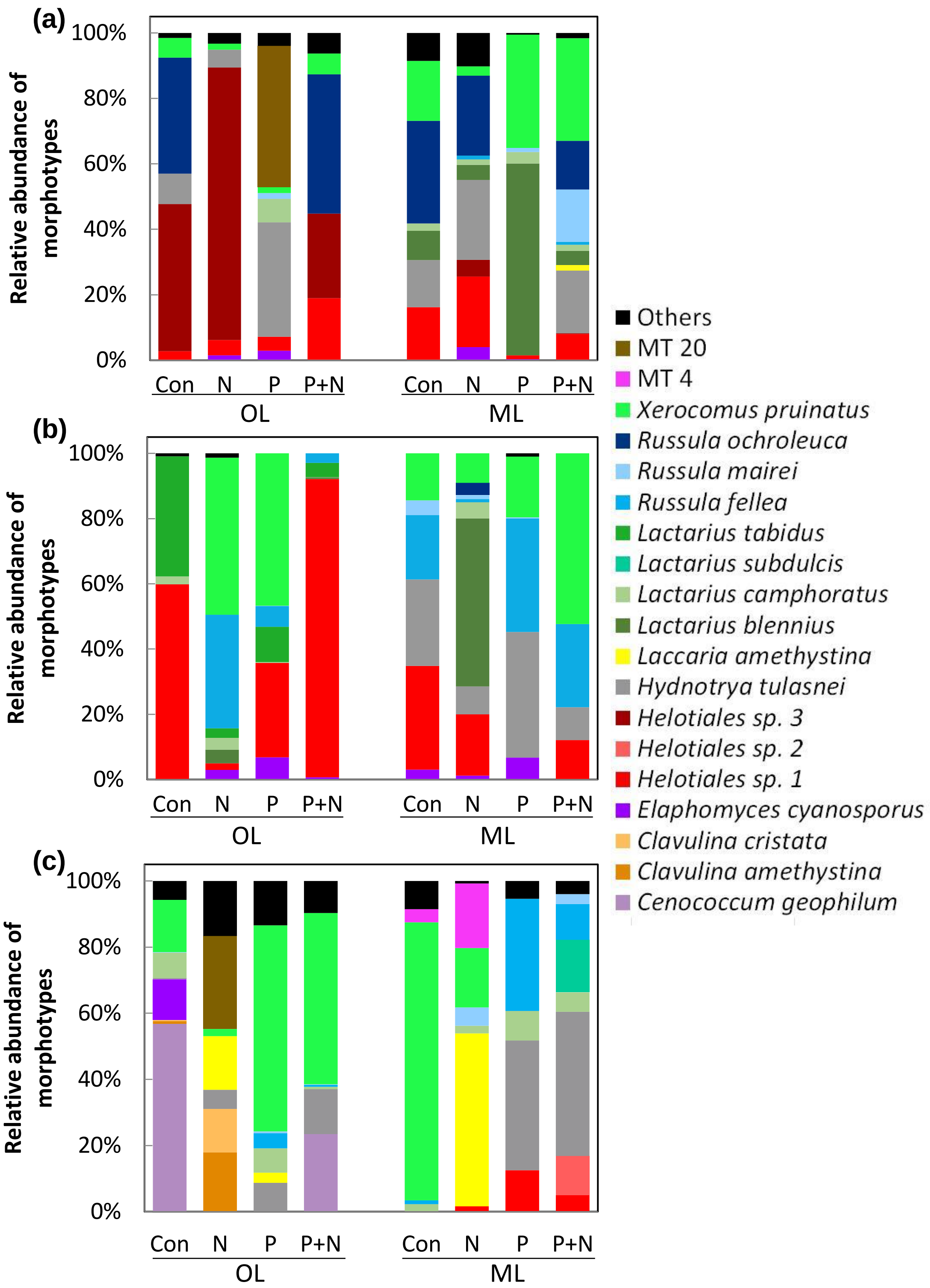


**Supplement Figure S1: Ectomycorrhizal fungal community colonizing beech (*Fagus sylvatica* L.) roots under different fertilization treatments (Con, N, P, P+N).** Trees were investigated in P-rich (a), P-medium (b) and P-poor (c) forests. Roots from the organic layer and the mineral topsoil were analyzed separately. Two fungal morphotypes (MT20, MT4) did not yield sequences. “Others” refers to the sum of rare morphotypes, which were not sequenced. Data indicate means (n = 3). ANOSIM revealed significant differences between the following groups: Forest: R = 0.171, $p$ = 0.001; Layer R = 0.018, $p$ = 0.171; Forest type x Layer: R = 0.211, $p$ = 0.001.

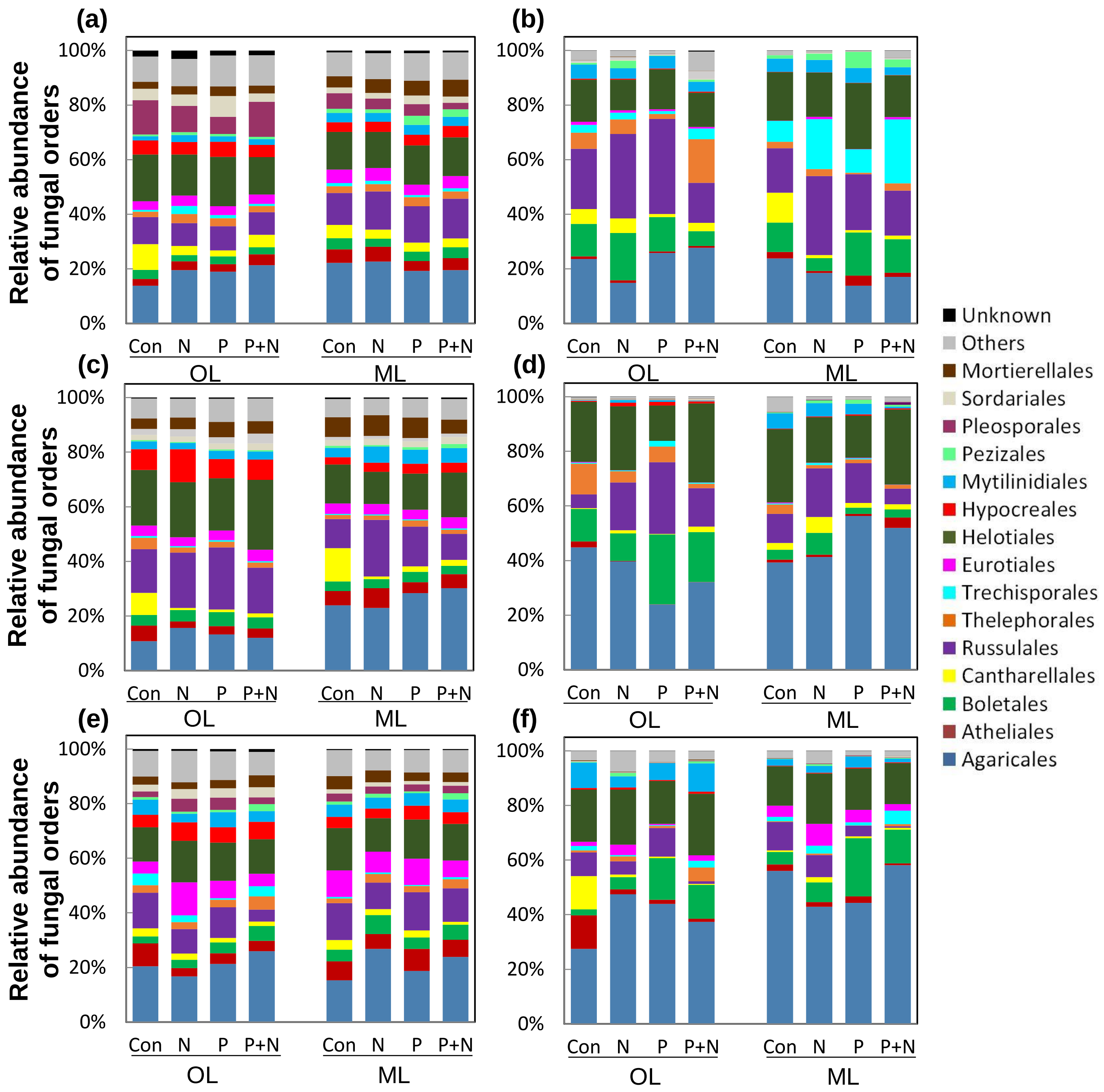


**Supplement Figure S2: Relative abundance of fungal orders residing in soil (a, c, e) and associated with roots (b, d, f) of beech forests (*Fagus sylvatica* L.) without fertilization (Con) or with addition of N, P or N+P.** Soil and fine root samples were collected in a P-rich (a, b), P-medium (c, d) and P-poor (e, f) forest in 2018. Fungi in all forests, soil layers, seasons and compartments were analyzed separately by Illumina sequencing. Samples from the spring and fall season were pooled for the analyses. Data indicate means (n = 6). Fungal order with > 1% of the sequences are shown: Basidiomycota: Agaricales – dark blue, Atheliales – dark red, Boletales - green, Cantharellales – yellow, Russulales – purple, Thelephorales – orange, Trechisporales - turqouise ;Ascomycota: Eurotiales – pink, Helotiales – dark green, Hypocreales – red, Mytinidiales – blue, Pezizales – mint green, Pleosporales – sand, Sordariales – dark purple ; Zygomycota: Mortierellales – brown; Others (= sum of fungal orders < 1% of the sequences) – grey, Unknown (= sum of fungal sequences without an annotation for a fungal order) – black. Significant differences between the forest, season and treatment were calculated by a linear mixed effect model using Poisson distribution with plot number as random effect. Data are shown in Table 4.

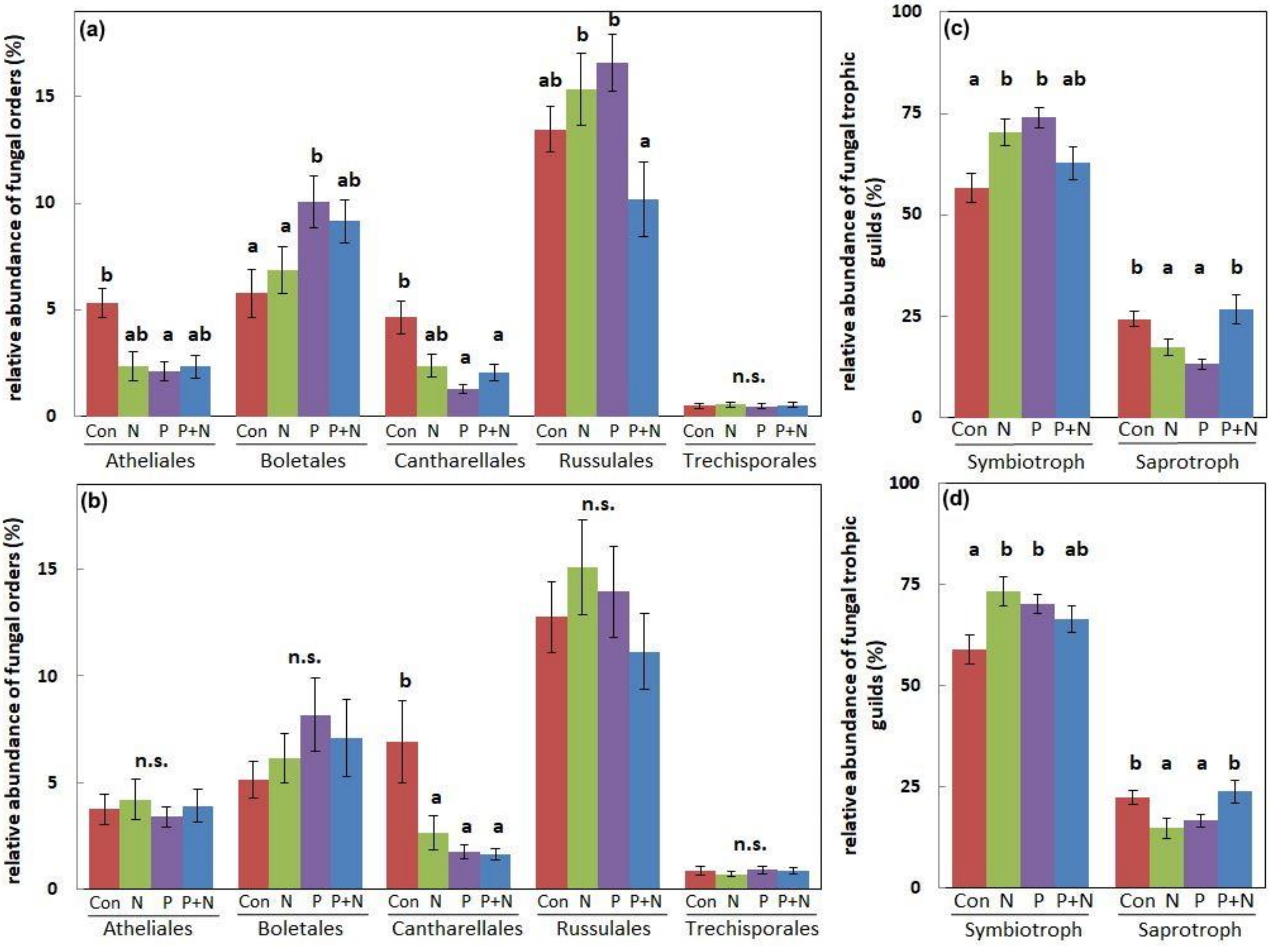


**Supplement Figure S3: Relative abundance of fungal orders (a-b) and trophic guilds (c-d) in response to different fertilization treatments (Con, N, P, P+N).** Fungal orders with significant effects are shown. Soil and fine root samples of the organic layer and mineral topsoil were collected in a P-rich, P-medium and P-poor beech forest (*Fagus sylvatica* L.) in spring and fall 2018. We merged the data of the forest types, habitat (soil, roots) and season (spring and fall) to evaluate effects by the treatments in the organic layer (a, c) and mineral topsoil (b, d). Data indicate means (n = 36 ± SE). Significant differences between the treatments were calculated by a linear mixed effect model using Poisson distribution and Tukey HSD posthoc test for the treatments with site as random effect and season as repeated measure. Different letters indicated significant differences for each fungal order separately. Con = red, N = green, P = purple, P+N = blue. n.s. = not significant

**Supplement Table S1: Characteristics of the research sites in the P-rich (HP, Bad Brückenau) the P-medium (MP, Mitterfels) and the P-poor forest (LP Luess).** Data were compiled from Lang et al. (2017). The parameters age, height and diameter refer to beech trees. Extractable P was determined with the resin method (Lang et al. 2017).

| Parameters | HP | MP | LP |
|---|---|---|---|
| **Location** | | | |
| Gauss-Krüger coordinates | 50°21'7.2"N<br>9°55'44.5"E | 48°58'34.1"N<br>12°52'46.7"E | 52°50'21.7"N<br>10°16'2.3"E |
| Altitude (m a.s.l.) | 809 | 1023 | 115 |
| **Climate** | | | |
| Mean annual temperature (°C) | 5.8 | 4.9 | 8.0 |
| Sum of annual precipitation (mm) | 1031 | 1200 | 779 |
| **Stand characteristics** | | | |
| Potential natural vegetation | Hordelymo-Fagetum | Dryopteris-Fagetum | Luzulo-Fagetum |
| Tree species composition (%) | *Fagus sylvatica* (99)<br>*Acer pseudoplatanus* (1) | *Fagus sylvatica* (96)<br>*Picea abies* (2)<br>*Abies alba* (2) | *Fagus sylvatica* (91)<br>*Quercus petraea* (9) |
| Age (a) | 137 | 131 | 132 |
| Height (mean tree) (m) | 26.8 | 20.8 | 27.3 |
| Diameter at breast height (cm) | 36.8 | 37.6 | 27.5 |
| Number of trees ($ha^{-1}$) | 335 | 252 | 480 |
| Basal area ($m^2$ $ha^{-1}$) | 35.6 | 28.1 | 36.7 |
| Standing volume ($m^3$ $ha^{-1}$) | 495 | 274 | 529 |
| **Soil characteristics** | | | |
| Soil type | Dystric skeletic cambisol | Hyperdystric chromic folic cambisol | Hyperdystric folic Cambisol |
| Parent material | Basalt | Paragneiss | Sandy till |
| Humus form | Mull-like Moder | Moder | Mor-like Moder |
| Texture (topsoil) | Silty clay loam | Loam | Loamy sand |
| Texture (subsoil) | Loam | Sandy loam | Sand |
| **Soil chemistry (A horizon 0 to 5 cm)** | | | |
| pH ($H_20$) | 3.8 | 3.6 | 3.5 |
| Total P (mg $kg^{-1}$) | 2966 | 1375 | 195 |
| Extractable P (mg $kg^{-1}$) | 116 | 70 | 11 |
| P in leaf litter (g $m^{-2}$ $a^{-1}$) | 0.229 | 0.213 | 0.156 |
| P in leaves (mg $g^{-1}$ dry mass) | 1.41 | 1.66 | 1.21 |

**Supplement Table S2: Weather conditions (temperature and precipitation) during the harvest season and deviation from climatic conditions in beech forests (*Fagus sylvatica* L.).** Soil samples were collected in a P-rich (HP), P-medium (MP) and P-poor (LP) forest in spring and fall 2018. We show the weather conditions in the month before and in the sampling month, using mean monthly temperatures and the sum of precipitation. The deviation from the long-term climatic conditions (dfc) was calculated as the monthly mean temperature of the sampling month minus the long-term mean in the respective month for the period 1981-2010. Data indicate monthly means of March and April for spring and of August and September for fall. Spring samples collected in April to early May and fall samples were collected in September to early October. The second table shows the weather and climatic data for each month.

| | temperature (°C) | | precipitation (mm) | |
|---|---|---|---|---|
| | Spring | fall | Spring | fall |
| during the harvest + month before | | | | |
| **HP** | 7.4 | 13.2 | 120.4 | 63.1 |
| **MP** | 6.8 | 12.1 | 57.1 | 110.6 |
| **LP** | 7.7 | 13.3 | 53.3 | 64.0 |
| deviation from climatic conditions | | | | |
| **HP** | +1.9 | +2.7 | -17.9 | -96.6 |
| **MP** | +0.6 | +1.5 | -37.5 | -13.9 |
| **LP** | +0.6 | +1.4 | -8.4 | -49.8 |

| | | **temperature (°C)** | | **precipitation (mm)** | |
|---|---|---|---|---|---|
| **site** | **month** | month | dfc | Month | dfc |
| | February | -3.1 | -2.8 | 13.6 | -59.8 |
| | March | 1.8 | -1.6 | 70.3 | -10.1 |
| | April | 13.0 | 5.3 | 50.1 | -7.8 |
| | May | 16.1 | 3.8 | 61.4 | -10.1 |
| **HP** | June | 17.6 | 2.6 | 46.3 | -20.3 |
| | July | 21.1 | 3.9 | 32.1 | -44.2 |
| | August | 20.2 | 3.3 | 26.3 | -35.9 |
| | September | 15.2 | 2.3 | 36.3 | -37.2 |
| | October | 11.1 | 3.0 | 26.8 | -59.7 |
| | February | -3.5 | -3.4 | 20.6 | -24.7 |
| | March | 1.3 | -2.3 | 43.8 | -6.8 |
| | April | 12.3 | 3.4 | 13.3 | -30.7 |
| | May | 15.5 | 3.1 | 67.9 | -28.0 |
| **MP** | June | 16.8 | 0.6 | 49.8 | -44.1 |
| | July | 19.0 | 1.2 | 68.2 | -39.9 |
| | August | 19.7 | 2.6 | 33.0 | -55.5 |
| | September | 14.1 | 1.1 | 73.9 | 7.2 |
| | October | 10.1 | 1.9 | 36.7 | -21.1 |
| | February | 1.3 | -3.1 | 3.2 | -17.1 |
| | March | 2.6 | -2.3 | 3.5 | -16.5 |
| | April | 12.7 | 3.5 | 49.8 | 8.1 |
| | May | 17.5 | 3.7 | 38.7 | -14.5 |
| **LP** | June | 18.4 | 2.0 | 14.1 | -47.0 |
| | July | 21.1 | 2.4 | 39.2 | -27.3 |
| | August | 20.2 | 2.1 | 30.1 | -36.7 |
| | September | 15.4 | 1.3 | 43.8 | -13.5 |
| | October | 11.1 | 1.4 | 20.2 | -36.3 |

**Supplement Table S3: Molecular information on ectomycorrhizal species colonizing root tips in beech forests (*Fagus sylvatica* L.) under fertilization treatments.** Roots were analyzed in P-rich, P-medium and P-poor forests. Roots from the organic layer and the mineral topsoil were analyzed separately. The table shows the original morphotype number, accession number of the best match (best match in the UNITE Genbank), the sequence length/length in the data base and % similarity of nucleotide sequence, and NCBI accession number (accession number under which the sequences of the present fungi have been deposited in NCBI Genbank). The ITS region of the fungal rDNA was amplified using the PCR primers ITS1F (5′CTTGGTCATTTAGAGGAAGTAA-3′) and ITS4 (5′TCCTCCGCTTATTGATATGC-3′) (White et al. 1990).

| Ectomycorrhizal fungal species | MT number | Reference accession number UNITE | Length and % similarity of nucleotide sequence | NCBI accession number |
|---|---|---|---|---|
| *Cenococcum geophylum* | MT26 | LC095162.1 | 542 (98.9%) | MT859114 |
| *Clavulina amethystina* | MT19 | MK422194.1 | 736 (96.66%) | MT859115 |
| *Clavulina cristata* | MT11 | MN947349.1 | 735 (99.03%) | MT859116 |
| *Elaphomyces cyanosporus* | MT33 | KX238826.1 | 613 (99.83%) | MT859117 |
| *Helotiales sp. 1* | MT1 | HM190117.1 | 736 (99.73%) | MT859118 |
| *Helotiales sp. 2* | MT34 | JF519582.1 | 621 (99.67%) | MT859119 |
| *Helotiales sp. 3* | MT35 | LC189022.2 | 576 (100%) | MT859120 |
| *Laccaria amethystina* | MT16 | MN947342.1 | 736 (100%) | MT859121 |
| *Lactarius blennius* | MT39 | MN947353.1 | 782 (99.23%) | MT859122 |
| *Lactarius camphoratus* | MT8 | MN992440.1 | 763 (99.86%) | MT859123 |
| *Lactarius subdulcis* | MT9 | HM189802.1 | 765 (99.61%) | MT859124 |
| *Russula fellea* | MT6 | MN959791.1 | 694 (99.63%) | MT859126 |
| *Russula mairei* | MT2 | MN947352.1 | 704 (99.15%) | MT859127 |
| *Russula ochroleuca* | MT38 | MT644930.1 | 741 (99.73%) | MT859128 |
| Xerocomus pruinatus | MT5 | MN947367.1 | 800 (99.38%) | MT859129 |
| *Hydnotrya tulasnei* | MT15 | GQ149454.1 | 762 (99.87%) | MT859130 |
| *Lactarius tabidus* | MT42 | HM189817.1 | 752 (99.6%) | MT859131 |
| Unknown - MT 4 | MT4 | | | |
| Unknown - MT 20 | MT20 | | | |

**Supplement Table S4: Soil and root chemistry in the organic layer and mineral top soil in spring and fall in response to fertilization treatments (Con, N, P, P+N) of beech forests (*Fagus sylvatica* L.).** Soils and roots were collected in a P-rich (HP), P-medium (MP) and P-poor (LP) forest in spring and fall 2018 and separated into organic layer and the mineral topsoil for analyses. Data indicate means (n = 3 ± SE). RWC = relative water content. Statistical information is provided in Table 1

**HP**

| **Spring** | | | | |
|---|---|---|---|---|
| **Organic layer** | **Con** | **N** | **P** | **P+N** |
| **Bulk soil** | | | | |
| RWC | 0.54 ± 0.03 | 0.42 ± 0.02 | 0.52 ± 0.07 | 0.46 ± 0.04 |
| $P_{tot}$ (mg $g^{-1}$ dw) | 2.59 ± 0.12 | 2.29 ± 0.11 | 2.08 ± 0.12 | 1.65 ± 0.07 |
| $P_{lab}$ (µg $g^{-1}$ dw) | 264.6 ± 25.1 | 286.6 ± 27.7 | 313.7 ± 18.4 | 354.7 ± 30.8 |
| N (mg $g^{-1}$ dw) | 17.7 ± 0.8 | 17.5 ± 0.9 | 19.2 ± 0.7 | 17.0 ± 0.9 |
| C (mg $g^{-1}$ dw) | 365.8 ± 21.4 | 349.4 ± 11.7 | 377.5 ± 10.3 | 329.5 ± 22.2 |
| C:N | 20.7 ± 0.3 | 20.0 ± 0.8 | 19.7 ± 0.2 | 19.4 ± 1.0 |
| N:P | 6.8 ± 0.5 | 7.7 ± 0.7 | 9.3 ± 0.6 | 10.4 ± 0.9 |
| pH | 4.51 ± 0.05 | 4.62 ± 0.07 | 4.53 ± 0.07 | 4.70 ± 0.07 |
| $NH_4^+$ (µmol $g^{-1}$ dw) | 402.9 ± 232.7 | 366.5 ± 81.2 | 562.3 ± 40.2 | 600.4 ± 214.1 |
| $NO_3^-$ (µmol $g^{-1}$ dw) | 39.1 ± 9.5 | 51.2 ± 6.6 | 56.0 ± 10.9 | 28.4 ± 11.9 |
| **Fine roots** | | | | |
| RWC | 0.71 ± 0.02 | 0.70 ± 0.02 | 0.71 ± 0.02 | 0.72 ± 0.03 |
| $P_{tot}$ (mg $g^{-1}$ dw) | 2.78 ± 0.07 | 1.97 ± 0.06 | 2.51 ± 0.04 | 2.16 ± 0.14 |
| $P_{lab}$ (µg $g^{-1}$ dw) | 510.1 ± 32.7 | 637.1 ± 59.3 | 733.1 ± 13.5 | 687.8 ± 77.3 |
| N (mg $g^{-1}$ dw) | 21.7 ± 0.7 | 20.7 ± 1.7 | 22.4 ± 1.2 | 21.0 ± 1.1 |
| C (mg $g^{-1}$ dw) | 471.4 ± 2.5 | 464.1 ± 1.9 | 463.6 ± 9.4 | 467.0 ± 1.7 |
| C:N | 21.7 ± 0.6 | 22.6 ± 1.6 | 20.8 ± 0.9 | 22.3 ± 1.3 |
| N:P | 7.8 ± 0.2 | 10.6 ± 1.2 | 8.9 ± 0.5 | 9.8 ± 0.6 |
| **Mineral topsoil** | | | | |
| **Bulk soil** | | | | |
| RWC | 0.46 ± 0.01 | 0.46 ± 0.01 | 0.45 ± 0.01 | 0.45 ± 0.01 |
| $P_{tot}$ (mg $g^{-1}$ dw) | 3.33 ± 0.14 | 3.42 ± 0.15 | 3.60 ± 0.14 | 3.78 ± 0.16 |
| $P_{lab}$ (µg $g^{-1}$ dw) | 233.0 ± 59.4 | 220.2 ± 34.6 | 325.3 ± 28.5 | 230.1 ± 15.7 |
| N (mg $g^{-1}$ dw) | 7.40 ± 0.52 | 7.56 ± 0.32 | 7.21 ± 0.55 | 7.70 ± 0.81 |
| C (mg $g^{-1}$ dw) | 107.3 ± 9.4 | 103.5 ± 3.4 | 104.4 ± 7.4 | 108.7 ± 14.0 |
| C:N | 14.5 ± 0.6 | 13.7 ± 0.2 | 14.5 ± 0.1 | 14.1 ± 0.3 |
| N:P | 2.2 ± 0.2 | 2.2 ± 0.1 | 2.0 ± 0.2 | 2.0 ± 0.2 |
| pH | 4.54 ± 0.25 | 4.51 ± 0.10 | 4.53 ± 0.12 | 4.54 ± 0.07 |
| $NH_4^+$ (µmol $g^{-1}$ dw) | 7.26 ± 2.91 | 12.6 ± 1.3 | 18.1 ± 3.4 | 64.2 ± 50.6 |
| $NO_3^-$ (µmol $g^{-1}$ dw) | 3.37 ± 1.40 | 1.31 ± 0.35 | 3.83 ± 2.30 | 2.34 ± 0.82 |
| **Fine roots** | | | | |
| RWC | 0.65 ± 0.02 | 0.65 ± 0.01 | 0.67 ± 0.01 | 0.64 ± 0.01 |
| $P_{tot}$ (mg $g^{-1}$ dw) | 1.22 ± 0.09 | 1.21 ± 0.11 | 1.44 ± 0.07 | 1.31 ± 0.08 |
| $P_{lab}$ (µg $g^{-1}$ dw) | 517.9 ± 11.1 | 598.1 ± 78.5 | 713.7 ± 19.8 | 662.7 ± 67.3 |
| N (mg $g^{-1}$ dw) | 14.0 ± 0.5 | 14.6 ± 0.3 | 14.6 ± 0.8 | 14.3 ± 0.5 |
| C (mg $g^{-1}$ dw) | 472.7 ± 2.8 | 463.1 ± 7.1 | 446.7 ± 10.2 | 460.8 ± 2.9 |
| C:N | 33.8 ± 1.2 | 31.7 ± 1.1 | 30.8 ± 1.6 | 32.3 ± 1.4 |
| N:P | 11.6 ± 1.1 | 12.2 ± 0.9 | 10.1 ± 0.2 | 10.9 ± 0.6 |

| **Fall** | | | | |
|---|---|---|---|---|
| **Organic layer** | **Con** | **N** | **P** | **P+N** |
| **Bulk soil** | | | | |
| RWC | 0.31 ± 0.05 | 0.37 ± 0.05 | 0.30 ± 0.01 | 0.36 ± 0.07 |
| $P_{tot}$ (mg $g^{-1}$ dw) | 1.51 ± 0.05 | 1.80 ± 0.19 | 1.66 ± 0.09 | 1.53 ± 0.29 |
| $P_{lab}$ (µg $g^{-1}$ dw) | 183.9 ± 32.2 | 180.3 ± 18.2 | 281.8 ± 12.5 | 216.2 ± 54.7 |
| N (mg $g^{-1}$ dw) | 12.9 ± 0.9 | 12.3 ± 1.9 | 13.8 ± 0.5 | 16.0 ± 4.3 |
| C (mg $g^{-1}$ dw) | 222.7 ± 18.8 | 178.6 ± 26.6 | 216.8 ± 3.2 | 265.4 ± 93.3 |
| C:N | 17.3 ± 0.8 | 14.6 ± 0.1 | 15.7 ± 0.3 | 15.9 ± 1.3 |
| N:P | 8.6 ± 0.6 | 7.1 ± 1.7 | 8.4 ± 0.4 | 9.6 ± 0.9 |
| pH | 4.00 ± 0.04 | 4.11 ± 0.07 | 4.10 ± 0.04 | 4.07 ± 0.18 |
| $NH_4^+$ (µmol $g^{-1}$ dw) | 272.4 ± 204.4 | 121.3 ± 61.8 | 108.3 ± 60.8 | 500.0 ± 225.0 |
| $NO_3^-$ (µmol $g^{-1}$ dw) | 68.3 ± 41.1 | 62.3 ± 27.2 | 42.2 ± 22.2 | 66.9 ± 38.5 |
| **Fine roots** | | | | |
| RWC | 0.61 ± 0.06 | 0.60 ± 0.02 | 0.61 ± 0.05 | 0.61 ± 0.01 |
| $P_{tot}$ (mg $g^{-1}$ dw) | 1.47 ± 0.19 | 1.10 ± 0.13 | 1.39 ± 0.10 | 1.32 ± 0.14 |
| $P_{lab}$ (µg $g^{-1}$ dw) | 581.5 ± 117.0 | 371.9 ± 48.1 | 633.2 ± 120.2 | 565.9 ± 113.6 |
| N (mg $g^{-1}$ dw) | 20.6 ± 0.9 | 17.0 ± 1.3 | 18.2 ± 0.6 | 19.3 ± 0.2 |
| C (mg $g^{-1}$ dw) | 448.2 ± 3.6 | 453.9 ± 7.8 | 438.9 ± 6.7 | 429.3 ± 24.9 |
| C:N | 21.9 ± 1.0 | 27.1 ± 2.4 | 24.2 ± 0.8 | 22.3 ± 1.1 |
| N:P | 14.4 ± 1.7 | 15.5 ± 0.6 | 13.2 ± 1.3 | 15.0 ± 1.9 |
| **Mineral topsoil** | | | | |
| **Bulk soil** | | | | |
| RWC | 0.27 ± 0.03 | 0.31 ± 0.01 | 0.32 ± 0.03 | 0.32 ± 0.01 |
| $P_{tot}$ (mg $g^{-1}$ dw) | 1.84 ± 0.21 | 2.14 ± 0.12 | 2.06 ± 0.20 | 2.09 ± 0.13 |
| $P_{lab}$ (µg $g^{-1}$ dw) | 109.0 ± 17.2 | 156.6 ± 32.4 | 180.9 ± 18.0 | 180.9 ± 34.4 |
| N (mg $g^{-1}$ dw) | 6.97 ± 0.55 | 7.95 ± 0.33 | 6.66 ± 0.43 | 8.81 ± 0.78 |
| C (mg $g^{-1}$ dw) | 93.9 ± 6.4 | 101.2 ± 3.1 | 91.5 ± 4.5 | 123.9 ± 19.3 |
| C:N | 13.5 ± 0.3 | 12.7 ± 0.2 | 13.8 ± 0.4 | 13.9 ± 0.9 |
| N:P | 3.9 ± 0.4 | 3.7 ± 0.1 | 3.3 ± 0.1 | 4.2 ± 0.3 |
| pH | 4.30 ± 0.02 | 4.32 ± 0.06 | 4.32 ± 0.02 | 4.27 ± 0.09 |
| $NH_4^+$ (µmol $g^{-1}$ dw) | 56.6 ± 18.4 | 38.8 ± 14.3 | 37.0 ± 13.0 | 48.4 ± 12.2 |
| $NO_3^-$ (µmol $g^{-1}$ dw) | 10.4 ± 1.7 | 24.0 ± 17.9 | 7.0 ± 1.4 | 42.0 ± 29.6 |
| **Fine roots** | | | | |
| RWC | 0.51 ± 0.06 | 0.60 ± 0.01 | 0.57 ± 0.01 | 0.59 ± 0.03 |
| $P_{tot}$ (mg $g^{-1}$ dw) | 1.15 ± 0.18 | 1.20 ± 0.10 | 1.19 ± 0.02 | 1.11 ± 0.05 |
| $P_{lab}$ (µg $g^{-1}$ dw) | 393.1 ± 94.4 | 348.9 ± 36.1 | 417.5 ± 37.9 | 459.6 ± 39.7 |
| N (mg $g^{-1}$ dw) | 18.0 ± 0.7 | 17.4 ± 0.3 | 17.0 ± 0.6 | 18.7 ± 1.4 |
| C (mg $g^{-1}$ dw) | 437.6 ± 12.7 | 434.7 ± 9.5 | 433.4 ± 0.4 | 436.4 ± 9.5 |
| C:N | 24.4 ± 1.5 | 25.1 ± 0.8 | 25.6 ± 0.9 | 23.5 ± 1.5 |
| N:P | 16.3 ± 1.9 | 14.6 ± 0.9 | 14.3 ± 0.6 | 16.8 ± 0.7 |

### Supplement Table S4 continued

**MP**

| Spring<br>Organic layer | Con | N | P | P+N |
|---|---|---|---|---|
| **Bulk soil** | | | | |
| RWC | 0.71 ± 0.02 | 0.71 ± 0.02 | 0.69 ± 0.01 | 0.72 ± 0.00 |
| $P_{tot}$ (mg $g^{-1}$ dw) | 1.29 ± 0.05 | 1.35 ± 0.09 | 1.41 ± 0.10 | 1.28 ± 0.09 |
| $P_{lab}$ (µg $g^{-1}$ dw) | 147.1 ± 13.5 | 127.4 ± 8.6 | 163.0 ± 7.6 | 173.5 ± 6.8 |
| N (mg $g^{-1}$ dw) | 23.1 ± 0.3 | 23.4 ± 0.4 | 22.8 ± 0.1 | 23.0 ± 0.2 |
| C (mg $g^{-1}$ dw) | 465.9 ± 2.9 | 471.3 ± 1.5 | 465.3 ± 4.3 | 470.7 ± 3.3 |
| C:N | 20.2 ± 0.2 | 20.1 ± 0.4 | 20.4 ± 0.2 | 20.5 ± 0.1 |
| N:P | 18.1 ± 0.9 | 17.5 ± 1.1 | 16.3 ± 1.1 | 18.1 ± 1.6 |
| pH | 3.72 ± 0.02 | 3.80 ± 0.02 | 3.76 ± 0.03 | 3.77 ± 0.03 |
| $NH_4^+$ (µmol $g^{-1}$ dw) | 144.7 ± 53.4 | 118.8 ± 21.7 | 191.0 ± 109.5 | 64.8 ± 22.1 |
| $NO_3^-$ (µmol $g^{-1}$ dw) | 4.87 ± 0.68 | 3.59 ± 1.54 | 4.55 ± 0.55 | 3.78 ± 0.58 |
| **Fine roots** | | | | |
| RWC | 0.70 ± 0.02 | 0.70 ± 0.01 | 0.71 ± 0.03 | 0.74 ± 0.01 |
| $P_{tot}$ (mg $g^{-1}$ dw) | 1.74 ± 0.04 | 1.37 ± 0.08 | 1.50 ± 0.06 | 1.65 ± 0.04 |
| $P_{lab}$ (µg $g^{-1}$ dw) | 555.1 ± 36.8 | 451.1 ± 28.4 | 696.6 ± 134.8 | 1020.2 ± 180.8 |
| N (mg $g^{-1}$ dw) | 17.7 ± 0.9 | 17.5 ± 1.3 | 17.9 ± 1.0 | 18.5 ± 1.6 |
| C (mg $g^{-1}$ dw) | 499.6 ± 2.0 | 495.0 ± 2.7 | 498.3 ± 4.3 | 496.3 ± 5.7 |
| C:N | 28.4 ± 1.5 | 28.7 ± 2.4 | 28.1 ± 2.0 | 27.4 ± 3.0 |
| N:P | 10.1 ± 0.4 | 12.8 ± 0.8 | 11.9 ± 0.3 | 11.2 ± 0.8 |
| **Mineral topsoil** | | | | |
| **Bulk soil** | | | | |
| RWC | 0.43 ± 0.02 | 0.42 ± 0.01 | 0.42 ± 0.01 | 0.42 ± 0.02 |
| $P_{tot}$ (mg $g^{-1}$ dw) | 1.58 ± 0.11 | 1.76 ± 0.07 | 1.47 ± 0.06 | 1.72 ± 0.05 |
| $P_{lab}$ (µg $g^{-1}$ dw) | 146.0 ± 27.3 | 136.6 ± 19.0 | 131.2 ± 23.4 | 152.2 ± 3.8 |
| N (mg $g^{-1}$ dw) | 6.86 ± 0.64 | 7.11 ± 0.19 | 6.17 ± 0.56 | 6.55 ± 0.38 |
| C (mg $g^{-1}$ dw) | 120.1 ± 11.4 | 121.2 ± 2.8 | 106.9 ± 11.1 | 112.5 ± 7.0 |
| C:N | 17.5 ± 0.2 | 17.0 ± 0.2 | 17.3 ± 0.2 | 17.2 ± 0.3 |
| N:P | 4.3 ± 0.3 | 4.0 ± 0.1 | 4.2 ± 0.2 | 3.8 ± 0.2 |
| pH | 3.92 ± 0.01 | 3.96 ± 0.05 | 3.97 ± 0.04 | 3.92 ± 0.02 |
| $NH_4^+$ (µmol $g^{-1}$ dw) | 13.3 ± 4.0 | 7.0 ± 1.7 | 41.9 ± 33.3 | 6.8 ± 2.6 |
| $NO_3^-$ (µmol $g^{-1}$ dw) | 1.64 ± 0.27 | 1.51 ± 0.10 | 1.71 ± 0.36 | 2.43 ± 0.21 |
| **Fine roots** | | | | |
| RWC | 0.39 ± 0.02 | 0.30 ± 0.05 | 0.32 ± 0.07 | 0.38 ± 0.02 |
| $P_{tot}$ (mg $g^{-1}$ dw) | 0.59 ± 0.02 | 1.09 ± 0.13 | 0.71 ± 0.08 | 1.00 ± 0.08 |
| $P_{lab}$ (µg $g^{-1}$ dw) | 297.1 ± 41.4 | 314.4 ± 9.5 | 359.9 ± 48.1 | 542.9 ± 45.9 |
| N (mg $g^{-1}$ dw) | 11.7 ± 0.1 | 14.4 ± 1.3 | 11.2 ± 0.4 | 12.7 ± 1.1 |
| C (mg $g^{-1}$ dw) | 485.9 ± 4.0 | 464.7 ± 15.4 | 484.9 ± 3.5 | 477.9 ± 10.0 |
| C:N | 41.4 ± 0.4 | 33.1 ± 4.3 | 43.4 ± 1.5 | 38.3 ± 3.7 |
| N:P | 19.9 ± 0.5 | 13.4 ± 1.7 | 15.9 ± 1.4 | 12.7 ± 0.4 |

| Fall<br>Organic layer | Con | N | P | P+N |
|---|---|---|---|---|
| **Bulk soil** | | | | |
| RWC | 0.51 ± 0.04 | 0.57 ± 0.06 | 0.56 ± 0.04 | 0.67 ± 0.06 |
| $P_{tot}$ (mg $g^{-1}$ dw) | 0.92 ± 0.01 | 0.89 ± 0.07 | 1.07 ± 0.06 | 0.97 ± 0.14 |
| $P_{lab}$ (µg $g^{-1}$ dw) | 98.3 ± 8.5 | 106.4 ± 2.3 | 149.4 ± 6.0 | 117.3 ± 10.7 |
| N (mg $g^{-1}$ dw) | 21.5 ± 1.4 | 20.4 ± 1.3 | 22.0 ± 1.7 | 23.6 ± 0.4 |
| C (mg $g^{-1}$ dw) | 428.0 ± 36.9 | 395.8 ± 36.0 | 408.5 ± 39.5 | 444.4 ± 24.4 |
| C:N | 19.9 ± 0.5 | 19.4 ± 0.5 | 18.6 ± 0.4 | 18.8 ± 0.7 |
| N:P | 23.3 ± 1.8 | 23.2 ± 2.6 | 20.9 ± 2.8 | 25.3 ± 3.6 |
| pH | 3.54 ± 0.03 | 3.69 ± 0.08 | 3.68 ± 0.03 | 3.61 ± 0.03 |
| $NH_4^+$ (µmol $g^{-1}$ dw) | 220.1 ± 25.1 | 107.8 ± 20.8 | 63.3 ± 21.8 | 73.4 ± 49.9 |
| $NO_3^-$ (µmol $g^{-1}$ dw) | 3.54 ± 1.07 | 4.04 ± 1.09 | 4.36 ± 1.04 | 4.83 ± 0.43 |
| **Fine roots** | | | | |
| RWC | 0.64 ± 0.01 | 0.64 ± 0.01 | 0.68 ± 0.01 | 0.67 ± 0.01 |
| $P_{tot}$ (mg $g^{-1}$ dw) | 0.96 ± 0.06 | 0.81 ± 0.02 | 1.11 ± 0.12 | 0.99 ± 0.12 |
| $P_{lab}$ (µg $g^{-1}$ dw) | 281.8 ± 23.3 | 251.3 ± 22.1 | 461.9 ± 39.2 | 385.2 ± 31.9 |
| N (mg $g^{-1}$ dw) | 22.5 ± 0.3 | 22.5 ± 0.4 | 23.9 ± 1.1 | 23.0 ± 0.5 |
| C (mg $g^{-1}$ dw) | 503.0 ± 2.3 | 504.9 ± 0.9 | 495.1 ± 3.4 | 506.0 ± 2.4 |
| C:N | 22.4 ± 0.4 | 22.4 ± 0.4 | 20.8 ± 1.1 | 22.0 ± 0.6 |
| N:P | 23.6 ± 1.1 | 27.9 ± 0.6 | 21.8 ± 1.4 | 23.9 ± 2.8 |
| **Mineral topsoil** | | | | |
| **Bulk soil** | | | | |
| RWC | 0.19 ± 0.08 | 0.34 ± 0.02 | 0.30 ± 0.01 | 0.38 ± 0.02 |
| $P_{tot}$ (mg $g^{-1}$ dw) | 1.07 ± 0.08 | 1.07 ± 0.10 | 0.87 ± 0.10 | 1.25 ± 0.02 |
| $P_{lab}$ (µg $g^{-1}$ dw) | 147.7 ± 34.5 | 97.1 ± 15.6 | 86.9 ± 5.8 | 174.3 ± 53.2 |
| N (mg $g^{-1}$ dw) | 6.80 ± 1.24 | 6.08 ± 0.49 | 4.99 ± 0.11 | 7.96 ± 0.62 |
| C (mg $g^{-1}$ dw) | 122.7 ± 23.2 | 99.7 ± 7.8 | 83.3 ± 3.5 | 131.4 ± 11.6 |
| C:N | 18.0 ± 0.3 | 16.4 ± 0.1 | 16.7 ± 0.4 | 16.5 ± 0.3 |
| N:P | 6.3 ± 0.7 | 5.7 ± 0.2 | 5.8 ± 0.5 | 6.3 ± 0.4 |
| pH | 3.90 ± 0.08 | 4.01 ± 0.01 | 4.07 ± 0.04 | 3.88 ± 0.04 |
| $NH_4^+$ (µmol $g^{-1}$ dw) | 24.9 ± 4.9 | 25.4 ± 10.6 | 39.2 ± 15.7 | 20.6 ± 4.4 |
| $NO_3^-$ (µmol $g^{-1}$ dw) | 10.07 ± 5.47 | 5.92 ± 1.57 | 5.50 ± 1.91 | 5.34 ± 0.50 |
| **Fine roots** | | | | |
| RWC | 0.42 ± 0.00 | 0.42 ± 0.01 | 0.41 ± 0.02 | 0.41 ± 0.02 |
| $P_{tot}$ (mg $g^{-1}$ dw) | 0.78 ± 0.06 | 0.69 ± 0.07 | 0.66 ± 0.01 | 0.84 ± 0.11 |
| $P_{lab}$ (µg $g^{-1}$ dw) | 272.1 ± 46.0 | 199.3 ± 26.4 | 216.7 ± 6.2 | 361.3 ± 39.4 |
| N (mg $g^{-1}$ dw) | 11.5 ± 0.6 | 11.1 ± 0.6 | 10.5 ± 0.9 | 11.3 ± 0.3 |
| C (mg $g^{-1}$ dw) | 462.5 ± 7.7 | 454.8 ± 13.2 | 451.0 ± 21.7 | 459.1 ± 12.6 |
| C:N | 40.4 ± 1.5 | 41.3 ± 3.3 | 43.3 ± 3.0 | 40.6 ± 0.4 |
| N:P | 14.8 ± 0.6 | 16.3 ± 1.1 | 15.9 ± 1.3 | 13.9 ± 2.1 |

### Supplement Table S4 continued

**LP**

| **Spring** | | | | |
|---|---|---|---|---|
| **organic layer** | **Con** | **N** | **P** | **P+N** |
| **Bulk soil** | | | | |
| RWC | 0.70 ± 0.02 | 0.65 ± 0.02 | 0.69 ± 0.03 | 0.66 ± 0.03 |
| $P_{tot}$ (mg g$^{-1}$ dw) | 0.62 ± 0.07 | 0.57 ± 0.06 | 0.72 ± 0.06 | 0.75 ± 0.05 |
| $P_{lab}$ (µg g$^{-1}$ dw) | 124.5 ± 24.7 | 119.4 ± 19.1 | 153.4 ± 25.3 | 190.2 ± 33.7 |
| N (mg g$^{-1}$ dw) | 16.9 ± 2.3 | 14.7 ± 1.6 | 17.0 ± 2.1 | 16.4 ± 1.6 |
| C (mg g$^{-1}$ dw) | 358.1 ± 32.3 | 315.7 ± 26.6 | 368.5 ± 44.2 | 331.3 ± 33.1 |
| C:N | 21.4 ± 0.9 | 21.6 ± 0.7 | 21.7 ± 0.9 | 20.2 ± 0.3 |
| N:P | 27.2 ± 1.2 | 25.8 ± 0.7 | 23.5 ± 1.1 | 21.8 ± 0.8 |
| pH | 4.38 ± 0.10 | 4.39 ± 0.08 | 4.34 ± 0.15 | 4.55 ± 0.03 |
| $NH_4^+$ (µmol g$^{-1}$ dw) | 278.1 ± 265.4 | 539.2 ± 122.3 | 23.5 ± 16.8 | 77.5 ± 66.7 |
| $NO_3^-$ (µmol g$^{-1}$ dw) | 42.5 ± 18.1 | 11.0 ± 2.4 | 12.7 ± 2.4 | 11.9 ± 2.4 |
| **Fine roots** | | | | |
| RWC | 0.69 ± 0.01 | 0.72 ± 0.00 | 0.70 ± 0.02 | 0.71 ± 0.02 |
| $P_{tot}$ (mg g$^{-1}$ dw) | 1.08 ± 0.14 | 1.15 ± 0.11 | 1.56 ± 0.12 | 1.31 ± 0.13 |
| $P_{lab}$ (µg g$^{-1}$ dw) | 596.9 ± 167.7 | 646.4 ± 92.5 | 953.8 ± 136.5 | 764.4 ± 224.4 |
| N (mg g$^{-1}$ dw) | 19.2 ± 2.7 | 20.3 ± 1.3 | 19.8 ± 1.7 | 20.9 ± 2.9 |
| C (mg g$^{-1}$ dw) | 491.2 ± 3.4 | 498.9 ± 3.8 | 495.1 ± 1.6 | 489.3 ± 7.1 |
| C:N | 26.6 ± 3.8 | 24.8 ± 1.7 | 25.4 ± 2.2 | 24.5 ± 4.0 |
| N:P | 17.7 ± 0.8 | 17.8 ± 0.9 | 12.7 ± 1.1 | 15.8 ± 0.9 |
| **Mineral topsoil** | | | | |
| **Bulk soil** | | | | |
| RWC | 0.18 ± 0.02 | 0.17 ± 0.01 | 0.21 ± 0.01 | 0.18 ± 0.01 |
| $P_{tot}$ (mg g$^{-1}$ dw) | 0.10 ± 0.01 | 0.10 ± 0.02 | 0.10 ± 0.01 | 0.10 ± 0.01 |
| $P_{lab}$ (µg g$^{-1}$ dw) | 23.5 ± 3.8 | 27.0 ± 5.3 | 38.0 ± 3.3 | 32.3 ± 4.7 |
| N (mg g$^{-1}$ dw) | 1.19 ± 0.14 | 1.40 ± 0.10 | 1.80 ± 0.27 | 1.38 ± 0.18 |
| C (mg g$^{-1}$ dw) | 26.2 ± 2.8 | 32.8 ± 2.2 | 41.8 ± 5.0 | 30.3 ± 3.0 |
| C:N | 22.1 ± 0.4 | 23.5 ± 1.0 | 23.5 ± 0.9 | 22.1 ± 0.7 |
| N:P | 12.2 ± 0.7 | 15.5 ± 3.1 | 17.0 ± 0.7 | 14.2 ± 2.3 |
| pH | 4.25 ± 0.06 | 4.41 ± 0.13 | 4.16 ± 0.05 | 4.35 ± 0.14 |
| $NH_4^+$ (µmol g$^{-1}$ dw) | 126.7 ± 111.8 | 15.4 ± 19.5 | 1.8 ± 1.1 | 27.3 ± 25.0 |
| $NO_3^-$ (µmol g$^{-1}$ dw) | 4.73 ± 0.62 | 7.20 ± 1.91 | 22.7 ± 16.6 | 5.08 ± 0.25 |
| **Fine roots** | | | | |
| RWC | 0.42 ± 0.01 | 0.43 ± 0.02 | 0.42 ± 0.01 | 0.42 ± 0.02 |
| $P_{tot}$ (mg g$^{-1}$ dw) | 0.55 ± 0.04 | 0.55 ± 0.07 | 0.73 ± 0.04 | 0.89 ± 0.10 |
| $P_{lab}$ (µg g$^{-1}$ dw) | 202.1 ± 8.9 | 218.5 ± 35.1 | 363.2 ± 19.8 | 310.1 ± 48.7 |
| N (mg g$^{-1}$ dw) | 12.4 ± 1.6 | 11.9 ± 1.2 | 12.0 ± 1.4 | 11.6 ± 1.0 |
| C (mg g$^{-1}$ dw) | 467.1 ± 20.8 | 466.5 ± 15.5 | 487.6 ± 12.0 | 459.9 ± 31.5 |
| C:N | 38.7 ± 4.0 | 39.8 ± 3.1 | 41.7 ± 4.4 | 39.6 ± 0.8 |
| N:P | 22.4 ± 1.6 | 21.9 ± 0.6 | 16.4 ± 1.2 | 13.3 ± 1.8 |

| **Fall** | | | | |
|---|---|---|---|---|
| **organic layer** | **Con** | **N** | **P** | **P+N** |
| **Bulk soil** | | | | |
| RWC | 0.25 ± 0.02 | 0.20 ± 0.06 | 0.27 ± 0.05 | 0.27 ± 0.01 |
| $P_{tot}$ (mg g$^{-1}$ dw) | 0.42 ± 0.03 | 0.40 ± 0.08 | 0.43 ± 0.13 | 0.49 ± 0.03 |
| $P_{lab}$ (µg g$^{-1}$ dw) | 47.2 ± 5.5 | 55.4 ± 10.4 | 67.9 ± 16.0 | 63.5 ± 8.0 |
| N (mg g$^{-1}$ dw) | 11.8 ± 1.4 | 11.2 ± 3.1 | 10.9 ± 1.1 | 15.4 ± 1.6 |
| C (mg g$^{-1}$ dw) | 242.5 ± 31.9 | 239.2 ± 61.9 | 228.6 ± 19.4 | 321.4 ± 37.4 |
| C:N | 20.4 ± 0.5 | 21.6 ± 0.8 | 21.1 ± 0.3 | 20.8 ± 0.8 |
| N:P | 28.3 ± 1.4 | 27.3 ± 2.4 | 29.1 ± 7.7 | 31.4 ± 3.3 |
| pH | 3.90 ± 0.05 | 3.98 ± 0.11 | 3.82 ± 0.11 | 3.99 ± 0.05 |
| $NH_4^+$ (µmol g$^{-1}$ dw) | 53.8 ± 47.4 | 467.7 ± 212.7 | 75.9 ± 67.9 | 261.5 ± 216.0 |
| $NO_3^-$ (µmol g$^{-1}$ dw) | 15.6 ± 3.8 | 63.8 ± 50.7 | 41.1 ± 28.1 | 92.8 ± 41.9 |
| **Fine roots** | | | | |
| RWC | 0.60 ± 0.02 | 0.54 ± 0.00 | 0.56 ± 0.01 | 0.57 ± 0.03 |
| $P_{tot}$ (mg g$^{-1}$ dw) | 0.99 ± 0.32 | 0.66 ± 0.04 | 0.81 ± 0.11 | 0.91 ± 0.13 |
| $P_{lab}$ (µg g$^{-1}$ dw) | 192.6 ± 10.9 | 198.5 ± 24.6 | 229.4 ± 34.3 | 214.6 ± 5.5 |
| N (mg g$^{-1}$ dw) | 15.0 ± 0.2 | 14.1 ± 2.9 | 14.1 ± 1.9 | 15.3 ± 0.1 |
| C (mg g$^{-1}$ dw) | 508.2 ± 0.9 | 504.7 ± 7.1 | 515.0 ± 8.3 | 511.0 ± 4.5 |
| C:N | 34.0 ± 0.5 | 38.6 ± 7.3 | 38.0 ± 5.6 | 33.5 ± 0.5 |
| N:P | 18.2 ± 4.7 | 21.2 ± 3.6 | 17.6 ± 1.4 | 17.5 ± 2.3 |
| **Mineral topsoil** | | | | |
| **Bulk soil** | | | | |
| RWC | 0.04 ± 0.00 | 0.04 ± 0.01 | 0.07 ± 0.02 | 0.05 ± 0.01 |
| $P_{tot}$ (mg g$^{-1}$ dw) | 0.08 ± 0.01 | 0.08 ± 0.02 | 0.12 ± 0.02 | 0.09 ± 0.02 |
| $P_{lab}$ (µg g$^{-1}$ dw) | 14.6 ± 2.0 | 14.5 ± 4.0 | 27.1 ± 3.1 | 26.0 ± 11.3 |
| N (mg g$^{-1}$ dw) | 0.94 ± 0.13 | 1.07 ± 0.48 | 2.00 ± 0.84 | 1.46 ± 0.67 |
| C (mg g$^{-1}$ dw) | 26.2 ± 2.7 | 30.3 ± 11.3 | 51.1 ± 17.7 | 38.5 ± 15.5 |
| C:N | 24.6 ± 0.7 | 27.1 ± 0.6 | 29.4 ± 0.5 | 33.6 ± 1.8 |
| N:P | 12.8 ± 3.1 | 12.8 ± 4.9 | 15.9 ± 4.3 | 16.2 ± 7.5 |
| pH | 4.26 ± 0.06 | 4.24 ± 0.08 | 4.10 ± 0.07 | 4.20 ± 0.04 |
| $NH_4^+$ (µmol g$^{-1}$ dw) | 51.9 ± 20.1 | 94.6 ± 26.5 | 21.8 ± 6.5 | 62.1 ± 40.1 |
| $NO_3^-$ (µmol g$^{-1}$ dw) | 2.52 ± 0.72 | 3.64 ± 1.96 | 3.15 ± 0.37 | 3.58 ± 1.04 |
| **Fine roots** | | | | |
| RWC | 0.51 ± 0.02 | 0.53 ± 0.01 | 0.53 ± 0.02 | 0.57 ± 0.01 |
| $P_{tot}$ (mg g$^{-1}$ dw) | 0.55 ± 0.03 | 0.59 ± 0.08 | 0.53 ± 0.04 | 0.72 ± 0.11 |
| $P_{lab}$ (µg g$^{-1}$ dw) | 114.0 ± 17.2 | 129.6 ± 35.9 | 171.4 ± 18.0 | 235.7 ± 69.9 |
| N (mg g$^{-1}$ dw) | 11.0 ± 0.3 | 12.8 ± 0.9 | 10.4 ± 0.3 | 13.8 ± 1.5 |
| C (mg g$^{-1}$ dw) | 487.2 ± 3.9 | 475.9 ± 7.8 | 481.4 ± 11.0 | 467.8 ± 7.6 |
| C:N | 44.3 ± 1.4 | 37.7 ± 3.0 | 46.3 ± 2.1 | 34.6 ± 3.1 |
| N:P | 19.9 ± 0.9 | 20.9 ± 1.2 | 19.9 ± 1.2 | 19.5 ± 1.8 |

**Supplement Table 5: Species richness, Shannon diversity and evenness of ectomycorrhizal root-colonizing fungi (EMF), root-associated fungi (RAF) and soil-residing fungi (SAF) of beech forests (*Fagus sylvatica* L.) in response to fertilization treatments (Con, N, P, P+N).** Soil and root samples were collected in a P-rich (HP), P-medium (MP) and P-poor (LP) forest in spring and fall 2018. The organic layer and mineral topsoil were analyzed separately. Data indicate means ± SE (RAF and SAF: n = 3, EMF: 3 samples per plot were added to reach species saturation). NA = not available

| | **Species richness** | | | | | | | | | |
|---|---|---|---|---|---|---|---|---|---|---|
| | **Spring** | | | | | **Fall** | | | | |
| | **Organic layer** | **Con** | **N** | **P** | **P+N** | **Organic layer** | **Con** | **N** | **P** | **P+N** |
| | EMF | NA | | | | EMF | 7 | 7 | 9 | 6 |
| **HP** | RAF | 177 ± 23.7 | 234 ± 14 | 242 ± 34.9 | 269 ± 39.8 | RAF | 357 ± 11.9 | 323 ± 2.3 | 331 ± 14.3 | 325 ± 6.8 |
| | SAF | 627 ± 9.1 | 597 ± 18 | 711 ± 70.3 | 776 ± 13.5 | SAF | 929 ± 10.4 | 861 ± 14.2 | 912 ± 7.3 | 931 ± 4.6 |
| | EMF | NA | | | | EMF | 4 | 8 | 7 | 6 |
| **MP** | RAF | 211 ± 6.4 | 202 ± 11.5 | 228 ± 10.9 | 173 ± 10.4 | RAF | 231 ± 20.6 | 221 ± 11.3 | 237 ± 0.6 | 261 ± 8 |
| | SAF | 787 ± 11.5 | 782 ± 28.9 | 770 ± 10.6 | 798 ± 20.5 | SAF | 841 ± 11.6 | 805 ± 31.9 | 784 ± 9.2 | 823 ± 13.9 |
| | EMF | NA | | | | EMF | 14 | 11 | 12 | 13 |
| **LP** | RAF | 230 ± 14.4 | 205 ± 15.5 | 244 ± 24.8 | 282 ± 20.4 | RAF | 257 ± 7.1 | 240 ± 5.4 | 261 ± 9.3 | 243 ± 44.5 |
| | SAF | 629 ± 46.9 | 583 ± 2.4 | 682 ± 12.1 | 620 ± 0.9 | SAF | 839 ± 34.9 | 872 ± 22.5 | 908 ± 31.6 | 897 ± 26.6 |
| | **Mineral topsoil** | | | | | **Mineral topsoil** | | | | |
| | EMF | NA | | | | EMF | 8 | 10 | 7 | 12 |
| **HP** | RAF | 171 ± 3.3 | 202 ± 5.5 | 183 ± 12.8 | 187 ± 13.4 | RAF | 188 ± 22 | 179 ± 21 | 194 ± 17.3 | 159 ± 22.2 |
| | SAF | 738 ± 20.2 | 816 ± 9.9 | 787 ± 23.2 | 774 ± 31.2 | SAF | 930 ± 2.9 | 908 ± 17.6 | 886 ± 10.6 | 933 ± 18.9 |
| | EMF | NA | | | | EMF | 6 | 9 | 7 | 4 |
| **MP** | RAF | 158 ± 3.8 | 146 ± 7.5 | 159 ± 16.9 | 203 ± 13.1 | RAF | 234 ± 47 | 230 ± 11.9 | 204 ± 11.9 | 234 ± 15.6 |
| | SAF | 771 ± 24.2 | 759 ± 7.9 | 756 ± 16.1 | 756 ± 9.9 | SAF | 734 ± 52.7 | 796 ± 10.4 | 744 ± 11.3 | 745 ± 25.9 |
| | EMF | NA | | | | EMF | 5 | 7 | 5 | 9 |
| **LP** | RAF | 216 ± 18.2 | 180 ± 20.2 | 188 ± 22.7 | 196 ± 4.6 | RAF | 199 ± 7.5 | 181 ± 21.3 | 176 ± 10.8 | 179 ± 8.1 |
| | SAF | 459 ± 10.6 | 390 ± 35.6 | 450 ± 44.7 | 483 ± 16.3 | SAF | 800 ± 14.5 | 806 ± 13.9 | 862 ± 9.9 | 821 ± 9 |

### Supplement Table S5 continued

| | **Shannon diversity**<br>**Spring**<br>**Organic layer** | **Con** | **N** | **P** | **P+N** | **Fall**<br>**Organic layer** | **Con** | **N** | **P** | **P+N** |
|---|---|---|---|---|---|---|---|---|---|---|
| | EMF | | NA | | | EMF | 1.29 | 0.72 | 1.41 | 1.45 |
| **HP** | RAF | 1.99 ± 0.26 | 3.09 ± 0.18 | 3.11 ± 0.27 | 3.41 ± 0.22 | RAF | 3.9 ± 0.04 | 3.7 ± 0.03 | 3.72 ± 0.14 | 3.76 ± 0.07 |
| | SAF | 4 ± 0.23 | 3.87 ± 0.12 | 4.68 ± 0.21 | 4.68 ± 0.01 | SAF | 5.17 ± 0.03 | 5.03 ± 0.04 | 5.13 ± 0 | 5.19 ± 0.01 |
| | EMF | | NA | | | EMF | 1.17 | 1.62 | 0.64 | 1.45 |
| **MP** | RAF | 2.96 ± 0.11 | 2.77 ± 0.12 | 2.86 ± 0.08 | 2.53 ± 0.1 | RAF | 3.02 ± 0.14 | 2.91 ± 0.1 | 2.77 ± 0.05 | 2.97 ± 0.04 |
| | SAF | 4.67 ± 0.04 | 4.73 ± 0.18 | 4.57 ± 0.05 | 4.73 ± 0.06 | SAF | 4.88 ± 0.03 | 4.7 ± 0.14 | 4.62 ± 0.12 | 4.62 ± 0.05 |
| | EMF | | NA | | | EMF | 1.38 | 1.94 | 1.41 | 1.53 |
| **LP** | RAF | 3.03 ± 0.08 | 2.67 ± 0.33 | 2.97 ± 0.4 | 3.15 ± 0.04 | RAF | 3.18 ± 0.22 | 3.09 ± 0.21 | 2.97 ± 0.21 | 2.56 ± 0.55 |
| | SAF | 4.61 ± 0.16 | 4.39 ± 0.05 | 4.53 ± 0.22 | 4.4 ± 0.13 | SAF | 4.98 ± 0.13 | 5.15 ± 0.02 | 5.21 ± 0.08 | 5.12 ± 0.07 |
| | **Mineral topsoil** | | | | | **Mineral topsoil** | | | | |
| | EMF | | NA | | | EMF | 1.91 | 2.04 | 1.86 | 1.66 |
| **HP** | RAF | 2.97 ± 0.17 | 3.09 ± 0.09 | 3.21 ± 0.03 | 3.23 ± 0.19 | RAF | 2.78 ± 0.12 | 2.7 ± 0.16 | 2.78 ± 0.14 | 2.56 ± 0.27 |
| | SAF | 4.85 ± 0.07 | 4.89 ± 0.06 | 4.86 ± 0.03 | 4.89 ± 0.03 | SAF | 5.17 ± 0 | 5.12 ± 0.04 | 5.08 ± 0.03 | 5.17 ± 0.03 |
| | EMF | | NA | | | EMF | 1.75 | 1.55 | 1.4 | 0.95 |
| **MP** | RAF | 2.81 ± 0.08 | 2.61 ± 0.34 | 2.67 ± 0.06 | 3.19 ± 0.11 | RAF | 2.87 ± 0.4 | 3.15 ± 0.04 | 2.89 ± 0.14 | 3.23 ± 0.16 |
| | SAF | 4.64 ± 0.04 | 4.71 ± 0.06 | 4.74 ± 0.1 | 4.81 ± 0.08 | SAF | 4.12 ± 0.23 | 4.58 ± 0.17 | 4.33 ± 0.06 | 4.44 ± 0.17 |
| | EMF | | NA | | | EMF | 0.62 | 1.32 | 1.72 | 1.37 |
| **LP** | RAF | 2.95 ± 0.11 | 2.77 ± 0.23 | 2.58 ± 0.36 | 3.11 ± 0.13 | RAF | 2.78 ± 0.12 | 2.59 ± 0.51 | 2.39 ± 0.07 | 2.81 ± 0.09 |
| | SAF | 3.97 ± 0.08 | 3.9 ± 0.1 | 4.06 ± 0.08 | 3.89 ± 0.08 | SAF | 4.9 ± 0.06 | 4.77 ± 0.04 | 5.01 ± 0.03 | 4.76 ± 0.15 |

**Supplement Table S5 continued**

| | **Evenness** | | | | | | | | | |
|---|---|---|---|---|---|---|---|---|---|---|
| | **Spring** | | | | | **Fall** | | | | |
| | **Organic layer** | **Con** | **N** | **P** | **P+N** | **Organic layer** | **Con** | **N** | **P** | **P+N** |
| | EMF | | NA | | | EMF | 0.66 | 0.37 | 0.79 | 0.66 |
| **HP** | RAF | 0.38 ± 0.04 | 0.57 ± 0.03 | 0.57 ± 0.04 | 0.61 ± 0.03 | RAF | 0.66 ± 0 | 0.64 ± 0 | 0.64 ± 0.02 | 0.65 ± 0.01 |
| | SAF | 0.62 ± 0.03 | 0.61 ± 0.02 | 0.71 ± 0.02 | 0.7 ± 0 | SAF | 0.76 ± 0 | 0.74 ± 0 | 0.75 ± 0 | 0.76 ± 0 |
| | EMF | | NA | | | EMF | 0.73 | 0.7 | 0.31 | 0.59 |
| **MP** | RAF | 0.55 ± 0.02 | 0.52 ± 0.02 | 0.53 ± 0.01 | 0.49 ± 0.01 | RAF | 0.56 ± 0.02 | 0.54 ± 0.02 | 0.51 ± 0.01 | 0.53 ± 0 |
| | SAF | 0.7 ± 0.01 | 0.71 ± 0.02 | 0.69 ± 0.01 | 0.71 ± 0.01 | SAF | 0.72 ± 0 | 0.7 ± 0.02 | 0.69 ± 0.02 | 0.69 ± 0.01 |
| | EMF | | NA | | | EMF | 0.52 | 0.81 | 0.55 | 0.79 |
| **LP** | RAF | 0.56 ± 0.01 | 0.5 ± 0.05 | 0.54 ± 0.06 | 0.56 ± 0.01 | RAF | 0.57 ± 0.04 | 0.56 ± 0.04 | 0.53 ± 0.04 | 0.46 ± 0.09 |
| | SAF | 0.72 ± 0.02 | 0.69 ± 0.01 | 0.69 ± 0.03 | 0.68 ± 0.02 | SAF | 0.74 ± 0.01 | 0.76 ± 0 | 0.76 ± 0.01 | 0.75 ± 0.01 |
| | **Mineral topsoil** | | | | | **Mineral topsoil** | | | | |
| | EMF | | NA | | | EMF | 0.87 | 0.82 | 0.78 | 0.75 |
| **HP** | RAF | 0.58 ± 0.03 | 0.58 ± 0.01 | 0.62 ± 0.01 | 0.62 ± 0.03 | RAF | 0.53 ± 0.03 | 0.52 ± 0.02 | 0.53 ± 0.02 | 0.51 ± 0.04 |
| | SAF | 0.73 ± 0.01 | 0.73 ± 0.01 | 0.73 ± 0 | 0.74 ± 0 | SAF | 0.76 ± 0 | 0.75 ± 0.01 | 0.75 ± 0 | 0.76 ± 0 |
| | EMF | | NA | | | EMF | 0.9 | 0.67 | 0.87 | 0.49 |
| **MP** | RAF | 0.55 ± 0.02 | 0.52 ± 0.06 | 0.53 ± 0 | 0.6 ± 0.02 | RAF | 0.53 ± 0.05 | 0.58 ± 0 | 0.54 ± 0.02 | 0.59 ± 0.02 |
| | SAF | 0.7 ± 0 | 0.71 ± 0.01 | 0.71 ± 0.01 | 0.73 ± 0.01 | SAF | 0.62 ± 0.03 | 0.69 ± 0.02 | 0.65 ± 0.01 | 0.67 ± 0.02 |
| | EMF | | NA | | | EMF | 0.39 | 0.68 | 0.78 | 0.85 |
| **LP** | RAF | 0.55 ± 0.01 | 0.53 ± 0.04 | 0.49 ± 0.06 | 0.59 ± 0.02 | RAF | 0.53 ± 0.02 | 0.5 ± 0.09 | 0.46 ± 0.02 | 0.54 ± 0.01 |
| | SAF | 0.65 ± 0.01 | 0.66 ± 0.02 | 0.67 ± 0 | 0.63 ± 0.01 | SAF | 0.73 ± 0.01 | 0.71 ± 0.01 | 0.74 ± 0 | 0.71 ± 0.02 |

**Supplement Table S6: Mortality (%) of beech fine roots (*Fagus sylvatica* L.) in different forests, soil layers and in response to fertilization treatments (Con, N, P, P+N).** Fine root samples were collected in a P-rich (HP), P-medium (MP) and P-poor (LP) forest in spring and fall 2018. The organic layer (OL) and mineral topsoil (ML) were analyzed separately; data for season were analyzed together. Data indicate means (n = 6 ± SE). Differences of means were tested by a linear mixed model and Tukey HSD posthoc test for the factors layer and treatment with plot number as random effect. We tested the effects of layer and treatment for each site. Bold letters indicate significant differences at p ≤ 0.05.

| Forest | Layer | Con | N | P | P+N | Treatment | | Layer | |
|---|---|---|---|---|---|---|---|---|---|
| | | | | | | F | *p* | F | *p* |
| HP | OL | 30.5 ±3.2 | 30.6 ±2.0 | 42.0 ±10.4 | 35.3 ±6.6 | 0.7 | 0.573 | 0.5 | 0.478 |
| | ML | 32.8 ±6.4 | 37.7 ±5.5 | 31.4 ±7.9 | 23.6 ±4.0 | 0.9 | 0.481 | | |
| MP | OL | 17.4 ±3.8 | 30.4 ±5.9 | 18.4 ±2.2 | 19.2 ±1.1 | 2.7 | 0.116 | 4.7 | **0.047** |
| | ML | 35.1 ±7.7 | 29.0 ±3.9 | 23.7 ±3.2 | 25.0 ±4.6 | 0.9 | 0.448 | | |
| LP | OL | 29.3 ±10.0 | 10.9 ±3.9 | 23.5 ±3.4 | 21.0 ±4.1 | 1.6 | 0.256 | 13.7 | **0.003** |
| | ML | 52.6 ±6.8 | 42.1 ±9.3 | 40.5 ±4.1 | 57.1 ±5.4 | 0.3 | 0.809 | | |

**Supplement Table S7: Relative abundance of all fungal orders obtained by Ilumina MiSeq in soil and associated with roots in beech forests (*Fagus sylvatica* L.).** Soil and fine root samples of the organic and mineral layer were collected in a P-rich, P-medium and P-poor forest in 2018. All data were pooled the abundance of an order was expressed relative to the total number of sequences.

| **> 1%** | | **0.1% - 1%** | |
|---|---|---|---|
| **abundance (%)** | **Order** | **abundance (%)** | **Order** |
| 27.34 | Agaricales | 0.75 | Sebacinales |
| 16.99 | Helotiales | 0.70 | Rhytismatales |
| 13.60 | Russulales | 0.69 | Tremellales |
| 7.36 | Boletales | 0.50 | Chaetosphaeriales |
| 3.91 | Mytilinidiales | 0.47 | Chaetothyriales |
| 3.36 | Atheliales | 0.43 | Incertae |
| 3.21 | Eurotiales | 0.36 | Auriculariales |
| 3.00 | Cantharellales | 0.28 | Archaeorhizomycetales |
| 2.89 | Thelephorales | 0.24 | Polyporales |
| 2.87 | Hypocreales | 0.22 | Trichosporonales |
| 2.40 | Mortierellales | 0.17 | Hymenochaetales |
| 2.19 | Trechisporales | 0.16 | Capnodiales |
| 2.07 | Pleosporales | 0.15 | Xylariales |
| 1.47 | Sordariales | 0.15 | Thelebolales |
| 1.06 | Pezizales | 0.13 | Agaricomycetes |
| | | 0.11 | Venturiales |

| **< 0.1 %** | | **< 0.01 %** | | **< 0.003 %** | |
|---|---|---|---|---|---|
| **abundance (%)** | **Order** | **abundance (%)** | **Order** | **abundance (%)** | **Order** |
| 0.082 | Togniniales | 0.0099 | Diaporthales | 0.0029 | Magnaporthales |
| 0.051 | Umbelopsidales | 0.0096 | Atractiellales | 0.0027 | Verrucariales |
| 0.050 | Phacidiales | 0.0094 | Geastrales | 0.0025 | Jaapiales |
| 0.049 | Leucosporidiales | 0.0092 | Microascales | 0.0024 | Lecanoromycetes |
| 0.045 | Zoopagales | 0.0091 | Tubeufiales | 0.0022 | Myrmecridiales |
| 0.045 | Coniochaetales | 0.0089 | Microbotryomycetes | 0.0019 | Onygenales |
| 0.037 | Mucorales | 0.0085 | Spizellomycetales | 0.0017 | Diversisporales |
| 0.036 | Phallales | 0.0083 | Erythrobasidiales | 0.0015 | Kriegeriales |
| 0.036 | Saccharomycetales | 0.0081 | Rhizophydiales | 0.0015 | Botryosphaeriales |
| 0.031 | Sporidiobolales | 0.0070 | Filobasidiales | 0.0014 | Georgefischeriales |
| 0.030 | Orbiliales | 0.0062 | Ustilaginales | 0.0013 | Cystofilobasidiales |
| 0.029 | Glomerales | 0.0062 | Lecanorales | 0.0012 | Exobasidiales |
| 0.029 | Glomerellales | 0.0059 | Diaporthales | 0.0010 | Myriangiales |
| 0.024 | Basidiobolales | 0.0055 | Malasseziales | 0.0009 | Umbilicariales |
| 0.022 | Pyxidiophorales | 0.0047 | Taphrinales | 0.0007 | Calosphaeriales |
| 0.018 | Dothideales | 0.0045 | Ophiostomatales | 0.0007 | Lobulomycetales |
| 0.013 | Hysterangiales | 0.0042 | Dothideomycetes | 0.0007 | Urocystidales |
| 0.011 | Corticiales | 0.0037 | Geoglossales | 0.0005 | Agaricostilbales |
| 0.010 | Ostropales | 0.0035 | Acarosporales | 0.0004 | Phomatosporales |
| | | | | 0.0003 | Cystobasidiomycetes |
| | | | | 0.0003 | Phaeomoniellales |
| | | | | 0.0003 | Dacrymycetales |
| | | | | 0.0003 | Gloeophyllales |
| | | | | 0.0001 | Erysiphales |

**Supplement Table S8: Relative abundances (%) of fungal orders and trophic guilds in three beech forests (*Fagus sylvatica* L.) in response to fertilization treatments (Con, N, P, P+N).** Soil and root samples were collected in a P-rich (HP), P-medium (MP) and P-poor (LP) forest in spring and fall 2018. The organic layer and mineral topsoil were analyzed separately. Data indicate means (n = 3 ± SE). Statistical information is provided in Table S9.

**HP**

| **Spring** | | | | |
|---|---|---|---|---|
| **Organic layer** | **Con** | **N** | **P** | **P+N** |
| **SAF - Bulk soil** | | | | |
| Agaricales | 12.4 ± 4.9 | 18.6 ± 4.6 | 17.6 ± 4.5 | 26.8 ± 13.2 |
| Atheliales | 0.0 ± 0.0 | 0.1 ± 0.0 | 0.2 ± 0.1 | 1.9 ± 1.5 |
| Boletales | 2.0 ± 1.2 | 0.7 ± 0.2 | 1.8 ± 0.4 | 1.3 ± 0.6 |
| Cantharellales | 7.1 ± 2.4 | 5.3 ± 0.5 | 2.3 ± 1.2 | 6.3 ± 4.6 |
| Russulales | 7.9 ± 3.4 | 4.0 ± 1.2 | 6.1 ± 1.4 | 3.6 ± 1.4 |
| Thelephorales | 1.6 ± 0.6 | 4.1 ± 0.9 | 3.6 ± 1.8 | 2.2 ± 0.4 |
| Trechisporales | 0.6 ± 0.3 | 5.0 ± 3.9 | 1.2 ± 0.8 | 0.7 ± 0.3 |
| Eurotiales | 0.6 ± 0.4 | 0.3 ± 0.1 | 0.4 ± 0.2 | 0.6 ± 0.3 |
| Helotiales | 22.5 ± 8.2 | 14.1 ± 0.9 | 22.1 ± 6.8 | 11.7 ± 1.3 |
| Hypocreales | 5.5 ± 1.5 | 4.5 ± 0.5 | 7.0 ± 1.4 | 4.3 ± 0.1 |
| Mytilinidiales | 0.1 ± 0.0 | 0.9 ± 0.9 | 0.1 ± 0.1 | 0.7 ± 0.4 |
| Pezizales | 0.2 ± 0.1 | 0.9 ± 0.3 | 0.5 ± 0.1 | 0.6 ± 0.1 |
| Pleosporales | 18.9 ± 6.9 | 16.3 ± 2.6 | 8.5 ± 4.5 | 19.1 ± 7.1 |
| Sordariales | 5.0 ± 1.6 | 5.8 ± 1.2 | 10.0 ± 6.3 | 3.6 ± 0.1 |
| Mortirellales | 1.7 ± 0.4 | 2.3 ± 0.4 | 3.4 ± 0.6 | 2.7 ± 0.2 |
| Symbiotroph | 17.7 ± 6.7 | 69.3 ± 2.6 | 47.8 ± 11.9 | 75.2 ± 12.3 |
| Saprotroph | 40.5 ± 6.5 | 7.9 ± 3.5 | 21.5 ± 7.8 | 13.4 ± 6.2 |
| Pathotroph | 2.0 ± 0.2 | 0.1 ± 0.0 | 1.8 ± 0.4 | 0.4 ± 0.3 |
| **RAF - Fine roots** | | | | |
| Agaricales | 14.4 ± 6.3 | 6.8 ± 0.7 | 17.6 ± 8.9 | 22.5 ± 4.7 |
| Atheliales | 0.5 ± 0.5 | 0.6 ± 0.5 | 0.2 ± 0.1 | 0.6 ± 0.6 |
| Boletales | 12.3 ± 8.7 | 21.6 ± 13.1 | 11.1 ± 7.6 | 1.7 ± 1.0 |
| Cantharellales | 5.1 ± 4.2 | 4.0 ± 4.0 | 0.1 ± 0.1 | 4.3 ± 4.1 |
| Russulales | 35.9 ± 11.4 | 42.6 ± 20.7 | 49.9 ± 17.5 | 16.2 ± 7.9 |
| Thelephorales | 8.0 ± 5.9 | 4.3 ± 4.1 | 1.5 ± 0.7 | 16.3 ± 8.2 |
| Trechisporales | 5.8 ± 4.4 | 3.9 ± 2.9 | 1.1 ± 0.5 | 4.5 ± 1.3 |
| Eurotiales | 0.0 ± 0.0 | 0.1 ± 0.0 | 0.1 ± 0.0 | 0.0 ± 0.0 |
| Helotiales | 11.4 ± 1.3 | 9.1 ± 1.9 | 13.6 ± 5.0 | 11.8 ± 2.6 |
| Hypocreales | 0.4 ± 0.0 | 0.3 ± 0.1 | 0.3 ± 0.1 | 0.5 ± 0.1 |
| Mytilinidiales | 0.5 ± 0.2 | 0.7 ± 0.5 | 2.7 ± 2.3 | 0.7 ± 0.6 |
| Pezizales | 1.1 ± 0.3 | 3.0 ± 1.6 | 0.2 ± 0.1 | 1.0 ± 0.4 |
| Pleosporales | 0.1 ± 0.0 | 0.3 ± 0.2 | 0.0 ± 0.0 | 0.4 ± 0.2 |
| Sordariales | 1.0 ± 0.4 | 1.1 ± 0.5 | 0.6 ± 0.2 | 5.7 ± 3.0 |
| Mortirellales | 0.1 ± 0.0 | 0.2 ± 0.1 | 0.0 ± 0.0 | 0.2 ± 0.0 |
| Symbiotroph | 58.7 ± 3.4 | 84.5 ± 1.3 | 73.7 ± 5.9 | 87.3 ± 6.1 |
| Saprotroph | 20.3 ± 3.2 | 4.1 ± 1.7 | 10.9 ± 3.8 | 6.9 ± 3.0 |
| Pathotroph | 1.1 ± 0.1 | 0.1 ± 0.0 | 1.0 ± 0.2 | 0.2 ± 0.1 |

| **Fall** | | | | |
|---|---|---|---|---|
| **Organic layer** | **Con** | **N** | **P** | **P+N** |
| **SAF - Bulk soil** | | | | |
| Agaricales | 18.5 ± 2.0 | 19.3 ± 0.4 | 20.6 ± 0.8 | 18.6 ± 0.5 |
| Atheliales | 5.4 ± 0.4 | 6.3 ± 0.1 | 6.1 ± 0.3 | 6.1 ± 0.2 |
| Boletales | 4.4 ± 0.5 | 3.9 ± 0.1 | 4.3 ± 0.4 | 4.1 ± 0.0 |
| Cantharellales | 7.4 ± 6.0 | 1.5 ± 0.1 | 1.5 ± 0.0 | 1.4 ± 0.1 |
| Russulales | 11.7 ± 1.1 | 13.0 ± 0.5 | 11.8 ± 1.2 | 14.0 ± 1.0 |
| Thelephorales | 2.8 ± 1.1 | 2.1 ± 0.1 | 2.9 ± 0.6 | 2.2 ± 0.3 |
| Trechisporales | 0.6 ± 0.1 | 1.0 ± 0.1 | 0.9 ± 0.1 | 0.9 ± 0.1 |
| Eurotiales | 6.2 ± 0.5 | 7.0 ± 0.2 | 6.8 ± 0.3 | 7.0 ± 0.2 |
| Helotiales | 15.1 ± 0.5 | 16.1 ± 0.1 | 15.7 ± 0.2 | 16.1 ± 0.1 |
| Hypocreales | 4.5 ± 0.9 | 5.0 ± 0.8 | 3.9 ± 0.2 | 4.7 ± 0.4 |
| Mytilinidiales | 3.4 ± 0.2 | 3.9 ± 0.1 | 4.5 ± 0.5 | 3.8 ± 0.2 |
| Pezizales | 1.1 ± 0.1 | 1.2 ± 0.1 | 1.4 ± 0.1 | 1.2 ± 0.1 |
| Pleosporales | 3.3 ± 0.6 | 3.3 ± 0.3 | 3.4 ± 0.5 | 3.3 ± 0.1 |
| Sordariales | 2.6 ± 0.2 | 2.1 ± 0.1 | 2.1 ± 0.2 | 2.3 ± 0.2 |
| Mortirellales | 3.6 ± 0.7 | 3.7 ± 0.0 | 3.6 ± 0.0 | 3.0 ± 0.1 |
| Symbiotroph | 62.1 ± 3.5 | 46.7 ± 16.8 | 66.5 ± 3.2 | 36.9 ± 14.9 |
| Saprotroph | 21.0 ± 2.3 | 26.6 ± 15.4 | 19.8 ± 3.7 | 45.7 ± 7.7 |
| Pathotroph | 3.0 ± 0.8 | 0.5 ± 0.1 | 0.8 ± 0.2 | 2.0 ± 1.3 |
| **RAF - Fine roots** | | | | |
| Agaricales | 35.8 ± 1.8 | 34.7 ± 4.0 | 38.8 ± 1.8 | 39.2 ± 3.8 |
| Atheliales | 1.1 ± 0.1 | 1.1 ± 0.1 | 1.2 ± 0.1 | 1.3 ± 0.2 |
| Boletales | 8.7 ± 0.6 | 9.5 ± 0.6 | 11.6 ± 2.0 | 15.1 ± 6.0 |
| Cantharellales | 7.3 ± 4.2 | 3.2 ± 0.3 | 3.6 ± 0.2 | 3.1 ± 0.7 |
| Russulales | 7.0 ± 0.6 | 15.3 ± 10.1 | 5.9 ± 1.4 | 6.6 ± 1.7 |
| Thelephorales | 2.8 ± 1.4 | 2.1 ± 0.4 | 2.9 ± 0.5 | 2.0 ± 0.1 |
| Trechisporales | 0.7 ± 0.3 | 1.0 ± 0.3 | 0.5 ± 0.2 | 2.1 ± 1.6 |
| Eurotiales | 2.3 ± 0.2 | 2.6 ± 0.5 | 2.2 ± 0.1 | 2.2 ± 0.3 |
| Helotiales | 19.4 ± 1.7 | 15.5 ± 2.3 | 19.1 ± 2.1 | 14.5 ± 0.4 |
| Hypocreales | 0.4 ± 0.1 | 0.4 ± 0.1 | 0.2 ± 0.0 | 0.3 ± 0.0 |
| Mytilinidiales | 10.4 ± 0.2 | 10.9 ± 1.4 | 10.6 ± 0.3 | 10.1 ± 1.2 |
| Pezizales | 0.3 ± 0.0 | 0.3 ± 0.1 | 0.4 ± 0.0 | 0.5 ± 0.0 |
| Pleosporales | 0.3 ± 0.1 | 0.2 ± 0.0 | 0.2 ± 0.0 | 0.2 ± 0.0 |
| Sordariales | 0.2 ± 0.0 | 0.2 ± 0.0 | 0.2 ± 0.0 | 0.1 ± 0.0 |
| Mortirellales | 0.1 ± 0.0 | 0.1 ± 0.0 | 0.0 ± 0.0 | 0.1 ± 0.0 |
| Symbiotroph | 80.8 ± 1.8 | 73.1 ± 8.3 | 83 ± 1.6 | 68.1 ± 7.4 |
| Saprotroph | 10.7 ± 1.2 | 13.4 ± 7.6 | 10.1 ± 1.8 | 22.9 ± 3.8 |
| Pathotroph | 1.6 ± 0.4 | 0.3 ± 0.0 | 0.5 ± 0.1 | 1.3 ± 0.6 |

### Supplement Table S8 continued

**HP**

| Spring<br>Mineral layer | Con | N | P | P+N |
|---|---|---|---|---|
| **SAF - Bulk soil** | | | | |
| Agaricales | 23.4 ± 7.3 | 25.6 ± 3.4 | 17.6 ± 5.5 | 18.1 ± 3.5 |
| Atheliales | 0.6 ± 0.1 | 2.9 ± 1.6 | 0.7 ± 0.2 | 1.2 ± 0.4 |
| Boletales | 2.6 ± 0.9 | 1.1 ± 0.2 | 2.9 ± 1.3 | 3.0 ± 0.8 |
| Cantharellales | 10.6 ± 5.8 | 5.6 ± 1.6 | 4.3 ± 0.4 | 5.1 ± 0.9 |
| Russulales | 9.8 ± 3.6 | 14.9 ± 1.7 | 16.2 ± 6.5 | 17.5 ± 1.2 |
| Thelephorales | 2.0 ± 0.2 | 2.9 ± 0.4 | 4.0 ± 0.4 | 3.0 ± 0.5 |
| Trechisporales | 1.6 ± 0.3 | 1.6 ± 0.4 | 0.9 ± 0.0 | 1.3 ± 0.2 |
| Eurotiales | 0.5 ± 0.3 | 0.9 ± 0.3 | 0.5 ± 0.1 | 0.7 ± 0.2 |
| Helotiales | 11.3 ± 1.7 | 10.7 ± 0.7 | 13.2 ± 0.8 | 12.5 ± 1.1 |
| Hypocreales | 3.8 ± 0.7 | 3.4 ± 0.2 | 4.2 ± 0.7 | 4.4 ± 1.3 |
| Mytilinidiales | 1.5 ± 0.6 | 2.1 ± 0.9 | 2.5 ± 1.0 | 2.3 ± 0.7 |
| Pezizales | 1.7 ± 0.3 | 1.7 ± 0.4 | 5.3 ± 2.2 | 5.0 ± 1.6 |
| Pleosporales | 11.7 ± 6.5 | 4.4 ± 3.4 | 4.5 ± 3.3 | 1.2 ± 0.4 |
| Sordariales | 2.4 ± 0.6 | 2.7 ± 0.6 | 3.6 ± 1.7 | 2.4 ± 0.6 |
| Mortirellales | 6.6 ± 1.7 | 7.9 ± 0.5 | 8.3 ± 2.5 | 10.5 ± 0.9 |
| Symbiotroph | 55.7 ± 10.1 | 53.3 ± 8.6 | 58.7 ± 11.9 | 41.2 ± 8.9 |
| Saprotroph | 29.1 ± 9.5 | 23.6 ± 8.1 | 26.3 ± 5.4 | 43.0 ± 7.3 |
| Pathotroph | 1.2 ± 0.7 | 0.1 ± 0.1 | 1.1 ± 0.7 | 0.6 ± 0.6 |
| **RAF - Fine roots** | | | | |
| Agaricales | 22.8 ± 7.2 | 29.7 ± 10.6 | 12.1 ± 2.1 | 20.7 ± 5.0 |
| Atheliales | 2.3 ± 1.1 | 1.6 ± 0.8 | 0.7 ± 0.5 | 2.2 ± 1.1 |
| Boletales | 11.4 ± 6.5 | 5.2 ± 3.3 | 17.6 ± 7.4 | 12.8 ± 9.1 |
| Cantharellales | 13.7 ± 1.3 | 2.7 ± 1.3 | 1.2 ± 0.8 | 1.8 ± 0.5 |
| Russulales | 10.9 ± 4.4 | 17.0 ± 6.3 | 24.6 ± 6.5 | 16.5 ± 9.1 |
| Thelephorales | 1.0 ± 0.4 | 5.2 ± 5.1 | 0.6 ± 0.2 | 4.0 ± 1.4 |
| Trechisporales | 6.0 ± 2.7 | 15.9 ± 2.4 | 10.1 ± 4.4 | 13.6 ± 0.4 |
| Eurotiales | 0.3 ± 0.2 | 0.8 ± 0.6 | 0.1 ± 0.1 | 1.7 ± 1.7 |
| Helotiales | 22.1 ± 7.1 | 11.9 ± 1.8 | 21.4 ± 7.1 | 14.4 ± 4.6 |
| Hypocreales | 0.1 ± 0.0 | 0.1 ± 0.1 | 0.0 ± 0.0 | 0.1 ± 0.0 |
| Mytilinidiales | 5.8 ± 2.7 | 5.6 ± 0.8 | 6.1 ± 4.2 | 4.2 ± 1.3 |
| Pezizales | 1.9 ± 0.3 | 3.4 ± 2.7 | 5.0 ± 2.4 | 5.1 ± 3.5 |
| Pleosporales | 0.0 ± 0.0 | 0.0 ± 0.0 | 0.0 ± 0.0 | 0.1 ± 0.1 |
| Sordariales | 0.1 ± 0.0 | 0.4 ± 0.2 | 0.1 ± 0.0 | 0.6 ± 0.2 |
| Mortirellales | 0.1 ± 0.0 | 0.0 ± 0.0 | 0.0 ± 0.0 | 0.1 ± 0.0 |
| Symbiotroph | 77.7 ± 5.1 | 76.5 ± 4.3 | 78.9 ± 5.9 | 70.4 ± 4.4 |
| Saprotroph | 14.6 ± 4.7 | 11.9 ± 4.0 | 13.4 ± 2.7 | 21.6 ± 3.6 |
| Pathotroph | 0.7 ± 0.4 | 0.1 ± 0.1 | 0.7 ± 0.4 | 0.4 ± 0.3 |

| Fall<br>Mineral layer | Con | N | P | P+N |
|---|---|---|---|---|
| **SAF - Bulk soil** | | | | |
| Agaricales | 21.4 ± 0.7 | 20.5 ± 0.6 | 19.1 ± 0.5 | 20.5 ± 0.4 |
| Atheliales | 6.8 ± 0.3 | 6.9 ± 0.2 | 6.8 ± 0.1 | 6.6 ± 0.2 |
| Boletales | 4.7 ± 0.3 | 4.4 ± 0.2 | 4.2 ± 0.1 | 4.7 ± 0.1 |
| Cantharellales | 2.0 ± 0.3 | 1.7 ± 0.1 | 2.3 ± 0.7 | 1.7 ± 0.0 |
| Russulales | 12.3 ± 0.5 | 13.5 ± 0.8 | 12.8 ± 1.8 | 12.4 ± 0.3 |
| Thelephorales | 3.1 ± 0.7 | 2.4 ± 0.2 | 2.4 ± 0.3 | 2.3 ± 0.2 |
| Trechisporales | 1.0 ± 0.1 | 0.9 ± 0.0 | 0.8 ± 0.1 | 0.9 ± 0.2 |
| Eurotiales | 7.1 ± 0.2 | 7.3 ± 0.1 | 7.1 ± 0.1 | 7.1 ± 0.1 |
| Helotiales | 14.9 ± 0.3 | 15.2 ± 0.6 | 15.7 ± 0.8 | 15.5 ± 0.4 |
| Hypocreales | 3.4 ± 0.1 | 3.9 ± 0.4 | 3.6 ± 0.1 | 4.6 ± 1.0 |
| Mytilinidiales | 4.5 ± 0.1 | 4.1 ± 0.1 | 4.3 ± 0.5 | 4.0 ± 0.2 |
| Pezizales | 1.3 ± 0.1 | 1.1 ± 0.0 | 1.4 ± 0.2 | 1.2 ± 0.0 |
| Pleosporales | 2.9 ± 0.1 | 3.3 ± 0.1 | 3.2 ± 0.0 | 3.1 ± 0.0 |
| Sordariales | 2.0 ± 0.1 | 1.9 ± 0.0 | 2.3 ± 0.1 | 2.2 ± 0.2 |
| Mortirellales | 3.0 ± 0.2 | 3.1 ± 0.1 | 3.2 ± 0.4 | 3.3 ± 0.2 |
| Symbiotroph | 18.7 ± 7.1 | 86.3 ± 6.7 | 56.1 ± 7.1 | 77 ± 11.2 |
| Saprotroph | 37.2 ± 2.7 | 2.9 ± 1.4 | 17 ± 3.5 | 8.3 ± 1.4 |
| Pathotroph | 2.9 ± 0.3 | 0.2 ± 0.1 | 1.7 ± 0.4 | 0.1 ± 0.0 |
| **RAF - Fine roots** | | | | |
| Agaricales | 21.8 ± 15.7 | 7.1 ± 1.7 | 19.4 ± 2.0 | 9.3 ± 1.2 |
| Atheliales | 2.3 ± 2.2 | 0.2 ± 0.1 | 11.8 ± 9.2 | 1.0 ± 0.9 |
| Boletales | 5.3 ± 2.8 | 3.6 ± 1.7 | 13.7 ± 5.6 | 2.4 ± 0.3 |
| Cantharellales | 17.3 ± 17.3 | 0.0 ± 0.0 | 0.0 ± 0.0 | 0.1 ± 0.1 |
| Russulales | 18.2 ± 10.3 | 34.8 ± 17.1 | 10.4 ± 10.1 | 23.9 ± 13.2 |
| Thelephorales | 3.9 ± 2.5 | 0.8 ± 0.5 | 0.6 ± 0.2 | 0.9 ± 0.5 |
| Trechisporales | 8.9 ± 4.5 | 23.9 ± 8.0 | 5.8 ± 2.6 | 33.5 ± 17.2 |
| Eurotiales | 0.1 ± 0.1 | 1.1 ± 0.8 | 0.0 ± 0.0 | 0.3 ± 0.2 |
| Helotiales | 17.1 ± 3.5 | 21.6 ± 7.0 | 28.5 ± 8.6 | 21.6 ± 1.3 |
| Hypocreales | 0.2 ± 0.0 | 0.2 ± 0.0 | 0.2 ± 0.1 | 0.1 ± 0.1 |
| Mytilinidiales | 3.9 ± 2.3 | 3.4 ± 0.9 | 2.3 ± 1.1 | 1.8 ± 1.3 |
| Pezizales | 0.3 ± 0.1 | 2.0 ± 1.1 | 6.4 ± 4.9 | 0.6 ± 0.1 |
| Pleosporales | 0.1 ± 0.0 | 0.2 ± 0.0 | 0.1 ± 0.0 | 0.0 ± 0.0 |
| Sordariales | 0.1 ± 0.1 | 0.6 ± 0.3 | 0.1 ± 0.1 | 0.4 ± 0.2 |
| Mortirellales | 0.0 ± 0.0 | 0.1 ± 0.0 | 0.0 ± 0.0 | 0.0 ± 0.0 |
| Symbiotroph | 59.2 ± 3.5 | 93 ± 3.3 | 77.9 ± 3.5 | 88.2 ± 5.6 |
| Saprotroph | 18.6 ± 1.3 | 1.6 ± 0.6 | 8.6 ± 1.7 | 4.4 ± 0.7 |
| Pathotroph | 1.6 ± 0.2 | 0.1 ± 0.1 | 0.9 ± 0.2 | 0.1 ± 0.0 |

### Supplement Table S8 continued

**MP**

| Spring Organic layer | Con | N | P | P+N |
|---|---|---|---|---|
| **SAF - Bulk soil** | | | | |
| Agaricales | 13.3 ± 0.8 | 14.8 ± 3.6 | 14.6 ± 2.5 | 12.6 ± 0.6 |
| Atheliales | 3.1 ± 0.9 | 2.2 ± 0.2 | 3.0 ± 0.3 | 2.8 ± 0.6 |
| Boletales | 7.8 ± 1.2 | 6.5 ± 2.0 | 8.6 ± 0.6 | 5.9 ± 2.6 |
| Cantharellales | 0.7 ± 0.1 | 0.7 ± 0.2 | 0.8 ± 0.2 | 0.8 ± 0.2 |
| Russulales | 21.8 ± 1.5 | 29.0 ± 1.6 | 26.0 ± 5.6 | 24.1 ± 7.4 |
| Thelephorales | 7.8 ± 4.0 | 3.5 ± 1.4 | 3.8 ± 0.9 | 1.2 ± 0.2 |
| Trechisporales | 1.6 ± 1.1 | 0.8 ± 0.2 | 0.5 ± 0.1 | 0.4 ± 0.0 |
| Eurotiales | 1.7 ± 0.7 | 2.0 ± 0.5 | 1.5 ± 0.3 | 1.7 ± 0.6 |
| Helotiales | 21.8 ± 0.9 | 20.8 ± 3.3 | 18.5 ± 2.0 | 27.3 ± 4.4 |
| Hypocreales | 3.8 ± 0.8 | 4.2 ± 0.6 | 4.8 ± 2.2 | 4.3 ± 0.6 |
| Mytilinidiales | 1.6 ± 0.1 | 1.6 ± 0.1 | 1.6 ± 0.1 | 1.6 ± 0.1 |
| Pezizales | 0.3 ± 0.0 | 0.3 ± 0.0 | 0.4 ± 0.0 | 0.3 ± 0.0 |
| Pleosporales | 1.3 ± 0.1 | 1.3 ± 0.1 | 1.4 ± 0.2 | 1.4 ± 0.1 |
| Sordariales | 2.6 ± 0.2 | 3.8 ± 0.8 | 3.9 ± 1.3 | 3.6 ± 0.7 |
| Mortirellales | 3.8 ± 0.6 | 2.0 ± 0.3 | 3.9 ± 0.4 | 3.6 ± 0.8 |
| Symbiotroph | 57.2 ± 7.5 | 59.7 ± 11.3 | 70.1 ± 2.5 | 42.7 ± 18.9 |
| Saprotroph | 26.9 ± 7.5 | 28.7 ± 2.5 | 18.3 ± 1.3 | 43 ± 18.6 |
| Pathotroph | 2.5 ± 1.0 | 0.1 ± 0.0 | 0.7 ± 0.2 | 0.3 ± 0.1 |
| **RAF - Fine roots** | | | | |
| Agaricales | 38.6 ± 14.1 | 42.2 ± 1.3 | 26.8 ± 3.1 | 37.8 ± 4.1 |
| Atheliales | 1.1 ± 1.1 | 0.3 ± 0.2 | 0.1 ± 0.1 | 0.2 ± 0.2 |
| Boletales | 16.7 ± 5.7 | 8.6 ± 2.9 | 20.6 ± 4.4 | 11.0 ± 4.4 |
| Cantharellales | 0.3 ± 0.1 | 0.7 ± 0.6 | 0.2 ± 0.2 | 3.5 ± 2.8 |
| Russulales | 0.8 ± 0.4 | 21.5 ± 7.7 | 26.2 ± 11.5 | 23.2 ± 3.6 |
| Thelephorales | 18.6 ± 14.6 | 4.3 ± 4.3 | 11.7 ± 11.3 | 0.6 ± 0.5 |
| Trechisporales | 0.1 ± 0.0 | 0.3 ± 0.2 | 0.3 ± 0.2 | 0.5 ± 0.4 |
| Eurotiales | 0.3 ± 0.2 | 0.1 ± 0.0 | 0.0 ± 0.0 | 0.0 ± 0.0 |
| Helotiales | 21.9 ± 5.6 | 18.9 ± 4.6 | 11.6 ± 2.0 | 20.8 ± 2.3 |
| Hypocreales | 0.3 ± 0.1 | 1.1 ± 0.7 | 0.3 ± 0.1 | 0.7 ± 0.2 |
| Mytilinidiales | 0.1 ± 0.0 | 1.1 ± 0.8 | 0.5 ± 0.3 | 0.0 ± 0.0 |
| Pezizales | 0.0 ± 0.0 | 0.1 ± 0.1 | 0.0 ± 0.0 | 0.0 ± 0.0 |
| Pleosporales | 0.1 ± 0.0 | 0.0 ± 0.0 | 0.0 ± 0.0 | 0.1 ± 0.0 |
| Sordariales | 0.4 ± 0.2 | 0.2 ± 0.0 | 0.3 ± 0.1 | 0.4 ± 0.2 |
| Mortirellales | 0.1 ± 0.0 | 0.1 ± 0.0 | 0.3 ± 0.2 | 0.1 ± 0.0 |
| Symbiotroph | 78.4 ± 3.7 | 79.7 ± 5.6 | 84.8 ± 1.3 | 71.1 ± 9.3 |
| Saprotroph | 13.6 ± 3.7 | 14.5 ± 1.2 | 9.3 ± 0.7 | 21.6 ± 9.2 |
| Pathotroph | 1.3 ± 0.5 | 0.1 ± 0.0 | 0.4 ± 0.1 | 0.3 ± 0.1 |

| Fall Organic layer | Con | N | P | P+N |
|---|---|---|---|---|
| **SAF - Bulk soil** | | | | |
| Agaricales | 10.7 ± 1.2 | 16.4 ± 1.8 | 12.7 ± 2.7 | 11.7 ± 0.6 |
| Atheliales | 5.6 ± 1.4 | 2.3 ± 0.2 | 3.0 ± 0.0 | 3.7 ± 0.6 |
| Boletales | 2.7 ± 0.5 | 3.0 ± 0.9 | 3.7 ± 0.1 | 3.3 ± 0.4 |
| Cantharellales | 6.7 ± 5.7 | 0.8 ± 0.1 | 0.9 ± 0.1 | 1.8 ± 0.1 |
| Russulales | 15.6 ± 6.0 | 17.0 ± 9.2 | 21.8 ± 8.5 | 17.0 ± 6.2 |
| Thelephorales | 2.7 ± 0.3 | 1.1 ± 0.0 | 1.3 ± 0.1 | 1.9 ± 0.6 |
| Trechisporales | 0.3 ± 0.0 | 0.3 ± 0.0 | 0.6 ± 0.1 | 0.4 ± 0.0 |
| Eurotiales | 4.8 ± 0.4 | 3.8 ± 0.3 | 4.4 ± 0.1 | 5.3 ± 0.4 |
| Helotiales | 18.8 ± 1.2 | 19.0 ± 4.9 | 19.2 ± 2.8 | 22.7 ± 3.4 |
| Hypocreales | 10.3 ± 4.3 | 16.0 ± 5.2 | 8.2 ± 0.7 | 8.2 ± 1.1 |
| Mytilinidiales | 3.6 ± 0.6 | 2.6 ± 0.4 | 3.6 ± 0.5 | 3.6 ± 0.1 |
| Pezizales | 0.6 ± 0.1 | 0.5 ± 0.0 | 0.5 ± 0.1 | 0.6 ± 0.1 |
| Pleosporales | 2.4 ± 0.3 | 2.4 ± 0.3 | 2.5 ± 0.1 | 2.8 ± 0.2 |
| Sordariales | 2.2 ± 0.3 | 2.0 ± 0.2 | 1.7 ± 0.1 | 2.9 ± 0.8 |
| Mortirellales | 3.6 ± 0.2 | 5.3 ± 0.9 | 6.2 ± 1.9 | 5.4 ± 0.7 |
| Symbiotroph | 53.9 ± 5.6 | 55.9 ± 15.0 | 78.6 ± 3.3 | 34.8 ± 14.1 |
| Saprotroph | 30.0 ± 5.7 | 37.2 ± 13.1 | 14.0 ± 2.6 | 51.6 ± 12.6 |
| Pathotroph | 2.0 ± 0.7 | 0.2 ± 0.1 | 0.6 ± 0.1 | 0.6 ± 0.4 |
| **RAF - Fine roots** | | | | |
| Agaricales | 50.0 ± 11.0 | 32.1 ± 2.6 | 23.7 ± 4.4 | 25.7 ± 4.7 |
| Atheliales | 2.6 ± 1.3 | 0.1 ± 0.0 | 0.2 ± 0.1 | 0.1 ± 0.0 |
| Boletales | 3.3 ± 0.4 | 12.7 ± 4.5 | 25.6 ± 5.9 | 28.3 ± 10.7 |
| Cantharellales | 0.2 ± 0.1 | 4.0 ± 4.0 | 0.1 ± 0.0 | 0.1 ± 0.0 |
| Russulales | 11.7 ± 5.5 | 11.8 ± 8.6 | 20.1 ± 8.5 | 2.9 ± 0.5 |
| Thelephorales | 2.6 ± 1.0 | 2.3 ± 0.6 | 2.9 ± 1.6 | 2.7 ± 0.5 |
| Trechisporales | 0.7 ± 0.3 | 0.7 ± 0.4 | 1.9 ± 1.8 | 0.4 ± 0.2 |
| Eurotiales | 0.1 ± 0.0 | 0.1 ± 0.1 | 0.1 ± 0.1 | 0.1 ± 0.0 |
| Helotiales | 26.0 ± 6.1 | 31.1 ± 3.8 | 19.5 ± 7.6 | 37.3 ± 7.3 |
| Hypocreales | 0.5 ± 0.1 | 1.6 ± 0.4 | 2.0 ± 1.5 | 0.7 ± 0.3 |
| Mytilinidiales | 0.4 ± 0.1 | 0.4 ± 0.1 | 2.2 ± 1.9 | 0.4 ± 0.1 |
| Pezizales | 0.1 ± 0.0 | 0.3 ± 0.2 | 0.1 ± 0.0 | 0.1 ± 0.0 |
| Pleosporales | 0.0 ± 0.0 | 0.1 ± 0.0 | 0.1 ± 0.0 | 0.1 ± 0.0 |
| Sordariales | 0.2 ± 0.0 | 0.4 ± 0.1 | 0.2 ± 0.1 | 0.8 ± 0.1 |
| Mortirellales | 0.0 ± 0.0 | 0.2 ± 0.1 | 0.0 ± 0.0 | 0.1 ± 0.0 |
| Symbiotroph | 76.7 ± 2.8 | 77.8 ± 7.5 | 89 ± 1.7 | 67.2 ± 7.0 |
| Saprotroph | 15.1 ± 2.8 | 18.6 ± 6.5 | 7.2 ± 1.3 | 25.9 ± 6.2 |
| Pathotroph | 1.1 ± 0.4 | 0.2 ± 0.0 | 0.3 ± 0.0 | 0.5 ± 0.2 |

### Supplement Table S8 continued

**MP**

| **Spring**<br>**Mineral layer** | **Con** | **N** | **P** | **P+N** |
|---|---|---|---|---|
| **SAF - Bulk soil** | | | | |
| Agaricales | 25.1 ± 1.3 | 27.4 ± 0.9 | 29.5 ± 2.3 | 28.5 ± 1.3 |
| Atheliales | 5.5 ± 1.2 | 5.1 ± 0.9 | 4.4 ± 0.8 | 4.0 ± 0.3 |
| Boletales | 5.7 ± 0.6 | 4.0 ± 0.2 | 4.6 ± 0.6 | 5.2 ± 1.2 |
| Cantharellales | 5.1 ± 3.0 | 1.2 ± 0.1 | 2.7 ± 1.2 | 2.8 ± 0.7 |
| Russulales | 16.5 ± 0.4 | 20.3 ± 3.1 | 19.2 ± 4.3 | 16.1 ± 2.6 |
| Thelephorales | 2.1 ± 0.3 | 3.8 ± 1.4 | 2.2 ± 1.0 | 1.7 ± 0.4 |
| Trechisporales | 0.6 ± 0.1 | 0.4 ± 0.1 | 0.4 ± 0.0 | 0.6 ± 0.0 |
| Eurotiales | 1.8 ± 0.6 | 2.2 ± 0.6 | 2.1 ± 0.3 | 1.5 ± 0.2 |
| Helotiales | 14.9 ± 1.2 | 12.2 ± 0.8 | 13.1 ± 1.5 | 16.0 ± 0.7 |
| Hypocreales | 2.6 ± 0.4 | 4.1 ± 0.9 | 3.3 ± 0.7 | 2.9 ± 0.2 |
| Mytilinidiales | 3.0 ± 0.1 | 5.3 ± 0.9 | 4.5 ± 1.0 | 5.0 ± 0.7 |
| Pezizales | 0.9 ± 0.1 | 1.0 ± 0.2 | 0.9 ± 0.3 | 0.7 ± 0.0 |
| Pleosporales | 1.9 ± 0.1 | 1.7 ± 0.1 | 1.6 ± 0.1 | 2.0 ± 0.1 |
| Sordariales | 1.3 ± 0.2 | 1.0 ± 0.1 | 1.1 ± 0.1 | 1.2 ± 0.1 |
| Mortirellales | 5.4 ± 1.1 | 4.4 ± 0.5 | 4.4 ± 0.8 | 4.6 ± 1.1 |
| Symbiotroph | 20.7 ± 7.2 | 30.9 ± 15.1 | 56.8 ± 0.9 | 63.6 ± 6.2 |
| Saprotroph | 35.3 ± 6.9 | 19.9 ± 11.7 | 15.1 ± 0.7 | 14.9 ± 2.9 |
| Pathotroph | 3.3 ± 0.4 | 1.0 ± 0.5 | 1.7 ± 0.4 | 1.3 ± 1.3 |
| **RAF - Fine roots** | | | | |
| Agaricales | 44.9 ± 12.1 | 41.0 ± 5.2 | 48.6 ± 15.6 | 46.4 ± 7.6 |
| Atheliales | 1.1 ± 0.9 | 1.0 ± 0.4 | 0.6 ± 0.5 | 5.0 ± 2.8 |
| Boletales | 4.0 ± 3.6 | 3.0 ± 1.8 | 2.5 ± 1.4 | 1.5 ± 1.1 |
| Cantharellales | 1.1 ± 0.7 | 1.6 ± 1.1 | 2.7 ± 1.4 | 1.8 ± 1.2 |
| Russulales | 4.8 ± 2.3 | 20.1 ± 12.3 | 23.4 ± 17.2 | 5.5 ± 2.2 |
| Thelephorales | 0.8 ± 0.8 | 0.1 ± 0.0 | 0.2 ± 0.1 | 0.0 ± 0.0 |
| Trechisporales | 0.9 ± 0.9 | 0.2 ± 0.2 | 0.3 ± 0.1 | 0.0 ± 0.0 |
| Eurotiales | 1.1 ± 1.1 | 0.0 ± 0.0 | 0.5 ± 0.4 | 0.0 ± 0.0 |
| Helotiales | 33.8 ± 7.4 | 16.9 ± 2.3 | 14.0 ± 4.1 | 33.6 ± 10.7 |
| Hypocreales | 0.2 ± 0.1 | 0.4 ± 0.2 | 0.4 ± 0.3 | 0.1 ± 0.0 |
| Mytilinidiales | 4.1 ± 1.4 | 9.6 ± 7.2 | 2.7 ± 1.9 | 0.9 ± 0.4 |
| Pezizales | 0.2 ± 0.2 | 2.7 ± 2.6 | 2.9 ± 2.7 | 1.5 ± 1.3 |
| Pleosporales | 0.1 ± 0.1 | 0.5 ± 0.5 | 0.0 ± 0.0 | 0.3 ± 0.3 |
| Sordariales | 0.0 ± 0.0 | 0.1 ± 0.1 | 0.1 ± 0.1 | 0.1 ± 0.0 |
| Mortirellales | 0.1 ± 0.0 | 0.1 ± 0.0 | 0.0 ± 0.0 | 0.1 ± 0.1 |
| Symbiotroph | 60.3 ± 3.6 | 65.4 ± 7.5 | 78.2 ± 0.5 | 81.6 ± 3.1 |
| Saprotroph | 17.7 ± 3.4 | 10.0 ± 5.8 | 7.7 ± 0.3 | 7.6 ± 1.5 |
| Pathotroph | 1.7 ± 0.2 | 0.6 ± 0.3 | 0.9 ± 0.2 | 0.7 ± 0.6 |

| **Fall**<br>**Mineral layer** | **Con** | **N** | **P** | **P+N** |
|---|---|---|---|---|
| **SAF - Bulk soil** | | | | |
| Agaricales | 20.3 ± 5.6 | 19.6 ± 2.9 | 26.0 ± 6.3 | 28.3 ± 7.9 |
| Atheliales | 5.7 ± 2.6 | 6.5 ± 1.6 | 4.3 ± 1.0 | 7.1 ± 3.6 |
| Boletales | 2.7 ± 0.7 | 3.1 ± 0.2 | 2.9 ± 0.7 | 2.4 ± 0.3 |
| Cantharellales | 20.2 ± 11.2 | 1.0 ± 0.2 | 1.4 ± 0.5 | 1.8 ± 0.7 |
| Russulales | 8.7 ± 3.1 | 15.1 ± 6.8 | 12.9 ± 4.9 | 6.6 ± 0.2 |
| Thelephorales | 1.2 ± 0.2 | 1.3 ± 0.2 | 2.6 ± 1.0 | 1.6 ± 0.1 |
| Trechisporales | 0.3 ± 0.1 | 0.4 ± 0.1 | 0.4 ± 0.0 | 0.5 ± 0.1 |
| Eurotiales | 4.7 ± 0.9 | 5.0 ± 0.9 | 4.4 ± 0.4 | 5.1 ± 0.3 |
| Helotiales | 12.6 ± 2.7 | 13.8 ± 2.6 | 13.7 ± 1.8 | 17.5 ± 2.2 |
| Hypocreales | 2.8 ± 0.3 | 3.9 ± 1.0 | 3.9 ± 0.9 | 3.9 ± 0.2 |
| Mytilinidiales | 3.3 ± 0.5 | 7.1 ± 2.9 | 6.4 ± 2.3 | 5.2 ± 0.9 |
| Pezizales | 0.7 ± 0.2 | 0.6 ± 0.1 | 1.0 ± 0.1 | 1.8 ± 0.2 |
| Pleosporales | 2.1 ± 0.4 | 2.9 ± 0.6 | 2.4 ± 0.3 | 2.9 ± 0.2 |
| Sordariales | 1.0 ± 0.2 | 1.2 ± 0.2 | 1.3 ± 0.3 | 1.4 ± 0.1 |
| Mortirellales | 6.8 ± 3.1 | 11.3 ± 3.7 | 8.6 ± 1.1 | 5.4 ± 1.7 |
| Symbiotroph | 56 ± 6.4 | 83.2 ± 1.7 | 63.7 ± 1.8 | 50.3 ± 15.1 |
| Saprotroph | 27.7 ± 6.3 | 12.6 ± 2.7 | 20.8 ± 2.5 | 39.4 ± 16.1 |
| Pathotroph | 4.8 ± 1.4 | 0.6 ± 0.4 | 1.1 ± 0.1 | 1.3 ± 0.7 |
| **RAF - Fine roots** | | | | |
| Agaricales | 37.7 ± 1.0 | 43.5 ± 8.2 | 55.7 ± 5.1 | 51.6 ± 6.6 |
| Atheliales | 1.2 ± 0.7 | 0.8 ± 0.6 | 0.5 ± 0.2 | 2.7 ± 2.5 |
| Boletales | 2.9 ± 0.7 | 14.6 ± 9.3 | 2.7 ± 1.5 | 3.2 ± 0.3 |
| Cantharellales | 3.0 ± 1.6 | 8.9 ± 8.9 | 0.4 ± 0.2 | 0.4 ± 0.3 |
| Russulales | 13.6 ± 1.2 | 9.1 ± 3.9 | 10.3 ± 3.4 | 3.1 ± 0.8 |
| Thelephorales | 4.8 ± 0.3 | 2.2 ± 0.4 | 2.9 ± 0.2 | 3.4 ± 0.9 |
| Trechisporales | 0.2 ± 0.1 | 1.1 ± 1.1 | 0.5 ± 0.3 | 0.1 ± 0.0 |
| Eurotiales | 0.1 ± 0.1 | 0.0 ± 0.0 | 0.0 ± 0.0 | 0.1 ± 0.0 |
| Helotiales | 21.7 ± 2.4 | 16.3 ± 2.1 | 19.8 ± 4.5 | 29.6 ± 8.0 |
| Hypocreales | 0.2 ± 0.1 | 0.3 ± 0.1 | 0.3 ± 0.1 | 0.5 ± 0.3 |
| Mytilinidiales | 9.6 ± 8.5 | 2.5 ± 1.6 | 5.1 ± 2.5 | 0.7 ± 0.1 |
| Pezizales | 0.4 ± 0.1 | 0.2 ± 0.1 | 0.5 ± 0.3 | 0.2 ± 0.1 |
| Pleosporales | 0.1 ± 0.0 | 0.1 ± 0.0 | 0.1 ± 0.0 | 3.1 ± 3.0 |
| Sordariales | 0.1 ± 0.0 | 0.1 ± 0.0 | 0.1 ± 0.0 | 0.2 ± 0.1 |
| Mortirellales | 0.0 ± 0.0 | 0.0 ± 0.0 | 0.0 ± 0.0 | 0.0 ± 0.0 |
| Symbiotroph | 77.8 ± 3.2 | 91.4 ± 0.8 | 81.5 ± 1.0 | 74.9 ± 7.6 |
| Saprotroph | 14.0 ± 3.1 | 6.5 ± 1.3 | 10.7 ± 1.3 | 19.8 ± 8.0 |
| Pathotroph | 2.5 ± 0.7 | 0.3 ± 0.2 | 0.7 ± 0.0 | 0.8 ± 0.4 |

### Supplement Table S8 continued

**LP**

| Spring<br>Organic layer | Con | N | P | P+N |
|---|---|---|---|---|
| **SAF - Bulk soil** | | | | |
| Agaricales | 25.5 ± 1.2 | 19.2 ± 2.8 | 30.8 ± 5.4 | 33.7 ± 3.2 |
| Atheliales | 5.2 ± 2.6 | 0.9 ± 0.5 | 1.1 ± 0.4 | 2.6 ± 1.8 |
| Boletales | 2.1 ± 0.2 | 2.9 ± 0.7 | 3.9 ± 1.0 | 6.9 ± 1.8 |
| Cantharellales | 4.2 ± 1.6 | 3.5 ± 1.5 | 1.6 ± 0.6 | 1.6 ± 1.4 |
| Russulales | 9.9 ± 2.4 | 10.8 ± 2.0 | 11.7 ± 5.3 | 1.2 ± 0.3 |
| Thelephorales | 3.1 ± 1.1 | 2.0 ± 0.3 | 3.6 ± 1.3 | 5.7 ± 2.4 |
| Trechisporales | 8.0 ± 3.2 | 4.4 ± 1.6 | 0.3 ± 0.1 | 5.1 ± 3.4 |
| Eurotiales | 0.8 ± 0.5 | 5.6 ± 2.8 | 2.3 ± 1.9 | 1.1 ± 0.6 |
| Helotiales | 11.4 ± 1.3 | 14.6 ± 1.0 | 10.3 ± 1.2 | 9.5 ± 1.4 |
| Hypocreales | 3.8 ± 1.8 | 5.9 ± 0.8 | 3.6 ± 0.3 | 5.3 ± 0.9 |
| Mytilinidiales | 5.4 ± 2.2 | 1.3 ± 0.6 | 4.9 ± 1.5 | 3.3 ± 1.3 |
| Pezizales | 0.6 ± 0.2 | 0.7 ± 0.1 | 0.7 ± 0.6 | 4.1 ± 1.2 |
| Pleosporales | 0.8 ± 0.5 | 7.3 ± 6.8 | 5.7 ± 3.8 | 1.3 ± 0.4 |
| Sordariales | 4.0 ± 2.2 | 5.7 ± 1.9 | 5.1 ± 1.2 | 5.1 ± 0.6 |
| Mortirellales | 3.4 ± 0.6 | 2.5 ± 0.3 | 2.6 ± 0.6 | 5.5 ± 0.4 |
| Symbiotroph | 44.3 ± 12.4 | 49.1 ± 12.5 | 77.0 ± 7.8 | 34.6 ± 3.2 |
| Saprotroph | 31.9 ± 3.0 | 41.3 ± 15.3 | 13.1 ± 4.9 | 53.8 ± 4.1 |
| Pathotroph | 2.3 ± 0.6 | 0.2 ± 0.0 | 0.2 ± 0.0 | 0.3 ± 0.2 |
| **RAF - Fine roots** | | | | |
| Agaricales | 29.3 ± 9.1 | 46.4 ± 11.0 | 32.3 ± 13.3 | 44.0 ± 11.6 |
| Atheliales | 7.8 ± 6.5 | 3.6 ± 3.5 | 6.2 ± 5.8 | 2.6 ± 2.4 |
| Boletales | 2.5 ± 0.7 | 5.4 ± 4.4 | 19.7 ± 8.7 | 11.2 ± 6.4 |
| Cantharellales | 14.3 ± 11.5 | 2.7 ± 2.7 | 1.5 ± 1.4 | 0.5 ± 0.3 |
| Russulales | 5.6 ± 5.3 | 3.3 ± 3.1 | 11.6 ± 8.6 | 1.2 ± 0.7 |
| Thelephorales | 0.7 ± 0.6 | 1.9 ± 1.2 | 0.4 ± 0.2 | 7.8 ± 4.7 |
| Trechisporales | 1.6 ± 0.7 | 0.2 ± 0.1 | 0.1 ± 0.0 | 0.4 ± 0.2 |
| Eurotiales | 1.2 ± 0.6 | 3.4 ± 3.3 | 0.4 ± 0.2 | 2.5 ± 1.5 |
| Helotiales | 18.1 ± 4.9 | 17.8 ± 4.2 | 13.3 ± 3.0 | 12.3 ± 3.4 |
| Hypocreales | 0.4 ± 0.2 | 0.4 ± 0.1 | 0.4 ± 0.2 | 0.5 ± 0.2 |
| Mytilinidiales | 11.6 ± 6.1 | 6.6 ± 3.2 | 10.0 ± 8.1 | 12.8 ± 10.5 |
| Pezizales | 0.3 ± 0.1 | 3.2 ± 3.1 | 0.1 ± 0.0 | 0.8 ± 0.5 |
| Pleosporales | 0.1 ± 0.0 | 0.1 ± 0.0 | 0.0 ± 0.0 | 0.1 ± 0.0 |
| Sordariales | 0.3 ± 0.1 | 0.3 ± 0.2 | 0.3 ± 0.2 | 0.3 ± 0.1 |
| Mortirellales | 0.3 ± 0.1 | 0.2 ± 0.1 | 0.3 ± 0.2 | 0.2 ± 0.1 |
| Symbiotroph | 71.9 ± 6.2 | 74.5 ± 6.2 | 88.2 ± 3.9 | 67.2 ± 1.6 |
| Saprotroph | 16.1 ± 1.4 | 20.7 ± 7.6 | 6.8 ± 2.5 | 26.9 ± 2.0 |
| Pathotroph | 1.3 ± 0.3 | 0.1 ± 0.0 | 0.2 ± 0.0 | 0.3 ± 0.1 |

| Fall<br>Organic layer | Con | N | P | P+N |
|---|---|---|---|---|
| **SAF - Bulk soil** | | | | |
| Agaricales | 11.7 ± 6.8 | 14.3 ± 0.7 | 15.3 ± 0.6 | 18.1 ± 3.1 |
| Atheliales | 3.0 ± 0.1 | 4.7 ± 0.1 | 5.6 ± 0.3 | 4.8 ± 0.3 |
| Boletales | 1.3 ± 0.2 | 3.5 ± 0.6 | 4.1 ± 0.5 | 3.9 ± 0.3 |
| Cantharellales | 14.0 ± 2.9 | 1.4 ± 0.2 | 1.6 ± 0.3 | 1.5 ± 0.1 |
| Russulales | 2.0 ± 0.2 | 7.2 ± 0.2 | 10.4 ± 1.6 | 8.1 ± 0.4 |
| Thelephorales | 1.0 ± 0.4 | 2.6 ± 0.6 | 2.2 ± 0.1 | 4.2 ± 1.6 |
| Trechisporales | 7.7 ± 0.8 | 0.8 ± 0.2 | 0.6 ± 0.0 | 1.3 ± 0.8 |
| Eurotiales | 14.0 ± 1.0 | 14.7 ± 3.9 | 9.4 ± 0.8 | 8.6 ± 0.6 |
| Helotiales | 5.6 ± 1.1 | 16.1 ± 0.9 | 16.4 ± 0.8 | 16.3 ± 0.9 |
| Hypocreales | 5.3 ± 0.4 | 8.1 ± 1.8 | 6.8 ± 0.0 | 7.5 ± 0.6 |
| Mytilinidiales | 1.1 ± 0.2 | 4.5 ± 0.1 | 6.1 ± 1.0 | 4.8 ± 0.2 |
| Pezizales | 3.4 ± 0.4 | 1.0 ± 0.2 | 0.9 ± 0.0 | 1.0 ± 0.1 |
| Pleosporales | 1.9 ± 0.2 | 3.9 ± 0.4 | 3.7 ± 0.0 | 3.6 ± 0.1 |
| Sordariales | 2.7 ± 0.8 | 2.1 ± 0.2 | 2.5 ± 0.3 | 2.0 ± 0.2 |
| Mortirellales | 14.3 ± 0.7 | 2.6 ± 0.5 | 3.2 ± 0.4 | 3.1 ± 0.5 |
| Symbiotroph | 19.2 ± 0.9 | 83.3 ± 8.7 | 53.1 ± 7.1 | 79.6 ± 4.0 |
| Saprotroph | 44.4 ± 2.3 | 2.5 ± 0.6 | 18.4 ± 5.0 | 6.0 ± 2.3 |
| Pathotroph | 5.1 ± 2.2 | 0.1 ± 0.1 | 2.1 ± 0.1 | 0.1 ± 0.0 |
| **RAF - Fine roots** | | | | |
| Agaricales | 26.6 ± 19.4 | 34.2 ± 8.4 | 47.7 ± 21.5 | 30.0 ± 13.3 |
| Atheliales | 18.5 ± 17.8 | 2.3 ± 2.0 | 0.5 ± 0.3 | 0.2 ± 0.1 |
| Boletales | 1.7 ± 0.1 | 2.7 ± 1.0 | 4.3 ± 1.9 | 14.9 ± 11.0 |
| Cantharellales | 0.0 ± 0.0 | 0.0 ± 0.0 | 0.7 ± 0.7 | 0.3 ± 0.2 |
| Russulales | 13.7 ± 11.2 | 5.1 ± 2.7 | 0.5 ± 0.2 | 0.2 ± 0.1 |
| Thelephorales | 0.5 ± 0.2 | 0.1 ± 0.0 | 1.9 ± 1.7 | 0.4 ± 0.3 |
| Trechisporales | 1.1 ± 0.6 | 1.5 ± 1.0 | 0.5 ± 0.1 | 5.5 ± 4.8 |
| Eurotiales | 1.3 ± 0.7 | 9.2 ± 8.9 | 0.6 ± 0.3 | 0.6 ± 0.3 |
| Helotiales | 20.3 ± 4.7 | 25.3 ± 4.0 | 24.5 ± 5.2 | 38.0 ± 13.9 |
| Hypocreales | 0.7 ± 0.3 | 1.5 ± 0.2 | 0.5 ± 0.1 | 1.2 ± 0.4 |
| Mytilinidiales | 7.3 ± 4.3 | 2.2 ± 1.3 | 12.1 ± 11.6 | 2.5 ± 0.7 |
| Pezizales | 0.2 ± 0.2 | 0.1 ± 0.0 | 0.1 ± 0.0 | 0.5 ± 0.2 |
| Pleosporales | 0.1 ± 0.0 | 0.5 ± 0.4 | 0.1 ± 0.1 | 0.4 ± 0.1 |
| Sordariales | 0.1 ± 0.0 | 0.2 ± 0.0 | 0.3 ± 0.2 | 0.2 ± 0.1 |
| Mortirellales | 0.1 ± 0.1 | 0.1 ± 0.1 | 0.1 ± 0.0 | 0.2 ± 0.1 |
| Symbiotroph | 59.5 ± 0.4 | 91.5 ± 4.3 | 76.4 ± 3.5 | 89.6 ± 2.0 |
| Saprotroph | 22.2 ± 1.1 | 1.4 ± 0.3 | 9.3 ± 2.5 | 3.2 ± 1.1 |
| Pathotroph | 2.7 ± 1.1 | 0 ± 0.0 | 1.1 ± 0.1 | 0.1 ± 0.0 |

### Supplement Table S8 continued

**LP**

| **Spring**<br>**Mineral layer** | **Con** | **N** | **P** | **P+N** |
|---|---|---|---|---|
| **SAF - Bulk soil** | | | | |
| Agaricales | 15.7 ± 3.4 | 39.5 ± 12.9 | 25.1 ± 7.4 | 36.5 ± 12.8 |
| Atheliales | 6.7 ± 6.1 | 5.4 ± 2.3 | 6.1 ± 3.1 | 7.8 ± 6.6 |
| Boletales | 5.3 ± 1.8 | 10.4 ± 6.8 | 4.9 ± 1.5 | 6.8 ± 2.5 |
| Cantharellales | 5.5 ± 5.5 | 3.3 ± 2.6 | 2.4 ± 2.3 | 0.2 ± 0.1 |
| Russulales | 20.2 ± 11.2 | 6.9 ± 4.3 | 19.8 ± 10.5 | 15.7 ± 4.3 |
| Thelephorales | 1.0 ± 0.5 | 6.5 ± 4.5 | 1.6 ± 1.2 | 2.4 ± 1.3 |
| Trechisporales | 0.3 ± 0.2 | 0.1 ± 0.1 | 0.2 ± 0.1 | 0.8 ± 0.4 |
| Eurotiales | 8.8 ± 4.3 | 7.0 ± 4.6 | 9.6 ± 3.6 | 1.0 ± 0.3 |
| Helotiales | 15.0 ± 3.8 | 6.2 ± 1.9 | 10.5 ± 2.7 | 8.1 ± 0.5 |
| Hypocreales | 3.6 ± 1.2 | 2.1 ± 0.5 | 5.8 ± 2.0 | 3.1 ± 0.5 |
| Mytilinidiales | 1.8 ± 0.7 | 1.6 ± 0.7 | 2.9 ± 0.7 | 3.0 ± 0.9 |
| Pezizales | 1.3 ± 0.7 | 0.9 ± 0.7 | 0.3 ± 0.2 | 3.8 ± 3.4 |
| Pleosporales | 0.3 ± 0.2 | 0.1 ± 0.0 | 0.2 ± 0.1 | 0.2 ± 0.1 |
| Sordariales | 1.1 ± 0.3 | 1.2 ± 0.5 | 0.7 ± 0.3 | 0.4 ± 0.3 |
| Mortierellales | 5.2 ± 1.4 | 5.0 ± 1.4 | 3.7 ± 1.9 | 4.8 ± 2.1 |
| Symbiotroph | 60.9 ± 0.7 | 61.9 ± 13.8 | 68.9 ± 2.8 | 45.7 ± 11 |
| Saprotroph | 26.8 ± 1.1 | 33 ± 12.7 | 18.4 ± 2.2 | 40.9 ± 3.7 |
| Pathotroph | 1.8 ± 0.4 | 0.4 ± 0.0 | 0.6 ± 0.1 | 1.0 ± 0.2 |
| **RAF - Fine roots** | | | | |
| Agaricales | 55.3 ± 16.4 | 43.8 ± 4.2 | 43.6 ± 7.0 | 58.2 ± 14.8 |
| Atheliales | 3.0 ± 2.6 | 2.5 ± 1.2 | 2.1 ± 2.0 | 1.2 ± 0.9 |
| Boletales | 7.6 ± 6.2 | 10.7 ± 4.3 | 25.1 ± 13.1 | 11.4 ± 5.4 |
| Cantharellales | 0.9 ± 0.4 | 2.5 ± 1.8 | 0.7 ± 0.7 | 1.3 ± 0.2 |
| Russulales | 6.0 ± 5.5 | 5.3 ± 4.6 | 4.5 ± 2.7 | 0.8 ± 0.2 |
| Thelephorales | 0.5 ± 0.5 | 0.7 ± 0.6 | 0.0 ± 0.0 | 0.9 ± 0.5 |
| Trechisporales | 2.3 ± 2.2 | 2.3 ± 2.3 | 0.2 ± 0.1 | 1.8 ± 1.3 |
| Eurotiales | 0.7 ± 0.3 | 6.3 ± 5.1 | 2.5 ± 2.2 | 1.1 ± 0.7 |
| Helotiales | 16.1 ± 6.2 | 17.8 ± 2.6 | 13.0 ± 3.7 | 17.5 ± 6.1 |
| Hypocreales | 0.2 ± 0.1 | 0.2 ± 0.1 | 0.6 ± 0.3 | 0.1 ± 0.0 |
| Mytilinidiales | 3.7 ± 0.8 | 3.1 ± 1.9 | 5.2 ± 1.3 | 2.3 ± 1.1 |
| Pezizales | 0.4 ± 0.3 | 0.5 ± 0.5 | 0.1 ± 0.1 | 0.4 ± 0.3 |
| Pleosporales | 0.1 ± 0.0 | 0.6 ± 0.5 | 0.6 ± 0.6 | 0.5 ± 0.5 |
| Sordariales | 0.0 ± 0.0 | 0.1 ± 0.0 | 0.0 ± 0.0 | 0.4 ± 0.3 |
| Mortierellales | 0.0 ± 0.0 | 0.1 ± 0.0 | 0.0 ± 0.0 | 0.0 ± 0.0 |
| Symbiotroph | 80.3 ± 0.3 | 80.8 ± 6.9 | 84.2 ± 1.4 | 72.6 ± 5.5 |
| Saprotroph | 13.5 ± 0.6 | 16.6 ± 6.3 | 9.4 ± 1.1 | 20.5 ± 1.8 |
| Pathotroph | 1.0 ± 0.2 | 0.2 ± 0.0 | 0.4 ± 0.1 | 0.6 ± 0.1 |

| **Fall**<br>**Mineral layer** | **Con** | **N** | **P** | **P+N** |
|---|---|---|---|---|
| **SAF - Bulk soil** | | | | |
| Agaricales | 15.7 ± 0.8 | 16.8 ± 0.7 | 15.4 ± 0.4 | 15.8 ± 0.7 |
| Atheliales | 7.3 ± 1.7 | 6.0 ± 0.4 | 8.6 ± 2.8 | 5.8 ± 0.3 |
| Boletales | 4.0 ± 0.5 | 6.3 ± 2.6 | 4.1 ± 0.2 | 5.0 ± 0.6 |
| Cantharellales | 1.5 ± 0.1 | 2.4 ± 0.8 | 1.9 ± 0.2 | 1.6 ± 0.0 |
| Russulales | 12.1 ± 1.4 | 10.4 ± 1.2 | 9.2 ± 0.6 | 10.1 ± 0.9 |
| Thelephorales | 2.0 ± 0.2 | 2.3 ± 0.1 | 2.3 ± 0.1 | 3.5 ± 0.8 |
| Trechisporales | 0.6 ± 0.1 | 0.6 ± 0.1 | 0.6 ± 0.0 | 0.7 ± 0.1 |
| Eurotiales | 9.8 ± 1.2 | 9.3 ± 1.1 | 8.8 ± 0.3 | 8.8 ± 0.3 |
| Helotiales | 16.5 ± 0.4 | 15.9 ± 0.6 | 17.4 ± 0.7 | 17.0 ± 0.6 |
| Hypocreales | 4.7 ± 0.2 | 4.6 ± 0.4 | 4.9 ± 0.2 | 5.0 ± 0.3 |
| Mytilinidiales | 5.5 ± 0.4 | 5.3 ± 0.2 | 5.5 ± 0.5 | 5.9 ± 0.3 |
| Pezizales | 1.2 ± 0.0 | 1.3 ± 0.3 | 1.0 ± 0.1 | 1.1 ± 0.0 |
| Pleosporales | 4.1 ± 0.2 | 4.0 ± 0.3 | 4.2 ± 0.3 | 4.4 ± 0.2 |
| Sordariales | 1.8 ± 0.1 | 1.6 ± 0.1 | 1.7 ± 0.2 | 1.8 ± 0.1 |
| Mortierellales | 3.7 ± 1.7 | 3.1 ± 1.1 | 3.6 ± 0.9 | 2.9 ± 0.6 |
| Symbiotroph | 60.1 ± 8.3 | 70.8 ± 17.4 | 57.3 ± 13.4 | 53.5 ± 12.0 |
| Saprotroph | 22.2 ± 3.6 | 24.6 ± 18.6 | 33.7 ± 11.3 | 42.7 ± 12.6 |
| Pathotroph | 1.1 ± 0.2 | 0.1 ± 0.1 | 0.6 ± 0.3 | 0.3 ± 0.3 |
| **RAF - Fine roots** | | | | |
| Agaricales | 47.0 ± 20.7 | 40.3 ± 9.8 | 44.4 ± 4.2 | 35.2 ± 17.0 |
| Atheliales | 0.9 ± 0.7 | 0.6 ± 0.1 | 2.5 ± 2.1 | 0.8 ± 0.5 |
| Boletales | 3.1 ± 1.2 | 6.0 ± 4.3 | 11.4 ± 7.6 | 24.4 ± 16.7 |
| Cantharellales | 0.0 ± 0.0 | 0.0 ± 0.0 | 0.2 ± 0.1 | 0.1 ± 0.0 |
| Russulales | 14.1 ± 9.7 | 10.6 ± 9.5 | 0.3 ± 0.1 | 0.3 ± 0.1 |
| Thelephorales | 0.2 ± 0.1 | 0.1 ± 0.1 | 0.2 ± 0.1 | 0.7 ± 0.3 |
| Trechisporales | 1.6 ± 1.3 | 2.5 ± 2.3 | 4.1 ± 1.4 | 8.5 ± 5.2 |
| Eurotiales | 11.0 ± 10.9 | 6.8 ± 6.6 | 11.0 ± 10.8 | 5.8 ± 5.7 |
| Helotiales | 14.6 ± 3.1 | 24.7 ± 12.4 | 22.1 ± 4.3 | 19.2 ± 5.2 |
| Hypocreales | 0.3 ± 0.2 | 0.4 ± 0.2 | 0.3 ± 0.1 | 0.4 ± 0.1 |
| Mytilinidiales | 0.6 ± 0.2 | 1.0 ± 0.6 | 1.3 ± 0.2 | 1.1 ± 0.1 |
| Pezizales | 0.1 ± 0.0 | 0.6 ± 0.5 | 0.2 ± 0.1 | 0.4 ± 0.3 |
| Pleosporales | 0.1 ± 0.1 | 0.1 ± 0.0 | 0.1 ± 0.0 | 0.1 ± 0.0 |
| Sordariales | 0.0 ± 0.0 | 0.0 ± 0.0 | 0.0 ± 0.0 | 0.1 ± 0.0 |
| Mortierellales | 0.0 ± 0.0 | 0.0 ± 0.0 | 0.0 ± 0.0 | 0.0 ± 0.0 |
| Symbiotroph | 79.9 ± 4.1 | 85.2 ± 8.6 | 78.4 ± 6.7 | 76.7 ± 6.0 |
| Saprotroph | 11.2 ± 1.8 | 12.4 ± 9.2 | 17.0 ± 5.6 | 21.4 ± 6.3 |
| Pathotroph | 0.6 ± 0.1 | 0.1 ± 0.0 | 0.4 ± 0.2 | 0.2 ± 0.1 |

**Supplement Table S9: Statistical information on the effects of forest site, season and fertilization treatment on the relative abundance of fungal orders and trophic guilds in the organic layer and mineral topsoil for Fig. 5 and Table S8.** Soils and roots were collected in a P-rich, P-medium and P-poor forest in spring and fall 2018 and separated into organic layer and the mineral topsoil for analyses. Means can be found in the supplementary information (Table S8). Differences among means per forest type, season, treatment and the two-way interactions (Forest x season, Forest x treatment, Season x treatment) were tested by a linear mixed effect model using Poisson distribution with plot number as random effect. Calculations were performed separately for the organic layer and mineral topsoil. Tukey HSD was used as posthoc test. Bold letters indicate significant differences at $p \leq 0.05$.

| | Forest | | Season | | Treatment | | Forest x Season | | Forest x Treatment | | Season x Treatment | | N | | P | | P+N | |
|---|---|---|---|---|---|---|---|---|---|---|---|---|---|---|---|---|---|---|
| | F | *P* | F | *p* | F | *p* | F | *p* | F | *p* | F | *P* | F | *p* | F | *p* | F | *p* |
| **SAF - Bulk Soil** | | | | | | | | | | | | | | | | | | |
| **Organic Layer** | | | | | | | | | | | | | | | | | | |
| Agaricales | 13.8 | **<0.001** | 0.1 | 0.705 | 0.7 | 0.554 | 0.1 | 0.944 | 0.2 | 0.965 | 0.6 | 0.617 | 1.8 | 0.190 | 1.0 | 0.321 | 1.1 | 0.308 |
| Atheliales | 0.5 | 0.597 | 1.8 | 0.189 | 1.2 | 0.315 | 0.9 | 0.396 | 1.1 | 0.384 | 0.6 | 0.598 | 1.6 | 0.214 | 1.3 | 0.268 | 1.5 | 0.224 |
| Boletales | 4.1 | **0.022** | 1.4 | 0.235 | 1.9 | 0.141 | 2.5 | 0.093 | 1.7 | 0.140 | 1.2 | 0.326 | 0.2 | 0.874 | 3.2 | 0.083 | 2.1 | 0.153 |
| Cantharellales | 1.4 | 0.249 | 1.8 | 0.191 | 2.9 | **0.043** | 2.1 | 0.139 | 0.2 | 0.971 | 0.1 | 0.945 | 4.2 | **0.048** | 4.2 | **0.049** | 4.2 | **0.050** |
| Russulales | 23.0 | **<0.001** | 0.0 | 0.966 | 0.4 | 0.752 | 4.0 | **0.024** | 0.6 | 0.722 | 0.3 | 0.816 | 0.1 | 0.994 | 0.1 | 0.774 | 0.5 | 0.452 |
| Thelephorales | 1.7 | 0.192 | 6.8 | **0.012** | 0.2 | 0.914 | 0.9 | 0.410 | 2.2 | 0.057 | 0.4 | 0.769 | 0.3 | 0.566 | 0.0 | 0.839 | 0.0 | 0.943 |
| Trechisporales | 164.1 | **<0.001** | 11.9 | **0.001** | 0.6 | 0.598 | 14.8 | **<0.001** | 0.7 | 0.628 | 0.5 | 0.656 | 0.1 | 0.749 | 0.0 | 0.839 | 0.0 | 0.921 |
| Eurotiales | 0.9 | 0.428 | 17.8 | **<0.001** | 1.1 | 0.364 | 3.6 | **0.035** | 1.2 | 0.319 | 0.3 | 0.800 | 1.0 | 0.320 | 0.1 | 0.833 | 0.1 | 0.744 |
| Helotiales | 2.0 | 0.149 | 1.6 | 0.206 | 0.4 | 0.782 | 6.1 | **0.005** | 0.8 | 0.547 | 1.4 | 0.268 | 0.1 | 0.722 | 0.1 | 0.774 | 0.2 | 0.639 |
| Hypocreales | 37.4 | **<0.001** | 9.6 | **0.003** | 0.9 | 0.460 | 12.0 | **<0.001** | 0.9 | 0.531 | 1.9 | 0.135 | 0.4 | 0.551 | 0.0 | 0.988 | 0.1 | 0.791 |
| Mytilinidiales | 5.5 | **0.007** | 0.1 | 0.798 | 0.4 | 0.773 | 2.0 | 0.142 | 0.3 | 0.917 | 0.3 | 0.796 | 1.1 | 0.390 | 0.1 | 0.770 | 0.0 | 0.853 |
| Pezizales | 1.1 | 0.331 | 0.0 | 0.998 | 1.0 | 0.414 | 2.8 | 0.072 | 0.8 | 0.573 | 1.3 | 0.301 | 1.3 | 0.270 | 0.1 | 0.762 | 2.5 | 0.122 |
| Pleosporales | 11.9 | **<0.001** | 5.8 | **0.019** | 1.8 | 0.169 | 18.6 | **<0.001** | 1.3 | 0.291 | 1.7 | 0.178 | 0.3 | 0.585 | 0.0 | 0.906 | 0.9 | 0.352 |
| Sordariales | 0.7 | 0.523 | 3.8 | 0.058 | 0.9 | 0.458 | 4.2 | **0.021** | 1.0 | 0.468 | 0.9 | 0.430 | 1.5 | 0.234 | 0.5 | 0.483 | 0.9 | 0.352 |
| Mortirellales | 102.3 | **<0.001** | 13.4 | **0.001** | 2.1 | 0.119 | 5.7 | **0.006** | 0.9 | 0.528 | 0.9 | 0.434 | 0.0 | 0.893 | 1.1 | 0.309 | 0.3 | 0.585 |
| Symbiotroph | 0.3 | 0.746 | 0.3 | 0.601 | 6.4 | **0.001** | 0.5 | 0.637 | 2.2 | 0.057 | 0.1 | 0.966 | 6.6 | **0.015** | 14.2 | **<0.001** | 1.1 | 0.313 |
| Saprotroph | 1.4 | 0.246 | 0.3 | 0.560 | 6.2 | **0.001** | 5.4 | **0.007** | 1.6 | 0.173 | 0.1 | 0.982 | 2.3 | 0.142 | 22.9 | **<0.001** | 0.2 | 0.623 |
| Pathtroph | 1.2 | 0.310 | 4.2 | **0.045** | 19.9 | **<0.001** | 1.6 | 0.209 | 1.3 | 0.279 | 0.8 | 0.525 | 31.7 | **<0.001** | 22.9 | **<0.001** | 17.5 | **<0.001** |

**Table S9 continued**

| | **Forest** | | **Season** | | **Treatment** | | **Forest * Season** | | **Forest * Treatment** | | **Season * Treatment** | | **N** | | **P** | | **P+N** | |
|---|---|---|---|---|---|---|---|---|---|---|---|---|---|---|---|---|---|---|
| | **F** | ***P*** | **F** | ***p*** | **F** | ***p*** | **F** | ***p*** | **F** | ***p*** | **F** | ***p*** | **F** | ***p*** | **F** | ***p*** | **F** | ***p*** |
| **SAF - Bulk Soil** | | | | | | | | | | | | | | | | | | |
| **Mineral Layer** | | | | | | | | | | | | | | | | | | |
| Agaricales | 20.5 | **<0.001** | 1.8 | 0.186 | 0.1 | 0.941 | 0.3 | 0.730 | 0.4 | 0.880 | 0.2 | 0.910 | 0.2 | 0.664 | 0.2 | 0.693 | 0.0 | 0.896 |
| Atheliales | 13.9 | **<0.001** | 10.4 | **0.002** | 0.2 | 0.861 | 11.0 | **<0.001** | 0.4 | 0.872 | 0.4 | 0.748 | 0.1 | 0.787 | 0.1 | 0.752 | 0.1 | 0.804 |
| Boletales | 11.4 | **<0.001** | 0.3 | 0.724 | 1.1 | 0.378 | 0.4 | 0.703 | 1.5 | 0.184 | 2.4 | 0.080 | 0.1 | 0.778 | 1.7 | 0.189 | 1.6 | 0.220 |
| Cantharellales | 3.6 | **0.035** | 0.1 | 0.860 | 1.0 | 0.411 | 0.6 | 0.560 | 0.7 | 0.681 | 0.4 | 0.724 | 4.2 | **0.049** | 4.3 | **0.044** | 4.2 | **0.049** |
| Russulales | 5.6 | **0.006** | 11.0 | **0.001** | 2.4 | 0.079 | 7.6 | **0.001** | 0.8 | 0.540 | 0.8 | 0.516 | 0.0 | 0.868 | 0.3 | 0.568 | 0.8 | 0.364 |
| Thelephorales | 43.9 | **<0.001** | 5.7 | 0.021 | 0.7 | 0.566 | 0.5 | 0.591 | 1.1 | 0.392 | 1.3 | 0.298 | 0.6 | 0.463 | 0.6 | 0.429 | 0.3 | 0.576 |
| Trechisporales | 30.6 | **<0.001** | 2.5 | 0.117 | 0.6 | 0.637 | 3.6 | **0.033** | 0.4 | 0.847 | 0.9 | 0.457 | 0.4 | 0.517 | 0.0 | 0.874 | 0.0 | 0.914 |
| Eurotiales | 0.8 | 0.455 | 11.1 | **0.002** | 0.1 | 0.948 | 0.5 | 0.586 | 0.1 | 0.996 | 0.2 | 0.904 | 0.1 | 0.810 | 0.0 | 0.909 | 0.1 | 0.734 |
| Helotiales | 2.9 | 0.063 | 2.9 | 0.096 | 0.4 | 0.777 | 0.6 | 0.536 | 0.5 | 0.815 | 0.4 | 0.726 | 0.4 | 0.535 | 0.3 | 0.579 | 1.7 | 0.208 |
| Hypocreales | 97.1 | **<0.001** | 0.1 | 0.713 | 1.3 | 0.302 | 0.6 | 0.548 | 0.9 | 0.538 | 0.3 | 0.813 | 0.6 | 0.429 | 0.7 | 0.394 | 0.8 | 0.375 |
| Mytilinidiales | 13.2 | **<0.001** | 0.2 | 0.661 | 1.5 | 0.222 | 10.2 | **<0.001** | 1.0 | 0.407 | 0.1 | 0.957 | 1.6 | 0.222 | 3.2 | 0.085 | 0.9 | 0.349 |
| Pezizales | 25.2 | **<0.001** | 8.7 | **0.005** | 2.3 | 0.088 | 10.1 | **<0.001** | 2.1 | 0.074 | 1.0 | 0.411 | 0.0 | 0.857 | 1.2 | 0.287 | 2.1 | 0.156 |
| Pleosporales | 3.7 | **0.032** | 14.5 | **<0.001** | 1.1 | 0.393 | 8.3 | **0.001** | 1.4 | 0.246 | 0.8 | 0.506 | 0.8 | 0.520 | 0.1 | 0.760 | 0.6 | 0.456 |
| Sordariales | 8.1 | **0.001** | 9.6 | **0.003** | 0.1 | 0.721 | 3.0 | 0.059 | 1.1 | 0.377 | 1.1 | 0.352 | 3.8 | 0.060 | 0.2 | 0.901 | 0.7 | 0.408 |
| Mortirellales | 57.4 | **<0.001** | 1.8 | 0.189 | 0.4 | 0.765 | 22.4 | **<0.001** | 1.1 | 0.404 | 1.3 | 0.293 | 0.3 | 0.573 | 0.1 | 0.708 | 0.1 | 0.765 |
| Symbiotroph | 1.0 | 0.386 | 5.8 | **0.020** | 4.3 | **0.009** | 2.0 | 0.142 | 1.4 | 0.223 | 3.9 | **0.015** | 5.7 | **0.022** | 6.4 | **0.016** | 2.0 | 0.166 |
| Saprotroph | 1.9 | 0.155 | 0.9 | 0.358 | 3.1 | **0.034** | 2.8 | 0.070 | 1.2 | 0.320 | 1.0 | 0.401 | 4.2 | **0.047** | 5.4 | **0.026** | 0.1 | 0.724 |
| Pathtroph | 10.5 | **<0.001** | 0.1 | 0.820 | 18.7 | **<0.001** | 1.3 | 0.286 | 2.2 | 0.056 | 1.6 | 0.195 | 26.9 | **<0.001** | 10.5 | **0.003** | 14.4 | **<0.001** |

**Table S9 continued**

| | Forest | | Season | | Treatment | | Forest * Season | | Forest * Treatment | | Season * Treatment | | N | | P | | P+N | |
|---|---|---|---|---|---|---|---|---|---|---|---|---|---|---|---|---|---|---|
| | F | *P* | F | *p* | F | *p* | F | *p* | F | *p* | F | *p* | F | *p* | F | *p* | F | *P* |
| **RAF - Fine roots** | | | | | | | | | | | | | | | | | | |
| **Organic Layer** | | | | | | | | | | | | | | | | | | |
| Agaricales | 10.8 | **<0.001** | 0.6 | 0.439 | 1.7 | 0.177 | 19.9 | **<0.001** | 1.2 | 0.317 | 1.6 | 0.207 | 1.0 | 0.329 | 0.8 | 0.365 | 0.0 | 0.938 |
| Atheliales | 16.4 | **<0.001** | 7.5 | **0.009** | 3.1 | **0.035** | 3.9 | **0.027** | 1.4 | 0.249 | 0.4 | 0.743 | 2.7 | 0.109 | 2.3 | 0.135 | 2.0 | 0.169 |
| Boletales | 7.8 | **0.001** | 0.1 | 0.721 | 1.1 | 0.339 | 0.5 | 0.632 | 1.1 | 0.387 | 0.8 | 0.525 | 0.4 | 0.540 | 5.2 | **0.030** | 0.9 | 0.336 |
| Cantharellales | 5.0 | **0.011** | 0.7 | 0.396 | 4.0 | **0.013** | 4.1 | **0.022** | 2.2 | 0.054 | 0.7 | 0.565 | 0.0 | 0.858 | 2.8 | 0.105 | 3.2 | 0.085 |
| Russulales | 11.3 | **<0.001** | 1.8 | 0.190 | 1.0 | 0.419 | 2.5 | 0.091 | 1.4 | 0.245 | 0.8 | 0.505 | 0.7 | 0.395 | 1.6 | 0.213 | 0.8 | 0.365 |
| Thelephorales | 0.6 | 0.535 | 5.1 | **0.028** | 0.6 | 0.608 | 0.8 | 0.450 | 1.1 | 0.364 | 0.5 | 0.672 | 1.4 | 0.246 | 0.4 | 0.525 | 0.1 | 0.818 |
| Trechisporales | 363.0 | **<0.001** | 96.2 | **<0.001** | 0.1 | 0.930 | 94.9 | **<0.001** | 0.1 | 0.999 | 0.5 | 0.662 | 0.0 | 0.906 | 0.0 | 0.986 | 0.0 | 0.920 |
| Eurotiales | 74.0 | **<0.001** | 55.2 | **<0.001** | 0.2 | 0.878 | 26.6 | **<0.001** | 1.1 | 0.409 | 0.2 | 0.895 | 1.6 | 0.213 | 0.2 | 0.660 | 0.0 | 0.899 |
| Helotiales | 18.6 | **<0.001** | 22.3 | **<0.001** | 1.1 | 0.359 | 1.6 | 0.215 | 2.0 | 0.155 | 0.2 | 0.863 | 0.0 | 0.926 | 0.9 | 0.346 | 0.2 | 0.701 |
| Hypocreales | 132.9 | **<0.001** | 9.6 | **0.003** | 2.2 | 0.099 | 5.8 | **0.005** | 1.3 | 0.284 | 0.5 | 0.665 | 1.1 | 0.298 | 0.2 | 0.684 | 0.6 | 0.462 |
| Mytilinidiales | 56.5 | **<0.001** | 81.2 | **<0.001** | 2.3 | 0.089 | 45.0 | **<0.001** | 1.0 | 0.451 | 0.2 | 0.884 | 0.8 | 0.686 | 0.4 | 0.550 | 0.1 | 0.714 |
| Pezizales | 14.0 | **<0.001** | 6.1 | **0.017** | 0.8 | 0.521 | 3.1 | 0.052 | 0.8 | 0.575 | 0.7 | 0.542 | 1.1 | 0.300 | 0.8 | 0.368 | 2.3 | 0.138 |
| Pleosporales | 2.9 | 0.062 | 0.1 | 0.754 | 1.7 | 0.180 | 1.7 | 0.193 | 0.5 | 0.813 | 0.5 | 0.664 | 1.0 | 0.336 | 1.9 | 0.177 | 1.2 | 0.278 |
| Sordariales | 6.8 | **0.003** | 12.0 | **0.001** | 0.7 | 0.663 | 8.5 | **0.001** | 0.8 | 0.494 | 1.4 | 0.249 | 0.2 | 0.683 | 2.2 | 0.151 | 2.0 | 0.170 |
| Mortirellales | 259.4 | **<0.001** | 3.2 | 0.079 | 2.4 | 0.127 | 1.8 | 0.171 | 0.2 | 0.654 | 2.7 | 0.055 | 0.1 | 0.800 | 0.0 | 0.970 | 0.5 | 0.495 |
| Symbiotroph | 0.3 | 0.741 | 0.3 | 0.598 | 6.5 | **0.001** | 0.5 | 0.628 | 2.2 | 0.055 | 0.1 | 0.965 | 6.7 | **0.014** | 14.2 | **<0.001** | 1.0 | 0.315 |
| Saprotroph | 1.5 | 0.241 | 0.4 | 0.555 | 6.2 | **0.001** | 5.5 | **0.007** | 1.6 | 0.173 | 0.1 | 0.982 | 2.3 | 0.141 | 22.9 | **<0.001** | 0.2 | 0.615 |
| Pathtroph | 1.1 | 0.332 | 4.1 | **0.047** | 21.0 | **<0.001** | 1.4 | 0.249 | 1.5 | 0.192 | 0.7 | 0.552 | 34.4 | **<0.001** | 14.3 | **<0.001** | 16.2 | **<0.001** |

**Table S9 continued**

| | Forest | | Season | | Treatment | | Forest * Season | | Forest * Treatment | | Season * Treatment | | N | | P | | P+N | |
|---|---|---|---|---|---|---|---|---|---|---|---|---|---|---|---|---|---|---|
| | F | ***P*** | F | ***P*** | F | ***P*** | F | ***p*** | F | ***p*** | F | ***p*** | F | ***p*** | F | ***p*** | F | ***P*** |
| **RAF - Fine roots** | | | | | | | | | | | | | | | | | | |
| **Mineral Layer** | | | | | | | | | | | | | | | | | | |
| Agaricales | 30.8 | **<0.001** | 3.7 | 0.060 | 0.2 | 0.872 | 1.8 | 0.173 | 1.0 | 0.439 | 1.2 | 0.307 | 0.2 | 0.628 | 0.2 | 0.680 | 0.3 | 0.570 |
| Atheliales | 6.5 | **0.003** | 0.3 | 0.595 | 0.6 | 0.619 | 0.4 | 0.657 | 0.6 | 0.712 | 1.2 | 0.315 | 0.6 | 0.453 | 0.3 | 0.569 | 0.1 | 0.724 |
| Boletales | 2.6 | 0.085 | 0.8 | 0.380 | 0.4 | 0.730 | 2.1 | 0.128 | 1.8 | 0.126 | 0.5 | 0.679 | 0.1 | 0.905 | 4.3 | **0.045** | 0.4 | 0.514 |
| Cantharellales | 0.7 | 0.505 | 0.0 | 0.982 | 2.6 | 0.066 | 0.2 | 0.843 | 1.9 | 0.093 | 0.1 | 0.955 | 1.7 | 0.201 | 5.4 | **0.023** | 1.1 | 0.306 |
| Russulales | 2.2 | 0.122 | 0.2 | 0.658 | 0.3 | 0.824 | 0.9 | 0.410 | 0.8 | 0.579 | 1.2 | 0.312 | 0.0 | 0.872 | 0.4 | 0.405 | 0.0 | 0.872 |
| Thelephorales | 0.7 | 0.500 | 0.7 | 0.405 | 0.9 | 0.445 | 3.9 | **0.026** | 0.9 | 0.530 | 2.4 | 0.079 | 0.2 | 0.671 | 2.3 | 0.140 | 0.0 | 0.891 |
| Trechisporales | 1781.7 | **<0.001** | 51.0 | **<0.001** | 1.6 | 0.195 | 43.9 | **<0.001** | 1.5 | 0.212 | 2.2 | 0.105 | 0.0 | 0.856 | 0.0 | 0.854 | 0.0 | 0.962 |
| Eurotiales | 61.3 | **<0.001** | 1.2 | 0.286 | 0.9 | 0.437 | 2.6 | 0.088 | 1.3 | 0.258 | 0.8 | 0.493 | 0.0 | 0.838 | 0.0 | 0.915 | 0.9 | 0.351 |
| Helotiales | 8.2 | **0.001** | 2.2 | 0.140 | 2.2 | 0.100 | 2.3 | 0.114 | 2.1 | 0.068 | 1.9 | 0.144 | 0.8 | 0.519 | 0.7 | 0.409 | 0.0 | 0.935 |
| Hypocreales | 151.9 | **<0.001** | 4.4 | **0.041** | 1.6 | 0.205 | 3.2 | 0.051 | 1.6 | 0.161 | 1.4 | 0.244 | 0.1 | 0.742 | 0.3 | 0.597 | 0.0 | 0.996 |
| Mytilinidiales | 0.1 | 0.946 | 0.1 | 0.785 | 0.8 | 0.474 | 2.3 | 0.110 | 0.7 | 0.662 | 0.6 | 0.596 | 0.0 | 0.827 | 0.4 | 0.548 | 2.4 | 0.130 |
| Pezizales | 2.8 | 0.069 | 2.4 | 0.127 | 1.0 | 0.396 | 0.2 | 0.793 | 0.9 | 0.509 | 0.7 | 0.585 | 1.7 | 0.196 | 3.0 | 0.092 | 1.9 | 0.183 |
| Pleosporales | 0.9 | 0.407 | 0.1 | 0.819 | 1.2 | 0.321 | 2.1 | 0.128 | 1.0 | 0.440 | 0.5 | 0.700 | 0.9 | 0.358 | 2.0 | 0.166 | 1.6 | 0.212 |
| Sordariales | 2.4 | 0.104 | 1.8 | 0.190 | 1.4 | 0.254 | 1.1 | 0.326 | 1.3 | 0.289 | 0.6 | 0.623 | 0.4 | 0.545 | 1.5 | 0.227 | 1.2 | 0.280 |
| Mortirellales | 58.8 | **<0.001** | 1.7 | 0.192 | 0.1 | 0.951 | 1.5 | 0.225 | 0.1 | 0.995 | 0.2 | 0.907 | 0.0 | 0.855 | 0.1 | 0.727 | 0.1 | 0.813 |
| Symbiotroph | 1.0 | 0.383 | 5.8 | **0.020** | 4.3 | **0.009** | 2.0 | 0.150 | 1.4 | 0.226 | 3.8 | **0.015** | 5.8 | **0.021** | 6.3 | **0.017** | 1.9 | 0.169 |
| Saprotroph | 1.9 | 0.154 | 0.8 | 0.365 | 3.1 | **0.034** | 2.9 | 0.067 | 1.2 | 0.325 | 1.0 | 0.411 | 4.3 | **0.047** | 5.2 | **0.029** | 0.1 | 0.711 |
| Pathtroph | 9.4 | **<0.001** | 0.1 | 0.858 | 17.9 | **<0.001** | 1.1 | 0.349 | 2.1 | 0.073 | 1.6 | 0.202 | 28.6 | **<0.001** | 9.9 | **0.003** | 13.4 | **<0.001** |
